# Optimizing multifunctional fluorescent ligands for intracellular labeling

**DOI:** 10.1101/2022.07.02.498544

**Authors:** Pratik Kumar, Jason D. Vevea, Ariana N. Tkachuk, Kirby Campbell, Emma T. Watson, Anthony X. Ayala, Jonathan B. Grimm, Edwin R. Chapman, David J. Solecki, Luke D. Lavis

## Abstract

Enzyme-based self-labeling tags enable covalent attachment of synthetic molecules to proteins inside living cells. A frontier of this field is designing multifunctional ligands that contain both fluorophores and affinity tags or pharmacological agents and can still efficiently enter cells. Self-labeling tag ligands with short linkers can enter cells readily but often show less activity due to steric issues; ligands with long linkers can be more potent but show lower cell permeability. Here, we overcome this tug-of-war between efficacy and cell-permeability by devising a rational strategy for making cell permeable multifunctional ligands for labeling HaloTag fusions. We found that the lactone–zwitterion equilibrium sconstant (*K*_L–Z_) of rhodamines inversely correlates with their distribution coefficients (log*D*_7.4_), suggesting that ligands based on dyes exhibiting low *K*_L–Z_ and high log*D*_7.4_ values, such as Si-rhodamines, would efficiently enter cells. We designed cell-permeable multifunctional HaloTag ligands with a biotin moiety to purify mitochondria or a JQ1 appendage to translocate BRD4 from euchromatin to the nucleolus or heterochromatin. We discovered that translocation of BRD4 to constitutive heterochromatin in cells expressing HaloTag–HP1a fusion proteins can lead to apparent increases in transcriptional activity. These new reagents enable affinity capture and translocation of intracellular proteins in living cells and the use of Si-rhodamines and other low *K*_L–Z_/high log*D*_7.4_ dye scaffolds will facilitate the design of new multifunctional chemical tools for biology.

**SIGNIFICANCE STATEMENT:** Understanding cellular processes requires tools to measure and manipulate proteins in living cells. Self- labeling tags, such as the HaloTag and SNAP-tag, enable modification of cellular proteins with synthetic molecules. Creating ligands for these systems that have more than one chemical motif remains challenging, however, due to competing demands between cell permeability and functionality. We discovered that multifunctional ligands based on Si-rhodamines efficiently entered cells and enabled affinity purification of mitochondria or translocation of nuclear proteins; the performance of these molecules could be verified by fluorescence microscopy. These compounds should be useful for a variety of biological experiments and our general framework will allow the design of other multifunctional ligands to study living systems.

## Introduction

Research at the interface of organic chemistry and protein biochemistry has generated powerful tools to visualize, purify, and manipulate cellular components.^1–3^ Introducing synthetic moieties into cells can be accomplished in different ways. Metabolic incorporation^4, 5^ utilizes endogenous enzymes to install nonnative moieties into cells while genetic code expansion,^6, 7^ engineered ligases,^8–12^ or self-labeling tags^13–15^ utilize exogenously expressed enzymes. In particular, engineering of enzyme–substrate interactions has produced self-labeling tags such as HaloTag^13^ and SNAP-tag^14^, which have found broad use in modern biology. These protein tags react specifically and irreversibly with a ligand motif that can be appended to a variety of functionalities, including fluorescent dyes, affinity tags, and pharmacological agents (**Fig. 1A**).^16–21^

**Figure 1.**
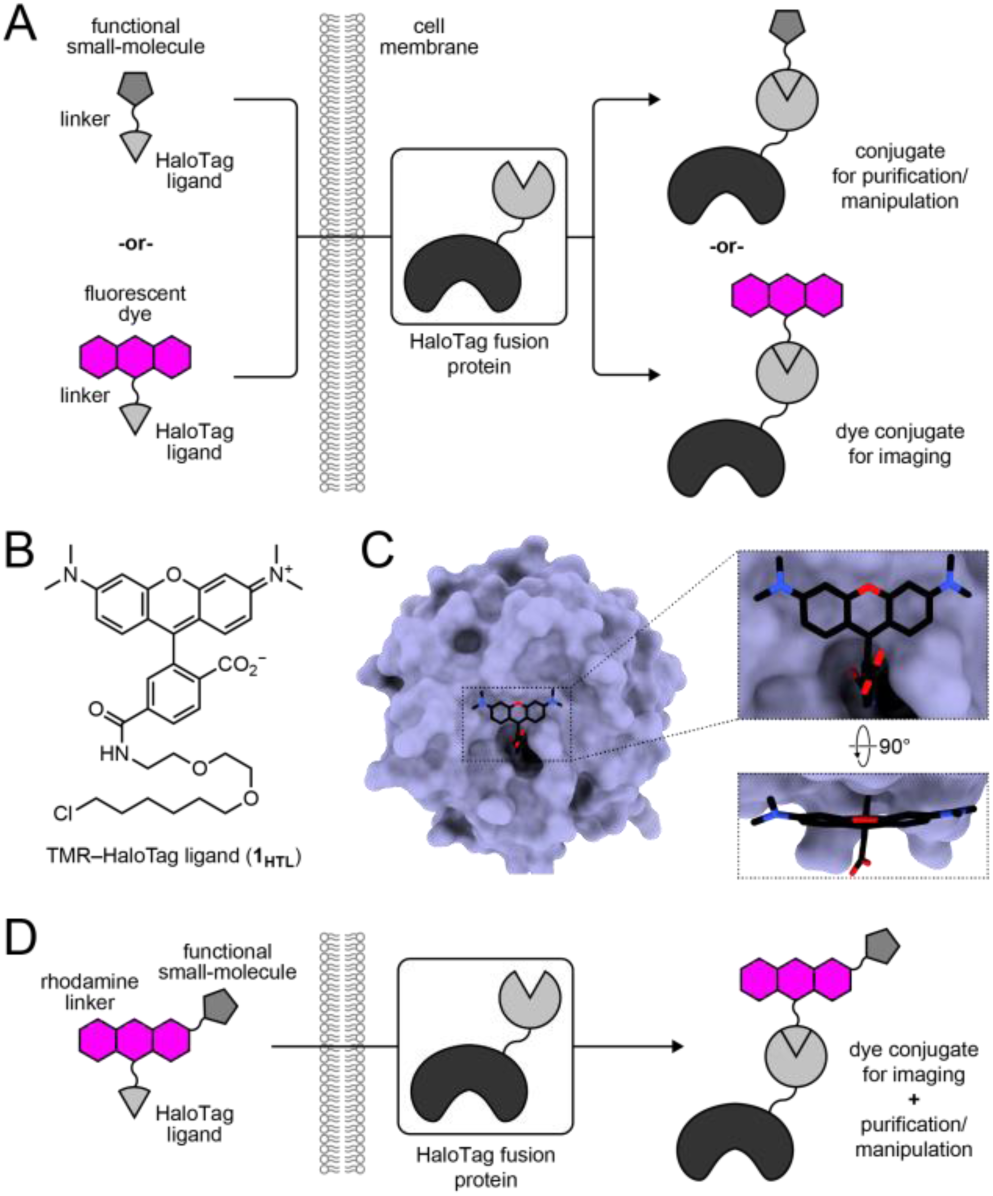
Rhodamine-based multifunctional ligands for self-labeling tags. (A) Schematic illustrating the modularity of self-labeling tag systems for singular function. (B) Chemical structure of **1_HTL_**. (C) Crystal structure of **1_HTL_** covalently bound to the HaloTag (PDB:6U32) with zoom-in of the dye–protein interface. (D) Schematic showing the concept of multifunctional ligands.

Despite the prospective flexibility of self-labeling tag systems, the major application of this technology has been to label proteins with small-molecule fluorophores. These systems were originally envisioned as complements to fluorescent proteins and the original protein engineering efforts that produced these tags focused on dye labeling. For example, the directed evolution campaign to generate the HaloTag used tetramethylrhodamine (TMR) linked to 1-chlorohexane ligand at the 6-position via a short polyethylene glycol (PEG) unit.^13^ Reaction of the TMR-HaloTag ligand (**1_HTL_**, **Fig. 1B**) with the HaloTag protein positions TMR close to the HaloTag surface (**Fig. 1C**; PDB:6U32).^22^ This general design of the PEG_2_- chlorohexane moiety (*i.e.*, the HaloTag ligand) attached to 6-carboxyrhodamines has generated a portfolio of spectrally distinct fluorescent HaloTag ligands that can function in cells, tissues, and animals.^16, 22–26^

The intimate dye–protein contact exemplified by the **1_HTL_**–HaloTag complex (**Fig. 1C**) can be advantageous for fluorescent ligands. The tight association can shift some dyes from nonfluorescent forms in water to fluorescent conjugates, yielding high-contrast fluorogenic systems.^26–30^ Association with the HaloTag can also improve brightness and photostability.^31^ Nevertheless, the relatively short PEG_2_-chloroalkane substrate motif can be problematic when the rhodamine moiety is replaced with other functionalities such as affinity tags or pharmacological agents. Given this issue, an emerging idea in the field is retain the dye moiety and append it with motifs for purification or perturbation, creating multifunctional ligands (**Fig. 1D**). This idea is predicated on several factors. First, as mentioned above, the directed evolution of the HaloTag utilized rhodamine ligand **1_HTL_**;^13^ rhodamine-containing HaloTag ligands show rapid labeling kinetics (∼10^7^ M^−1^s^−1^)^21, 32^ even when functionality is added to the dye moiety.^21^ Second, improvements in dye chemistry allows the straightforward synthesis of functionalized fluorophores where different moieties are attached to the dye for labeling^33^ or sensing^34–39^ applications. Third, the incorporation of a fluorophore enables visualizing the subcellular distribution of ligands inside cells, allowing verification of their performance using fluorescence imaging.

Multifunctional fluorescent ligands combine different functionalities—ligand, dye, and affinity tag or pharmacological agent—into a single compound. This yields relatively large molecules that can show variable cell-permeability. We set out to elucidate general design principles for multifunctional fluorescent ligands, exploring how the chemical properties of the parent rhodamine structure^27–30, 40, 41^ affects the permeability of multifunctional ligands built from that dye. We discovered that the lactone–zwitterion equilibrium constant (*K*_L–Z_) of rhodamine dyes is correlated with their octanol–water distribution coefficient at physiological pH (log*D*_7.4_), a common metric used in medicinal chemistry.^42^ Although addition of functionality onto the dye structure affects *K*_L–Z_ and log*D*_7.4_, the changes are relatively minor; the parent dye exerts a strong effect on the resulting multifunctional ligand properties. Dyes with relatively low *K*_L–Z_ values and high log*D*_7.4_, such as the Si-rhodamine-based Janelia Fluor 646 (JF_646_) and JF_635_, were particularly useful scaffolds for constructing cell-permeable multifunctional ligands. We prepared fluorogenic HaloTag ligands bearing the polar affinity tag biotin or the lipophilic pharmacological agent JQ1. These compounds are useful for advanced cell biological experiments, and the general concepts described below should aid the design of new multifunctional chemical tools for biology.

## Results and Discussion

### *K*_L–Z_ and log*D*_7.4_ are correlated

To optimize rhodamine-based multifunctional fluorescent ligands we first considered a key property of this dye class: the equilibrium between the nonfluorescent and lipophilic lactone (L) form, and the fluorescent and polar zwitterion form (Z; **Fig. 2A**). In previous work we showed the lactone–zwitterion equilibrium constant (*K*_L–Z_) exerts a strong effect on the performance of rhodamine ligands in biological systems.^40^ Dyes with high *K*_L–Z_ values, such as Janelia Fluor 549 (JF_549_, **2**; *K*_L–Z_ = 3.5), predominantly exist in the zwitterion form to yield bright and environmentally insensitive ligands. JF_549_ is structurally similar to TMR, the base dye of **1_HTL_**, but contains azetidines instead of *N,N*-dimethylamino groups, which increase brightness and photostability.^43^ The *K*_L–Z_ can be tuned lower by replacing the xanthene oxygen in **2** with a *gem*-dimethylcarbon (*e.g.*, JF_608_, **3**; *K*_L–Z_ = 0.091) or by installing 3,3-difluoroazetidine auxochromes, as in JF_525_ (**4**; *K*_L–Z_ = 0.068). Carborhodamine **3** exhibits a 60-nm bathochromic shift in absorption maximum (*λ*_abs_) and fluorescence emission maximum (*λ*_em_) relative to JF_549_ (**2**), whereas the 3,3-difluoroazetidine groups in **4** cause a 24-nm hypsochromic shift in *λ*_abs_ and *λ*_em_.^21^ Ligands based on JF_608_ or JF_525_ exhibit improved membrane permeability and bioavailability in animals.^27^ Dyes with even lower *K*_L–Z_ values can be accessed by incorporating a *gem*-dimethylsilicon substituent (*e.g.*, JF_646_, **5**; *K*_L–Z_ = 0.0014), which also causes a 100-nm bathochromic shift in *λ*_abs_ or *λ*_em_, or by combining C(CH_3_)_2_- or Si(CH_3_)_2_-substitutions with fluorinated azetidine auxochromes, as in JF_635_ (**6**; *K*_L–Z_ = 0.0001) and JF_585_ (**7**; *K*_L–Z_ = 0.001).^44–46^ Compounds based on **5**–**7** predominantly exist in the lactone form in aqueous media but can shift to the fluorescent zwitterion upon binding their cognate biomolecular target.^27^

**Figure 2.**
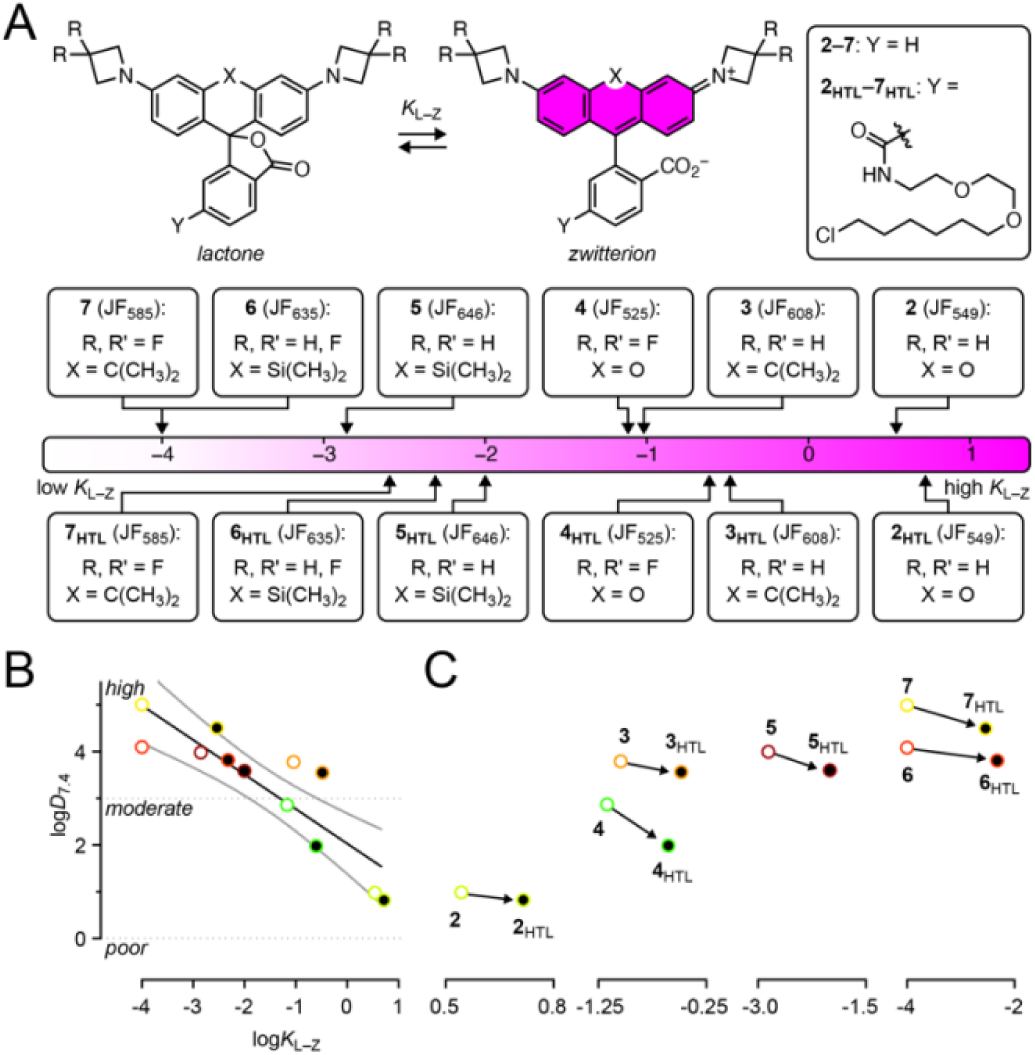
Relationship between K_L–Z_ and logD_7.4_ for rhodamines. (A) Dynamic equilibrium between the nonfluorescent lactone (L) and fluorescent zwitterion (Z) along with the lactone–zwitterion equilibrium constant (K_L–Z_) for rhodamine-based Janelia Fluor (JF) dyes **2**–**7** and their HaloTag ligands (**2_HTL_**–**7_HTL_**). K_L– Z_ values are in the log scale. (B) Correlation between logD_7.4_ versus log K_L–Z_ for **2**–**7** and **2_HTL_**–**7_HTL_**. Plot shows linear fit and 95% confidence interval. Dashed lines denote expected cellular permeability based on logD_7.4_. (C) Individual pairs of dye and their HaloTag ligands.

We then investigated how incorporation of the HaloTag ligand affects *K*_L–Z_. As for the free dyes **2**–**7**, we measured the *K*_L–Z_ values of rhodamine HaloTag ligands (**2_HTL_**–**7_HTL_**; **Fig. 2B**) in 1:1 (v/v) dioxane:water mixtures.^27, 40^ Incorporating a HaloTag ligand on the rhodamine structure consistently increases *K*_L–Z_ across the compound series (**Fig. 2C,D**), likely due to the electron-withdrawing carboxamide on the pendant phenyl ring that decreases the nucleophilicity of the *para*-carboxylate group that is responsible for lactone formation. We then investigated whether the *K*_L–Z_ was correlated to the distribution coefficients at pH 7.4 (log*D*_7.4_). log*D*_7.4_ values are employed in medicinal chemistry to rationalize the permeability of small-

molecule pharmacological agents,^42, 47^ but this parameter has not applied to self-labeling tag ligands. We measured the log*D*_7.4_ values for **2**–**7** and **2_HTL_**–**7_HTL_** using equilibrated mixtures of octanol and phosphate-buffered saline, pH 7.4 (octanol:PBS).^48^ Although the relationship between log*D*_7.4_ and cell-permeability is complex, cellular entry is optimal when log*D*_7.4_ is greater than 1 but less than 3–5 (see: Lipinkski);^49–51^ the permeability of higher molecular weight molecules benefits from log*D*_7.4_ values at the higher end of this range.^47^ Based on this general rule, we hypothesized that functional ligands based on JF_608_ (**3**; log*D*_7.4_ = 3.78), JF_646_ (**5**; log*D*_7.4_ = 3.98), and JF_635_ (**6**; log*D*_7.4_ = 4.10) could serve as effective scaffolds for cell-permeable compounds. We also surmised that the utility of JF_549_ (**2**) could be limited due to its lower log*D*_7.4_

= 0.98, which is at the edge the optimal range. Although JF_525_ (**4**) and JF_585_ (**7**) also exhibit potentially useful log*D*_7.4_ values—2.86 and 5.01, respectively—the 3,3-difluoroazetidine moiety in these compounds prevents attachment of functional small molecules; these dyes were not investigated further as scaffolds for multifunctional ligands. Overall, we found an inverse correlation between *K*_L–Z_ and log*D*_7.4_ (**Fig. 2C**), consistent with previous observations that dyes with low *K*_L–Z_ values exhibit high cell permeability and *vice versa*.^40^

### Synthesis and properties of biotin-JF-HaloTag ligands

We then prepared a series of multifunctional ligands to test how the parent dye scaffold affects the properties of the resulting compound. We initially explored compounds containing biotin, a molecule that is widely used in modern biotechnology due to its small size and tight association with the protein avidin.^52^ We coupled the 3-carboxyazetidine-containing compounds **8**–**11** with the commercially available biotin-PEG_2_-NH_2_ (**12**) to synthesize the biotin-JF-HaloTag ligand compounds **13_HTL_**–**16_HTL_** (**Scheme 1**, **SI Appendix**). The 3ʺ-carboxy-JF_549_-HaloTag ligand (**8**) and 3ʺ-carboxy-JF_646_-HaloTag ligand (**10**) starting materials were prepared as previously described.^35^ The JF_608_ and JF_635_ derivatives **9** and **11** were synthesized using an analogous sequence in five steps (**Scheme S1 and S2**, **SI Appendix**).

**Scheme 1.**
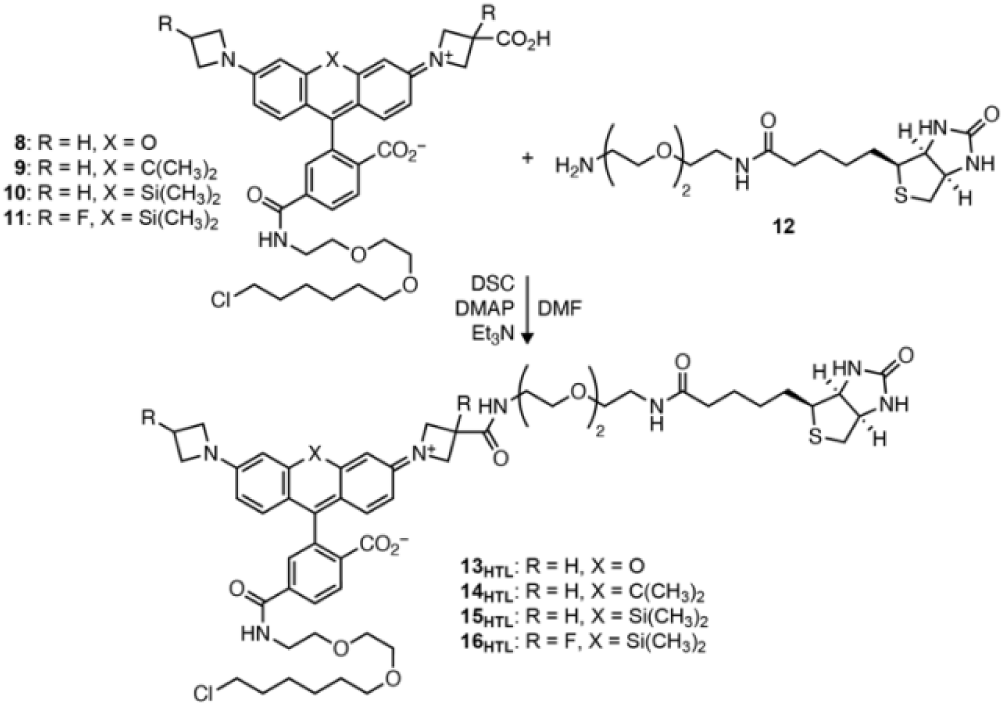
Synthesis of biotin-JF-HaloTag ligands **13_HTL_**–**16_HTL_** from 3ʺ-carboxyazetidine dyes **8**–**11** and biotin amine **12**.wsws

The spectral properties of the biotin-JF-HaloTag ligands **13_HTL_**–**16_HTL_**and their HaloTag conjugates were similar to the parent dyes **2**–**6** and HaloTag ligands **2_HTL_**–**6_HTL_** (**Fig. 3A,B**).^27, 43^ JF_549_ (**2**) shows *λ*_abs_/*λ*_em_ = 549 nm/571 nm, a large extinction coefficient (*ε* = 101,000 M^−1^cm^−1^), and high fluorescence quantum yield (*Φ*_f_ = 0.88). The electron-withdrawing carboxamide group on the pendant phenyl ring in **2_HTL_** causes a slight bathochromic shift (*λ*_abs_/*λ*_em_ = 554 nm/577 nm) along with a decreased *ε* = 70,000 M^−1^cm^−1^ and *Φ*_f_ =0.82. Binding of **2_HTL_** to the HaloTag protein elicits an additional bathochromic shift with largely unchanged brightness (*ε* = 83,000 M^−1^cm^−1^, *Φ*_f_ = 0.81). The incorporation of the biotin moiety in **13_HTL_** causes a slight hypsochromic shift compared to **2_HTL_** with *λ*_abs_/*λ*_em_ = 552 nm/575 nm; this is due to the electron-withdrawing carboxamide on the azetidine.^27^ **13_HTL_** shows *ε* = 92,000 M^−1^cm^−1^ and high *Φ*_f_ = 0.90; the spectral properties of the HaloTag conjugate of **13_HTL_** (**13–HT**) are quite similar to the free ligand (*λ*_abs_/*λ*_em_ = 552 nm/573 nm, *ε* = 82,900 M^−1^ cm^−1^, and *Φ*_f_ = 0.86). In summary, incorporation of the biotin moiety into the JF_549_-HaloTag ligand does not substantially affect the spectral properties of the free or HaloTag-bound ligand.

**Figure 3.**
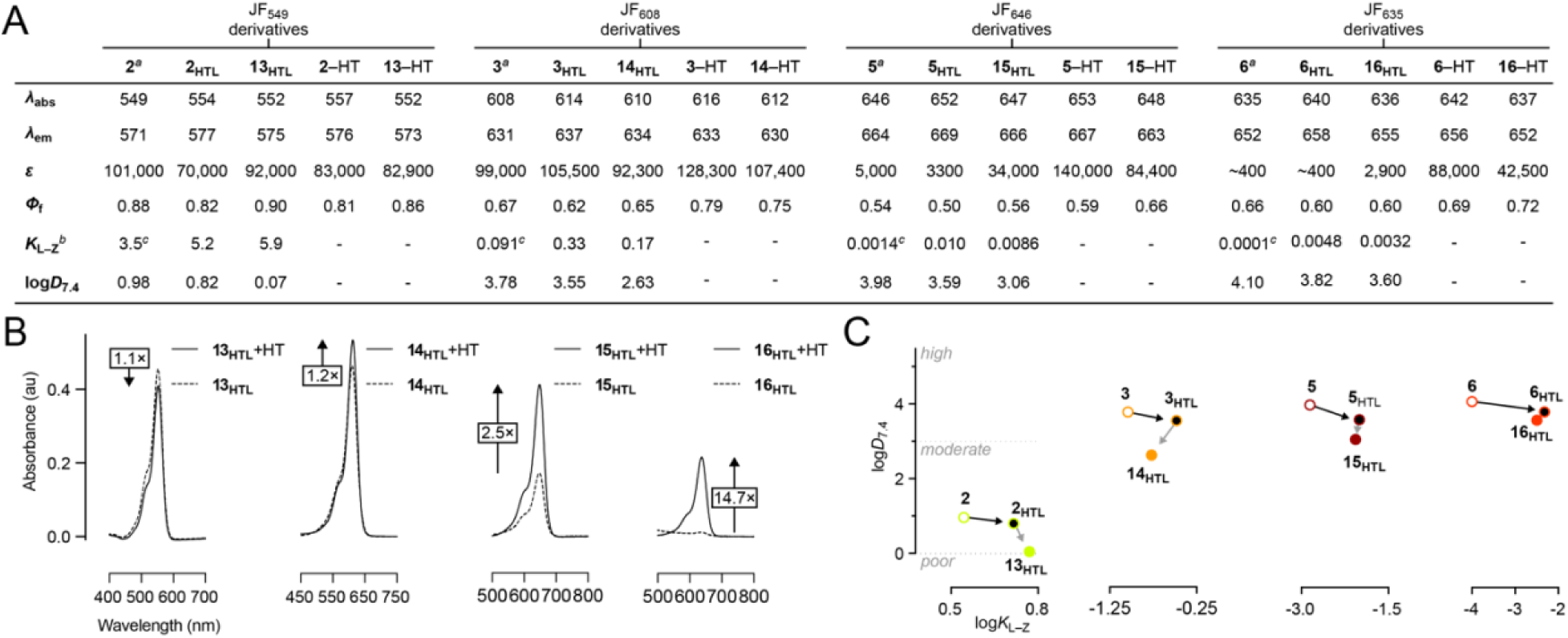
Evaluation of spectral and chemical properties, and live-cell labeling of biotin-rhodamine- HaloTag ligands. (A) Spectral properties, *K*_L–Z_ and *logD_7_*_.4_ of parent rhodamines (**2**–**6**), their HaloTag ligands (**2_HTL_**–**6_HTL_**), biotin-rhodamine-HaloTag ligands (**13_HTL_**–**16_HTL_**), and their HaloTag conjugates. Spectral measurements were taken in 10 mM HEPES, pH 7.3. ^a^Data taken from refs. ^27^ and ^43^. *λ*_abs_/*λ*_em_ are in nm, and *ε* are in M^−1^cm^−1^. ^b^*K*_L–Z_ measurements were performed in 1:1 (v/v) dioxane–water. ^c^Data taken from ref. ^27^ (B) Absorbance of **13_HTL_**–**16_HTL_** in the absence or presence (+HT) of excess HaloTag protein. (C) Correlation between log*D*_7.4_ versus log*K*_L–Z_. Dashed lines denote expected cellular permeability based on log*D*_7.4_.

The slight decrease in *λ*_abs_ and *λ*_em_ for the biotin-containing **13_HTL_** relative to **2_HTL_** was mirrored in the biotin- JF_608_-HaloTag ligand (**14_HTL_**), biotin-JF_635_-HaloTag ligand (**15_HTL_**), biotin-JF_646_-HaloTag ligand (**16_HTL_**) compared to their parent ligands **3_HTL_**–**6_HTL_** (**Fig. 3A**). Like their parent dyes and ligands, however, the absorptivity of **14_HTL_**–**16_HTL_**was strongly affected by environment. The carborhodamine-based biotin- JF_608_-HaloTag ligand (**14_HTL_**) shows *ε* = 92,300 M^−1^ cm^−1^ but conjugation to the HaloTag protein to yield **14**–HT moderately increases the absorptivity (*ε* = 107,400 M^−1^cm^−1^; **Fig 3A,B**). The absorptivity of Si-rhodamine-based biotin-JF_646_-HaloTag ligand (**15_HTL_**; *ε* = 34,000 M^−1^cm^−1^) increases to *ε* = 84,400 M^−1^cm^−1^ as the HaloTag conjugate **15**–HT. The biotin-JF_635_-HaloTag ligand (**16_HTL_**) exhibits a lower extinction coefficient (*ε* = 2,900 M^−1^cm^−1^) due to the fluorine-induced shift in the lactone–zwitterion equilibrium. The HaloTag adduct **16**–HT exhibits increased absorptivity with *ε* = 42,500 M^−1^cm^−1^. In addition to substantial increases in absorptivity upon binding the HaloTag protein, these carbo- and Si-rhodamine ligands show a modest increase in quantum yield. **14_HTL_** exhibits *Φ*_f_ = 0.65 and **14**–HT shows *Φ*_f_ = 0.75; **15_HTL_** exhibits *Φ*_f_ = 0.56 and **15**–HT shows *Φ*_f_ = 0.66; **16_HTL_** exhibits *Φ*_f_ = 0.60 and **16**–HT shows *Φ*_f_ = 0.72; Together, the changes in *ε* and *Φ*_f_ give degrees of fluorogenicity of 1.2-fold, 2.5-fold, and 14.7-fold for **14_HTL_**, **15_HTL_**, and **16_HTL_**, respectively (**Fig. 3B**).

We then measured the *K*_L–Z_ and log*D*_7.4_ values of the new biotin-containing HaloTag ligands (**Fig. 3A,C**). The biotin-JF_549_-HaloTag ligand (**13_HTL_**) exhibits a *K*_L–Z_ = 5.9, which is higher than both JF_549_ (**2**; *K*_L–Z_ = 3.5) and JF_549_-HaloTag ligand (**2_HTL_**; *K*_L–Z_ = 5.2), suggesting the polar biotin moiety can modestly stabilize the zwitterion form of the rhodamine, perhaps through direct interaction with the dye. This effect of biotin is also seen in the decreased log*D*_7.4_ = 0.07 measured for biotin-JF_549_-HaloTag ligand (**13_HTL_**), which places it below the optimal range for cell-permeant molecules. For ligands based on dyes with lower *K*_L–Z_ values (**14_HTL_**–**16_HTL_**), the incorporation of the biotin moiety caused a modest decrease in *K*_L–Z_ compared to the unfunctionalized HaloTag ligands **3_HTL_**–**6_HTL_** due to electron-withdrawing 3ʺ-carboxamide. The *K*_L–Z_ of carborhodamine-based biotin-JF_608_-HaloTag ligand (**14_HTL_**; *K*_L–Z_ = 0.17) is higher than the parent JF_608_ (**3**; *K*_L–Z_ = 0.091) but lower than the JF_608_-HaloTag ligand (**3_HTL_**; *K*_L–Z_ = 0.33). The Si-rhodamine-based biotin- JF_646_-HaloTag ligand follows the same trend: compound **15_HTL_** shows *K*_L–Z_ = 0.0086, higher than JF_646_ (**6**; *K*_L–Z_ = 0.0014), but lower than **5_HTL_** (*K*_L–Z_ = 0.010). The fluorine atoms in JF_635_-based ligands further decrease the *K*_L–Z_ values with biotin-JF_635_-HaloTag ligand (**16_HTL_**) exhibiting a *K*_L–Z_ = 0.0032, which is higher than free JF_635_ (**6**; *K*_L–Z_ = 0.0001) but lower than value than **6_HTL_**(*K*_L–Z_ = 0.0048). The lower *K*_L–Z_ values are reflected in the relatively high log*D*_7.4_ values measured for **14_HTL_** (log*D*_7.4_ = 2.63), **15 _HTL_** (log*D*_7.4_ = 3.06), or **16 _HTL_** (log*D*_7.4_ = 3.60). Overall, we find that incorporating biotin on rhodamine-HaloTag ligand has a modest effect on *K*_L–Z_ and decreases the log*D*_7.4_ (**Fig. 3C**). This decrease in log*D*_7.4_ is substantial for the JF_549_-based **13_HTL_** but more modest for compounds based on carborhodamines and Si-rhodamines.

**Live cell imaging with multifunctional ligands.** Based on these *in vitro* log*D*_7.4_ measurements, we predicted that the JF_549_ compound (**13_HTL_**; log*D*_7.4_ = 0.07) would exhibit poor cell permeability but the other compounds **14_HTL_**–**16_HTL_**(log*D*_7.4_ = 2.63–3.60) would readily enter cells. To test this, we evaluated biotin ligands **13_HTL_**–**16_HTL_**in U2OS cells transiently transfected with plasmids encoding the HaloTag sequence fused to the sequence of the following proteins: (i) <u>cell-surface</u>-localized platelet-derived growth factor receptor (PDGFR); (ii) outer <u>mitochondria</u> membrane localized TOMM20; (iii) <u>endoplasmic reticulum</u>-localized Sec61β; (iv) <u>nucleus</u>-localized histone H2B. Labeling HaloTag fusions at different subcellular locations was critical since some rhodamine derivatives inherently localize to specific subcellular locales.^53–55^ Carborhodamine and Si-rhodamine based biotin ligands **14_HTL_**–**16_HTL_**exhibit bright and robust labeling of HaloTag fusions at different subcellular locations, whereas biotin-JF_549_-HaloTag ligand (**13_HTL_**; log*D*_7.4_

= 0.07) exhibit poor intracellular labeling, only labeling the cell-surface-localized PDGFR-HaloTag fusion (**Fig. 4**). We note that the biotin-JF_549_-HaloTag ligand (**13_HTL_**) has a similar design to the recently reported biotin-TMR-HaloTag ligand, whose *in vitro* labeling kinetics approached the unmodified TMR-HaloTag ligand^21^ (**1_HTL_**); this TMR compound has not been evaluated in living cells. We measured the loading kinetics of **13_HTL_**–**16_HTL_** in U2OS cells stably expressing HaloTag–H2B (**Fig. S1**). Concurrent to the *in vitro* and imaging experiments, incubation of 200 nM **13_HTL_** results in negligible intracellular labeling over 4 h whereas the carborhodamine ligand (**14_HTL_**) and Si-rhodamine ligands (**15_HTL_** and **16_HTL_**) label intracellular proteins, reaching maximal labeling at 4 h and 1 h, respectively.

**Figure 4.**
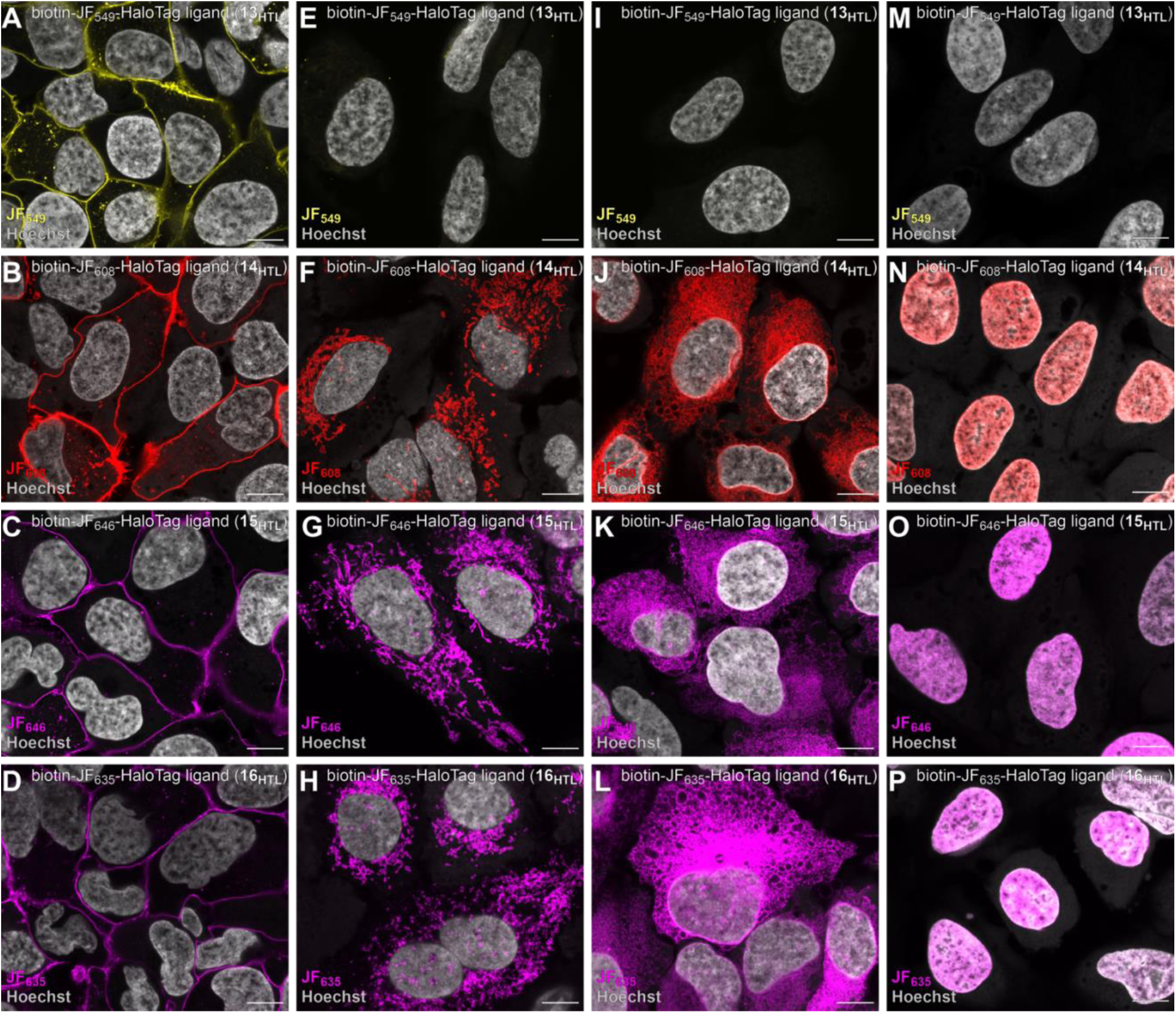
**Evaluation of live-cell labeling of biotin-rhodamine-HaloTag ligands 13_HTL_**–**16_HTL_.** (A–P) Airyscan fluorescence microscopy images of U2OS cells, expressing HaloTag fusions at different cellular locations, incubated with **13_HTL_**–**16_HTL_**; cells were fixed before imaging. (A–D) Cell-surfaced localized HaloTag–PDGFR. (E–H) Outer mitochondrial membrane localized HaloTag–TOMM20. (I–L) Endoplasmic reticulum membrane localized HaloTag–Sec61β. (M–P) Nucleus localized HaloTag–histone H2B. Cells were imaged after incubating with 100 nM for 1 h with biotin-JF_549_-HaloTag ligand (**13_HTL_**; A, E, I, M), biotin- JF_608_-HaloTag ligand (**14_HTL_**; B, F, J, N), biotin-JF_646_-HaloTag ligand (**15_HTL_**; C, G, K, O), or biotin-JF_635_- HaloTag ligand (**16_HTL_**; D, H, L, P) followed by fixation and counterstaining with Hoechst 33342. Image sets A/E/I/M, B/F/J/N, C/G/K/O, and D/H/L/P used the same microscope settings. Scale bar: 10 μm.

### Affinity purification using biotin-rhodamine-HaloTag ligands

We then applied these biotin-containing multifunctional ligands for affinity purification of mitochondria from HEK293T cells expressing a fusion protein consisting of the outer membrane protein 25 (OMP25), monomeric superfolder green fluorescent protein (msGFP), and the HaloTag (**Fig. 5A**). This construct localizes HaloTag to the outer mitochondrial membrane^56, 57^ and allows straightforward measurement of labeling and capture efficiency of biotin ligands (**13_HTL_–16_HTL_**) using a pulse–chase assay coupled with sodium dodecyl sulfate–polyacrylamide gel electrophoresis (SDS–PAGE) followed by in-gel fluorescence.

**Figure 5.**
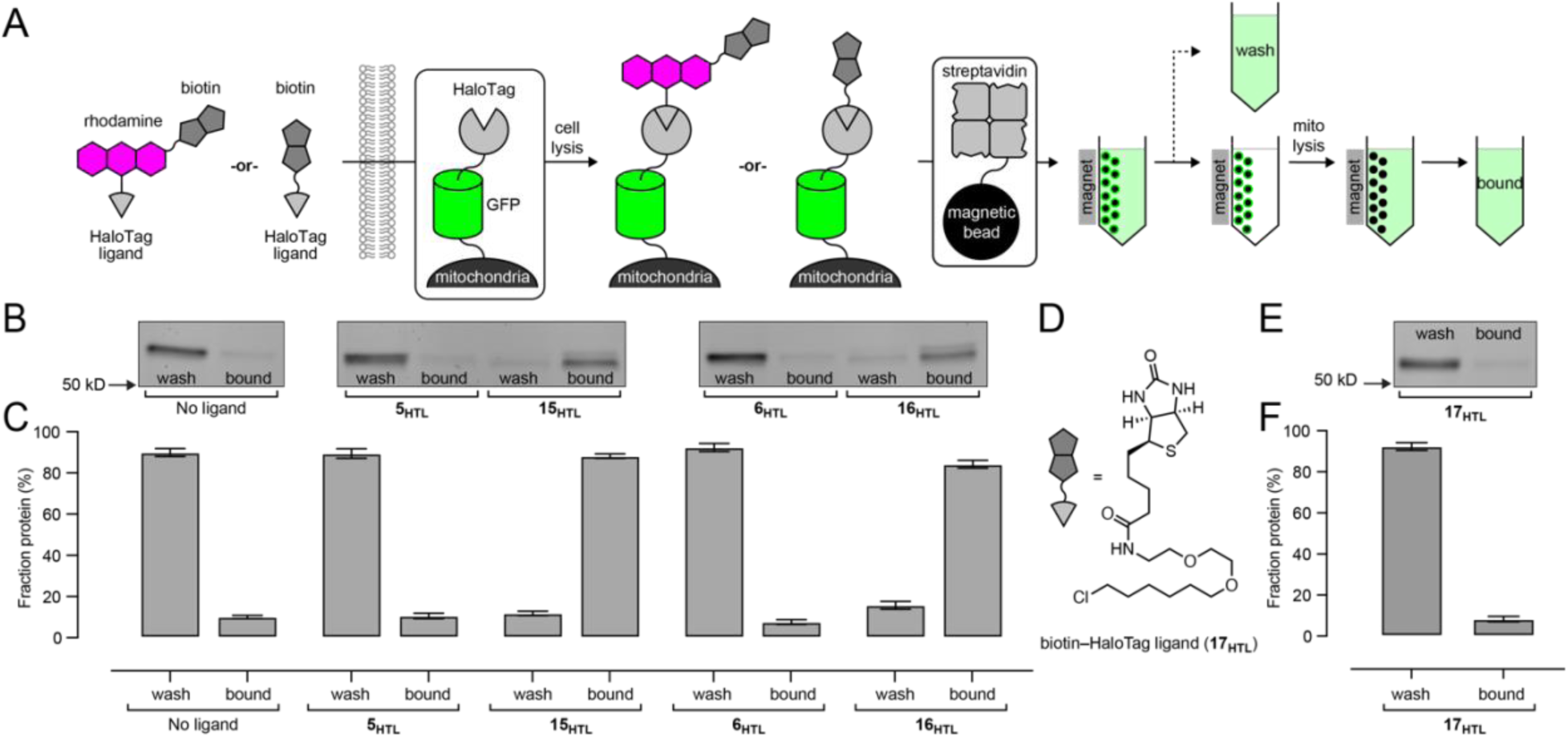
**Evaluation of Si-rhodamine containing biotin-JF_646_-HaloTag ligand (**15_HTL_**) and biotin-JF_635_- HaloTag ligand (**16_HTL_**), and commercial biotin-HaloTag ligand (**17_HTL_**) for affinity purification of mitochondria from HEK293T cells.** (A) Schematic of the assay to evaluate live-cell intracellular labeling and affinity purification of biotin–HaloTag conjugates. (B, C) SDS–PAGE/in-gel fluorescence (B) and quantification (C) showing the amount of msGFP–HaloTag fusion protein bound to streptavidin after labeling without any ligand, biotin-free parent ligands (**5_HTL_** or **6_HTL_**), and **15_HTL_** or **16_HTL_**. (D) Chemical structure of **17_HTL_**. (E, F) SDS–PAGE/in-gel fluorescence (E) and quantification (F) showing the amount of msGFP–HaloTag fusion protein bound to streptavidin after labeling with commercial biotin-HaloTag ligand **17_HTL_**. Data from at least three independent trials (mean ± SEM).

We first evaluated the labeling efficiency of biotin ligands (**13_HTL_–16_HTL_**) at 100 nM and 1 µM. We confirmed that 1 µM of parent ligands **2_HTL_–6_HTL_** showed near complete labeling of the msGFP–HaloTag fusion (**Fig. S2**). The biotin-JF_549_-HaloTag ligand (**13_HTL_**) gave negligible labeling (< 3%) at both 100 nM and 1 µM concentration (**Fig. S3**, **Fig. S4**), however, consistent with the log*D*_7.4_ measurements and imaging results in U2OS cells. The carborhodamine-based biotin-JF_608_-HaloTag ligand (**14_HTL_**) gave substantially higher labeling efficiency: 11.3 ± 6.4% (mean ± SEM; 100 nM) and 57.7 ± 9.6% (1 µM). The Si-rhodamine ligands provided highest overall labeling efficiency with JF_646_-based **15_HTL_** showing 13.6 ± 3.7% (100 nM) and 61.7 ± 6.8% (1 µM) and JF_635_-based **16_HTL_** yielding 18.0 ± 4.9% (100 nM) and 69.0 ± 4.2% (1 µM). Si-rhodamine ligands **15_HTL_** and **16_HTL_** also showed robust labeling in live HEK293T cells with signals that that overlapped with the mitochondrial-localized GFP (**Fig. S5**).

We used **15_HTL_** and **16_HTL_** to purify mitochondria. Cells were incubated with **15_HTL_** and **16_HTL_**, followed by washing and cell lysis. The crude supernatant was incubated with streptavidin-coated magnetic microbeads and washed. The bead-bound mitochondria were lysed, and the resulting solution was analyzed by SDS– PAGE/in-gel fluorescence (**Fig. 5B**). We verified that the parent, biotin-free ligands **5_HTL_** and **6_HTL_** gave no appreciable protein capture, but 100 nM biotin-JF_646_-HaloTag ligand (**15_HTL_**) and biotin-JF_635_-HaloTag ligand (**16_HTL_**) gave substantial capture efficiency of 84.2 ± 1.8% and 88.2 ± 1.1% (**Fig. 5C**). Use of 10- fold higher concentrations of **15_HTL_** and **16_HTL_**(1 µM) did not increase mitochondria pulldown efficiency (**Fig. S6**), demonstrating that the numerous HaloTag proteins on each mitochondrion facilitates efficient capture even without saturated labeling with a biotin-containing ligand.

We performed the same affinity protocol using the commercial biotin-HaloTag ligand (**17_HTL_**; **Fig. 5D**), which contains the standard PEG_2_-chloroalkane HaloTag ligand directly attached to the biotin carboxyl group. Pulse–chase labeling revealed that 1 µM of **17_HTL_**provides higher intracellular labeling (92.1 ± 0.4%; **Fig. S7**) compared to ligands **14_HTL_**–**16_HTL_**. Despite the increased degree of labeling, however, **17_HTL_**afforded negligible capture efficiency of mitochondria (8.6 ± 1.4%; **Fig. 5E**, **F**), similar to the no ligand control (7.0 ± 0.5%; **Fig. S8**). The capture efficiency remained low even with 10-fold higher concentration (10 µM; **Fig. S8**). We surmised that the linker in **17_HTL_** is too short to allow binding of the biotin–HaloTag conjugate to streptavidin for efficient affinity purification from HaloTag-expressing cells. This is consistent with the intimate association of ligand and protein in HaloTag conjugates (**Fig. 1C**). Attempts to remedy this by incorporating a longer PEG linker between the biotin and the chloroalkane yields commercially available **18_HTL_**, which exhibits low cell permeability,^58^ rendering it unsuitable for experiments in live cells (**Fig. S7**). These results demonstrate that the existing biotin ligands are suboptimal for streptavidin-mediated affinity purification using the HaloTag system due to poor cell permeability or insufficient linker length. In contrast, inserting a Si-rhodamine unit between the polar biotin and HaloTag ligand balances cellular permeability and functionality; biotin ligands **15_HTL_** and **16_HTL_** enter cells, efficiently label HaloTag proteins, enable visualization, and present a biotin moiety accessible for affinity capture (**Fig. 3–4**).

### Protein translocation using JQ1-rhodamine-HaloTag ligands

We extended this multifunctional ligand strategy by appending JF-HaloTag ligands with the pharmacological agent (*S*)-JQ1, a binder and inhibitor of the bromodomain and extra-terminal motif (BET) family protein BRD4. Our goal was to develop reagents that could be added to cells and cause rapid translocation of BRD4 to nuclear regions where the HaloTag protein was expressed (**Fig. 6A**). We focused on Si-rhodamine-based ligands since they are fluorogenic (**Fig. 3B**) and typically show efficient cellular labeling (**Fig. 4**), circumventing the need to wash out excess compound. As before,^35^ the 3-carboxyazetidine motif was used as a convenient attachment point for (*S*)-JQ1-PEG_2_-NH_2_ (**Scheme S3**), yielding JF_646_-based **19_HTL_** and JF_635_-based **20_HTL_** (**Fig. 6B** and **Scheme S4**).

**Figure 6.**
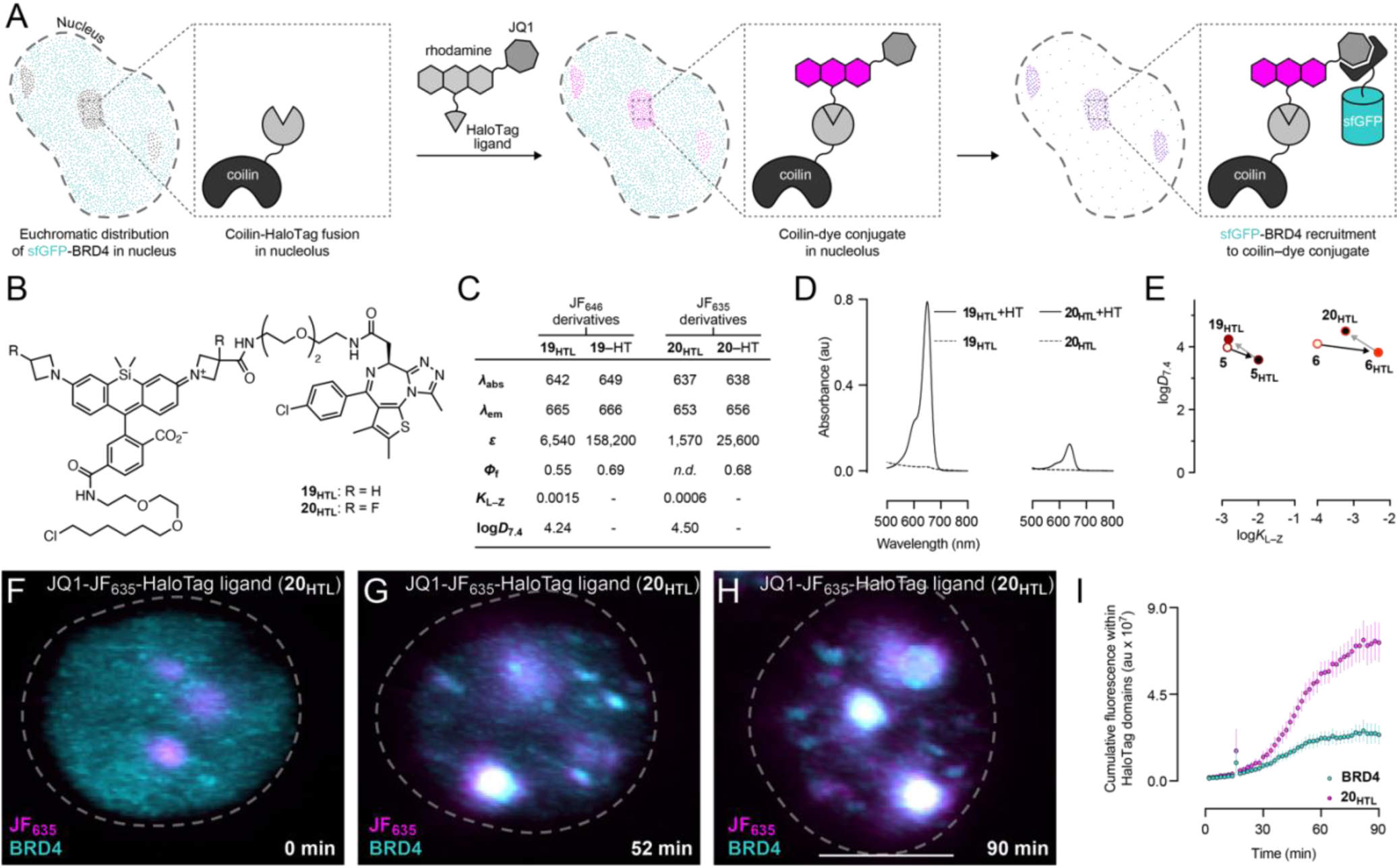
**Comparison of (*S*)-JQ1-JF_646_-HaloTag ligand (**19_HTL_**) and (*S*)-JQ1-JF_635_-HaloTag ligand (**20_HTL_**) for BRD4 translocation.** (A) Schematic illustrating the manipulation of BRD4 localization. (B) Chemical structure of (S)-JQ1-JF-HaloTag ligands **19_HTL_** and **20_HTL_**. (C) Spectral and chemical properties of **19_HTL_** and **20_HTL_**, and their HaloTag conjugates. All measurements were taken in 10 mM HEPES, pH 7.3. *λ*_abs_/*λ*_em_ are in nm, and *ε* are in M^−1^cm^−1^. *K*_L–Z_ measurements were performed in 1:1 (v/v) dioxane–water. (D) Absorbance of **19_HTL_** and **20_HTL_** in the absence or presence (+HT) of excess HaloTag protein. (E) Correlation between log*D*_7.4_ versus log*K*_L–Z_. (F–H) Extracted fluorescence from time course lattice light sheet microscopy in Neuro2A cells (n > 10 nuclei) expressing coilin–HaloTag and sfGFP–BRD4 upon incubation with **19_HTL_** or **20_HTL_**. Scale bar: 5 μm. (I) Cumulative rhodamine and BRD4 fluorescence within HaloTag domains (mean ± SEM).

We expected that the *λ*_abs_, *λ*_em_, and *Φ*_f_ of (*S*)-JQ1-JF_646_-HaloTag ligand (**19_HTL_**) and (*S*)-JQ1-JF_635_-HaloTag ligand (**20_HTL_**) would be analogous to their biotin-containing counterparts. We also surmised the lipophilicity of JQ1 would decrease *ε* and *K*_L–Z_ but increase log*D*_7.4_, although the degree of these changes is difficult to predict. Indeed, **19_HTL_** gave *λ*_abs_/*λ*_em_ = 642 nm/665 nm, *Φ*_f_ = 0.56—similar to biotin-JF_646_- HaloTag ligand (**15_HTL_**; **Fig. 6B**). The absorptivity was lower (*ε* = 6,540 M^−1^cm^−1^) as was the *K*_L–Z_ = 0.0015; the log*D*_7.4_ = 4.24 was higher than **15_HTL_**. Binding to the HaloTag (**19**–HT) confers the expected small bathochromic shift but a substantial increase in absorptivity and fluorescence quantum yield (*ε* = 158,200 M^−1^cm^−1^, *Φ*_f_ = 0.69). The fluorinated (*S*)-JQ1-JF_635_-HaloTag ligand (**20_HTL_**) shows lower *λ*_abs_/*λ*_em_ = 637 nm/653 nm and lower absorptivity (*ε* = 1,570 M^−1^cm^−1^) relative to **19_HTL_**; the small extinction coefficient prevented accurate measurement of fluorescence quantum yield of the free ligand. This JF_635_ compound showed *K*_L–Z_ = 0.0006 and log*D*_7.4_ = 4.50, similar to the free JF_635_-HaloTag ligand (**6_HTL_**). Binding to HaloTag increases absorptivity (*ε* = 25,600 M^−1^cm^−1^) with the **19**–HT conjugate exhibiting *Φ*_f_ = 0.68. Compounds **19_HTL_** and **20_HTL_** do not show substantial increases in absorption or fluorescence when incubated with recombinant BRD4 (**Fig. S9**), showing that fluorogenic effect observed in cells is mostly driven by HaloTag binding (**Fig. 6D**). We concluded that JQ1 exerts some effect on the properties of the molecule by lowering *K*_L–Z_ and increasing log*D*_7.4_, but, like the results with biotin-containing multifunctional ligands, this effect is relatively minor (**Fig. 6E**). The parent dye properties dominate in these multifunctional compounds even when attaching a hydrophobic moiety like JQ1.

We then tested the ability of JQ1 ligands to enter living cells and recruit BRD4 to defined genomic regions using lattice light sheet fluorescence microscopy (LLSM). Neuro2a cells were transiently transfected with plasmids encoding three fusion proteins: (i) HaloTag protein fused to coilin, which localizes to the nucleolus; (ii), superfolder GFP–BRD4; and (iii) histone H2B–mCherry as a nuclear marker. The Cajal body component coilin localizes in distinct puncta in the nucleus^59^ compared to the diffuse euchromatic distribution of BRD4. Upon addition to cells, both (*S*)-JQ1-JF_646_-HaloTag ligand (**19_HTL_**) and (*S*)-JQ1-JF_635_- HaloTag ligand (**20_HTL_**) rapidly labeled the coilin–HaloTag in a fluorogenic manner (**Fig. 6F–H** and **Fig. S11**) and caused effectively simultaneous BRD4 accumulation at coilin-rich sites in the nucleus (**Fig. 6I**), consistent with the rapid mobility of this protein.^60^ Ligand **19_HTL_** maintains higher brightness than **20_HTL_** (**Fig. S10**), consistent with the *in vitro* spectroscopic measurements (**Fig. 6C**). JF_646_-based **19_HTL_**labels faster than JF_635_-derived **20_HTL_**, perhaps due to the lower log*D*_7.4_, but **20_HTL_** ultimately results in higher BRD4 signal in the regions of interest (**Fig. S10**).

Given its lower fluorescence background and more efficacious recruitment of BRD4, we focused (*S*)-JQ1- JF_635_-HaloTag ligand (**20_HTL_**) for subsequent experiments. To test the generality of **20_HTL_** in recruiting BRD4, we imaged cells expressing heterochromatin protein 1a (HP1a) fused to HaloTag (**Fig. 7A**); HP1a is localized within constitutive heterochromatin due to its N-terminal chromodomain. Consistent with the experiments in cells expressing coilin–HaloTag, adding **20_HTL_** to cells with HP1a–HaloTag altered the uniform euchromatic distribution of BRD4 and localized it rapidly to HP1a (**Fig. 7B–D** and **Fig. S12**). Neither JF_635_-HaloTag ligand (**6_HTL_**), which lacks JQ1, nor (*R*)-JQ1-JF_635_-HaloTag ligand (**21_HTL_**; **Scheme S5**), which contains the inactive enantiomer (*R*)-JQ1, recruited BRD4 to HP1a–HaloTag (**Fig. 7E, Fig. S14**).

**Figure 7.**
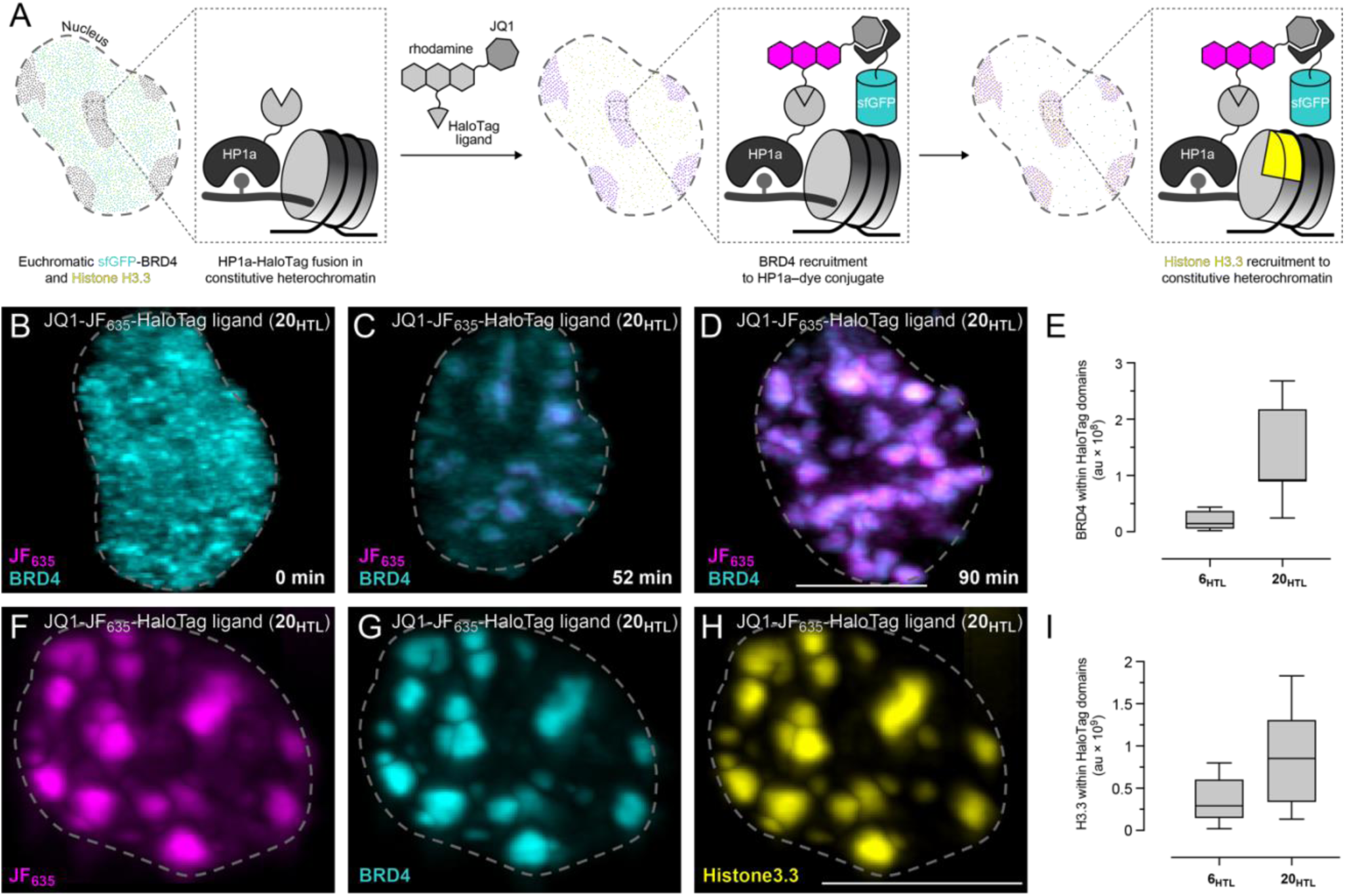
**Evaluation of (*S*)-JQ1-JF_635_-HaloTag ligand (**20_HTL_**) for altering chromatin.** (A) Schematic illustrating the translocation of BRD4 to constitutive heterochromatin and Histone H3.3 accumulation. (B– H) Data from time course lattice light sheet microscopy of Neuro2A cells (n > 10 nuclei) expressing HP1a– HaloTag and sfGFP–BRD4 after incubation with **20_HTL_** or **6_HTL_**. (B–D) Representative maximum intensity projections at mentioned times after incubation with **20_HTL_**. (E) Box and Whisker plot of cumulative BRD4 within HaloTag domains after incubation with **6_HTL_** or **20_HTL_**. (F–H) Representative maximum intensity projections after incubation with **20_HTL_** and 1 µM JFX_554_-SNAP-tag ligand (**22_HTL_**) from cells also expressing histone H3.3–SNAP-tag. (I) Box and Whisker plot of cumulative H3.3 within HaloTag domains after incubation with **6_HTL_** or **20_HTL_** from cells also expressing histone H3.3–SNAP-tag. The dashed line represents the nuclear boundary. Scale bars: 5 μm

Finally, we assessed whether recruitment of BRD4 to transcriptionally repressed constitutive heterochromatin domain has functional consequences. Cells expressing sfGFP–BRD4, HP1a–HaloTag, and histone H3.3–SNAP-tag^14^ were incubated with **20_HTL_** and JFX_554_-SNAP-tag ligand^31^ (**22_HTL_**). As before, we observed rapid translocation of BRD4 to HP1a–HaloTag domains, but we also saw depletion of histone- H3.3 from euchromatin and accumulation of H3.3 in the same subnuclear regions (**Fig. 7F–H**); this effect required the JQ1 functionality on the ligand (**Fig. 7I** and **Fig. S15**). Histone H3.3 is usually absent in constitutive heterochromatin and accumulates in transcriptionally active nucleosomes.^61–63^ This movement of histone H3.3 to heterochromatin upon BRD4 recruitment is consistent with BRD4’s role in releasing paused RNA polymerase Pol II^64^ and this result demonstrates that targeting JQ1 using **20_HTL_** to HP1a can substantially alter chromatin in living cells.

## Conclusion

The ability to label specific proteins with synthetic small-molecules in living cells is a powerful technique for biology.^1–3^ The generality of these systems has led to the development of many ligands to probe, perturb, or purify cellular components (**Fig. 1**). Nevertheless, designing cell-permeable ligands remains challenging, especially those with multiple functionalities. We took a medicinal chemistry approach by first measuring the distribution coefficients (log*D*_7.4_) of free rhodamine dyes and their HaloTag ligands. We compared these values against lactone–zwitterion equilibrium constants (*K*_L–Z_) and found an inverse correlation. This suggested that dyes with low *K*_L–Z_ values, and therefore high distribution coefficients, could be useful scaffolds for creating cell-permeable multifunctional ligands (**Fig. 2**). We tested this idea by synthesizing biotin-containing HaloTag ligands based on four different dyes with decreasing *K*_L–Z_ values: JF_549_, JF_608_, JF_646_, and JF_635_ (**13_HTL_**–**16_HTL_**; **Fig. 3**). Ligand **13_HTL_**, based on the high *K*_L–Z_/low log*D*_7.4_ dye JF_549_ showed poor intracellular labeling, whereas ligands **14_HTL_**–**16_HTL_**, based on the low *K*_L–Z_/high log*D*_7.4_ dyes JF_608_, JF_646_, and JF_635_, showed more efficient labeling of various subcellular locations (**Fig. 4**). Ligands **15_HTL_** and **16_HTL_** permitted efficient affinity capture of mitochondria in contrast to the commercial biotin-HaloTag ligand (**Fig. 5**). We extended this strategy to prepare fluorogenic JQ1-containing ligands **19_HTL_** and **20_HTL_**, which efficiently translocated BRD4 from euchromatin to subnuclear regions containing coilin (**Fig. 6**) or the constitutive heterochromatin marker HP1a. The recruitment of BRD4 to HP1a appears to make heterochromatin transcriptionally more active as evidenced by a corresponding increase in histone H3.3 (**Fig. 7**).

Looking forward, we expect the use of low *K*_L–Z_/high log*D*_7.4_ dyes as scaffolds will generate a palette of useful multifunctional ligands for biology. Although incorporating a rhodamine into a molecule is not without cost, advances in fluorophore chemistry over the past two decades,^40, 55, 65^ have simplified the synthesis of rhodamines. We show this additional effort is warranted—adding the Si-rhodamine moieties in biotin-containing ligands **15_HTL_** and **16_HTL_**was critical in allowing efficient capture of mitochondria as the commercial ligands **17_HTL_** or **18_HTL_** were either too short or too polar to give appreciable affinity capture after incubation with living cells. The inclusion of a fluorophore also allowed facile quantification of the biotin labeling in living cells through fluorescence microscopy and capture efficiency using SDS–PAGE. For JQ1-containing ligands **19_HTL_** and **20_HTL_**, having a fluorogenic dye was necessary to observe both the labeling of HaloTag fusion proteins and the translocation of BRD4 inside living cells without intermediate washing steps. Overall, designing other multifunctional ligands by leveraging the structure–activity relationships of rhodamines^26, 28–30, 40, 66–68^ will yield a valuable toolbox of probes where what you see is what you get.

## Supporting Information

Synthetic schemes, supplementary Figures, experimental details, and characterization for all new compounds (PDF)

## Acknowledgements

We are indebted to Dr Sharon King and Dr Rebecca Petersen of the Department of Developmental Neurobiology Neuroimaging Laboratory (St. Jude Children’s Research Hospital), for maintaining and aligning the instruments used in this study’s lattice-light sheet imaging sessions. This article is subject to HHMI’s Open Access to Publications policy. HHMI lab heads have previously granted a nonexclusive CC BY 4.0 license to the public and a sublicensable license to HHMI in their research articles. Pursuant to those licenses, the author-accepted manuscript of this article can be made freely available under a CC BY 4.0 license immediately upon publication.

## Preprint Servers

This manuscript was deposited as a preprint on bioRxiv under the CC-BY 4.0 International license.

## Author Contributions

This manuscript was written with contributions from all authors.

## Classification

Chemistry (major), Cell Biology (minor)

## This PDF file includes

Main Text, Scheme 1, and Figures 1 to 7

