## Supplementary Information for "Optimizing multifunctional fluorescent ligands for intracellular labeling"

SUPPLEMENTARY FIGURES AND SCHEMES ..... S2
SPECTROSCOPY AND CELL BIOLOGY METHODS ..... S20
SYNTHETIC ORGANIC CHEMISTRY METHODS ..... S27
EXPERIMENTALS AND CHARACTERIZATION FOR ALL NEW COMPOUNDS ..... S28
NMR SPECTRA AND HPLC TRACES ..... S41
REFERENCES ..... S60

#### Supplementary Figures and Schemes

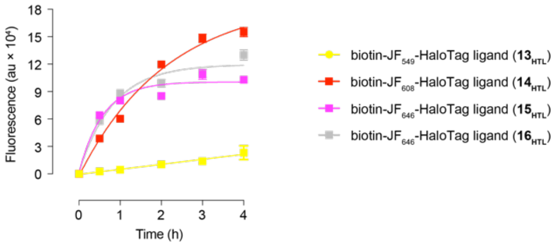


**Figure S1.** Loading curves for biotin-JF-HaloTag ligands (**$13_{HTL}$–$16_{HTL}$**) in live U2OS.H2B.HaloTag stable cells. Cells were treated with 200 nM of the ligand: biotin-$JF_{549}$-HaloTag ligand (**$13_{HTL}$**, yellow); biotin-$JF_{608}$-HaloTag ligand (**$14_{HTL}$**, red); biotin-$JF_{646}$-HaloTag ligand (**$15_{HTL}$**, magenta); biotin-$JF_{635}$-HaloTag ligand (**$16_{HTL}$**, gray).

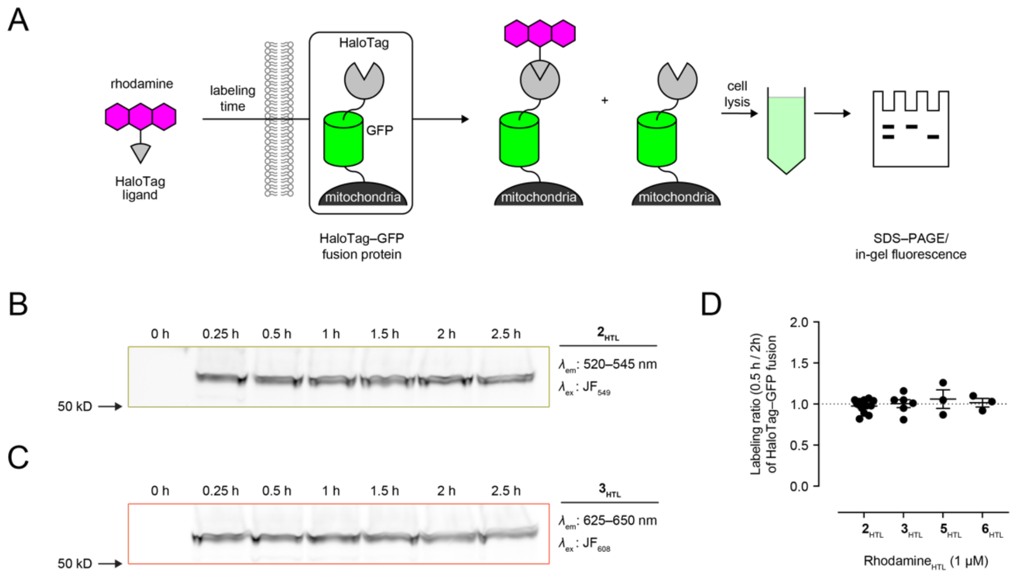


**Figure S2.** Time course evaluation of JF-HaloTag ligands for intracellular labeling in live HEK293T cells expressing msGFP–HaloTag fusion localized to the mitochondrial outer membrane. (*A*) Schematic of the experimental design to evaluate time-dependent labeling after incubation with $JF_{549}$-HaloTag ligand (**2**$_{\mathbf{HTL}}$), $JF_{608}$-HaloTag ligand (**3**$_{\mathbf{HTL}}$), $JF_{646}$-HaloTag ligand (**5**$_{\mathbf{HTL}}$), and $JF_{635}$-HaloTag ligand (**6**$_{\mathbf{HTL}}$). (*B, C*) Representative images for SDS-PAGE/in-gel fluorescence analyses of cell lysate after incubating living cells with **2**$_{\mathbf{HTL}}$ (*B*) or **3**$_{\mathbf{HTL}}$ (*C*) for given duration. (*D*) Fluorescence ratio of labeling at 0.5 h and 2 h shows complete labeling of HaloTag fusion in living cells after 0.5 h of incubation with 1 μM JF-HaloTag ligand. Data from at least three independent trials; solid lines represent mean, and error bars represent ± SEM.

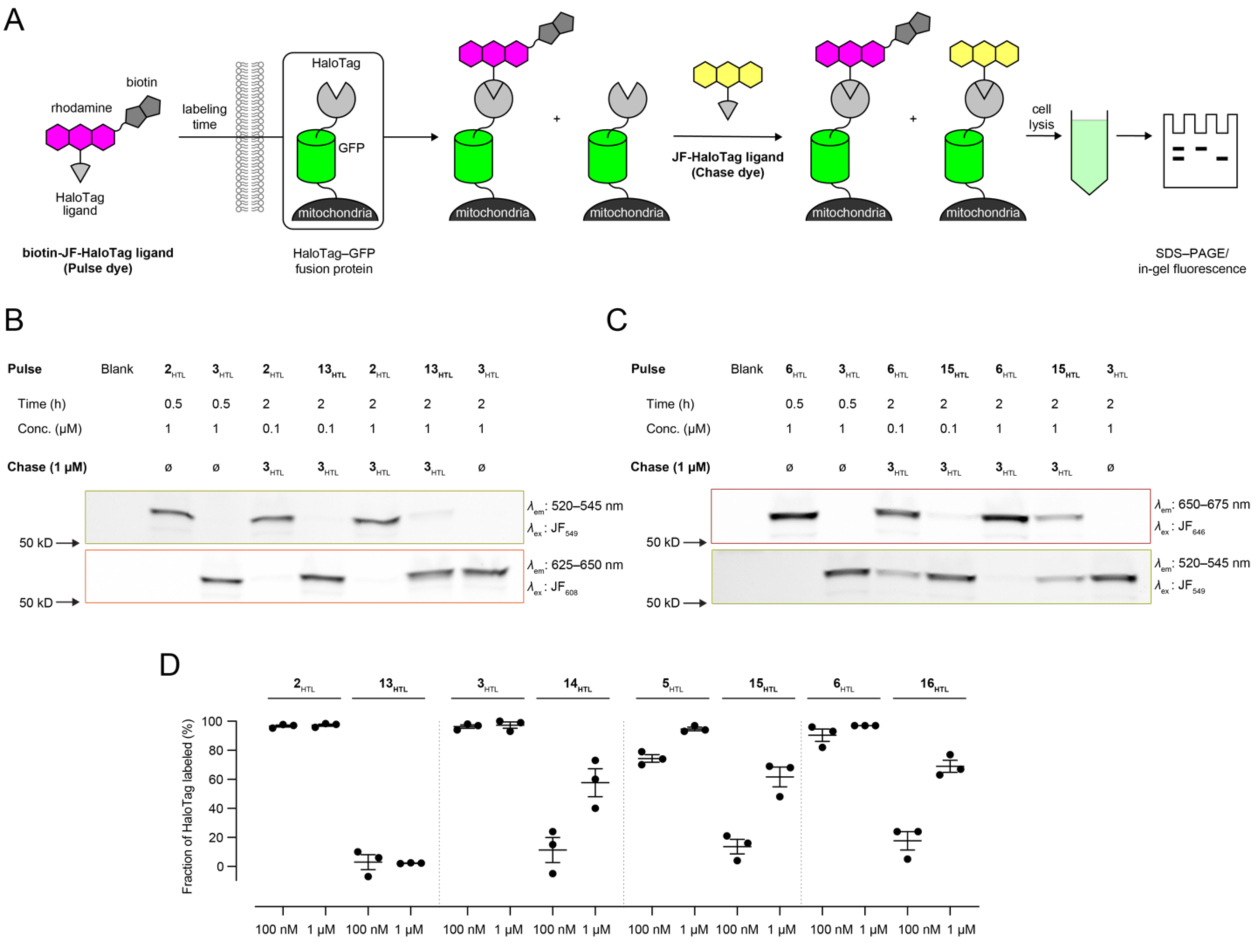


**Figure S3.** Evaluation of intracellular labeling efficiency of biotin-JF-HaloTag ligands (**$13_{HTL}$–$16_{HTL}$**) in live HEK293T cells expressing msGFP–HaloTag fusion localized to the mitochondrial outer membrane. (*A*) Schematic of the assay to evaluate the labeling efficiency of HaloTag–msGFP fusion localized to the mitochondrial outer membrane after incubation with biotin-JF-HaloTag ligands (**$13_{HTL}$–$16_{HTL}$**; pulse) and JF-HaloTag ligands (**$2_{HTL}$** or **$3_{HTL}$**; chase). (*B, C*) Representative image for SDS-PAGE/in-gel fluorescence analyses for determining labeling efficiency of biotin-$JF_{549}$-HaloTag ligand (**$13_{HTL}$**) and (*C*) biotin-$JF_{646}$-HaloTag ligand (**$15_{HTL}$**). (*D*) Quantification of fraction of HaloTag–msGFP labeled by 100 nM or 1 μM pulse ligand after 2 h incubation. Data from at least three independent trials; solid lines represent mean, and error bars represent ± SEM.

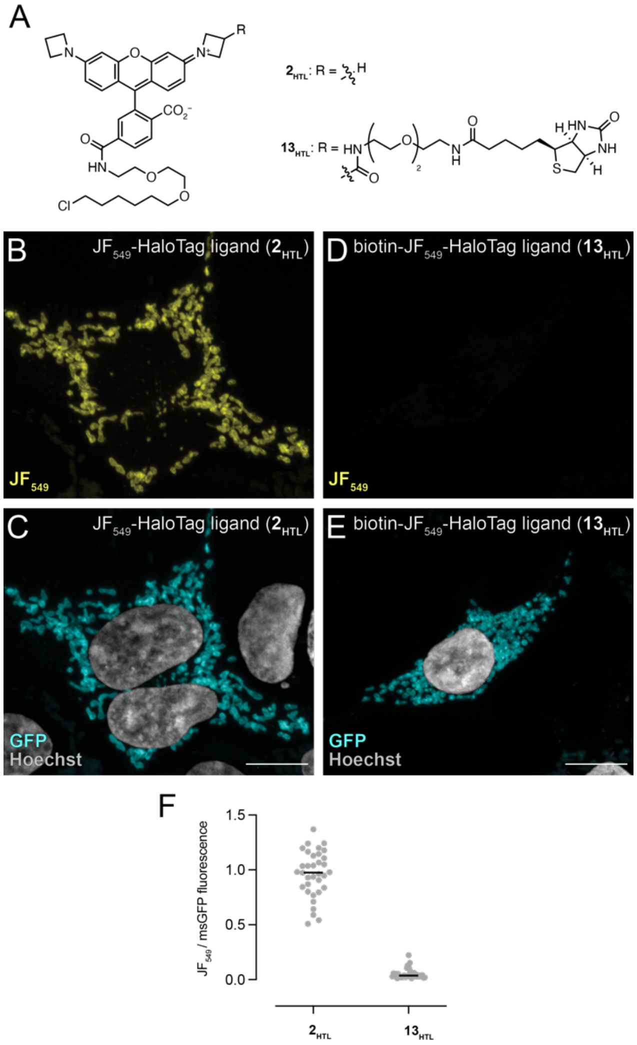


**Figure S4.** Evaluation of biotin-JF549-HaloTag ligand (**13HTL**) for intracellular labeling in live HEK293T cells expressing msGFP–HaloTag fusion localized to the mitochondrial outer membrane. (*A*) Chemical structures of the JF549-HaloTag ligand (**2HTL**) and biotin-JF549-HaloTag ligand (**13HTL**). (*B–E*) Airyscan fluorescence microscopy images of cells after incubation with either **2HTL** (*B, C*) or **13HTL** (*D, E*) and counterstained with Hoechst 33342; cells were fixed before imaging. Images in *B* and *D* used the same microscope settings; scale bars: 10 µm. (*F*) Quantification from intracellular fluorescence intensity ratio, from the live-cell imaging experiments, of HaloTag fusion labeled with JF549-HaloTag ligand (**2HTL**; 100 nM) or biotin-JF549-HaloTag ligand (**13HTL**; 100 nM).

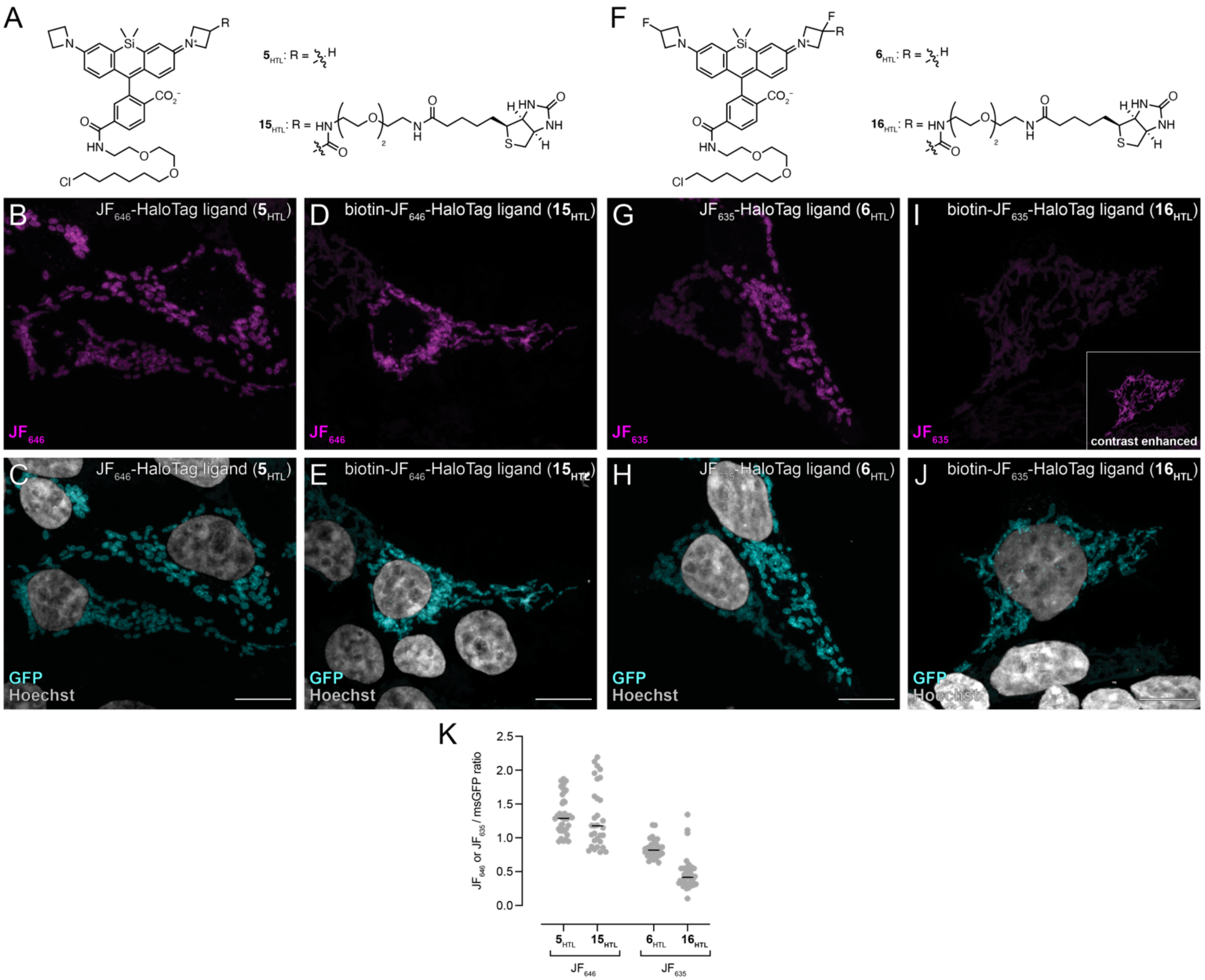

**Figure S5.** Evaluation of biotin-JF$_{646}$-HaloTag ligand (**15$_{HTL}$**) and biotin-JF$_{635}$-HaloTag ligand (**16$_{HTL}$**) for intracellular labeling in live HEK293T cells expressing msGFP–HaloTag fusion localized to the mitochondrial outer membrane. (*A*) Chemical structures of **15$_{HTL}$** and JF$_{646}$-HaloTag ligand (**5$_{HTL}$**). (*B–E*) Airyscan fluorescence microscopy images of cells incubated with **5$_{HTL}$** (*B, C*) or **15$_{HTL}$** (*D, E*) and counterstained with Hoechst 33342; cells were fixed before imaging. (*F*) Chemical structures of **16$_{HTL}$** and JF$_{635}$-HaloTag ligand (**6$_{HTL}$**). (*G–J*) Airyscan fluorescence microscopy images of cells incubated with **6$_{HTL}$** (*G, H*) or **16$_{HTL}$** (*I, J*) and counterstained with Hoechst 33342; cells were fixed before imaging. Image sets *B/D/G/I* and *C/E/H/J* each used the same microscope settings; scale bars: 10 µm. (*K*) Quantification from intracellular fluorescence intensity ratio, from the live-cell imaging experiments, of HaloTag fusion labeled with **5$_{HTL}$**, **15$_{HTL}$**, **6$_{HTL}$**, or **16$_{HTL}$**; *n* = 36 (**5$_{HTL}$**), *n* = 31 (**15$_{HTL}$**), *n* = 30 (**6$_{HTL}$**), and *n* = 36 (**16$_{HTL}$**), taken from three independent experiments per ligand with each *n* representing a delineated cell; solid lines represent mean. **Note**: both JF$_{635}$ ligands have lower intensity than the JF$_{646}$ ligands under equivalent imaging conditions. These results are consistent with the *in vitro* spectral characterization of the HaloTag conjugates.

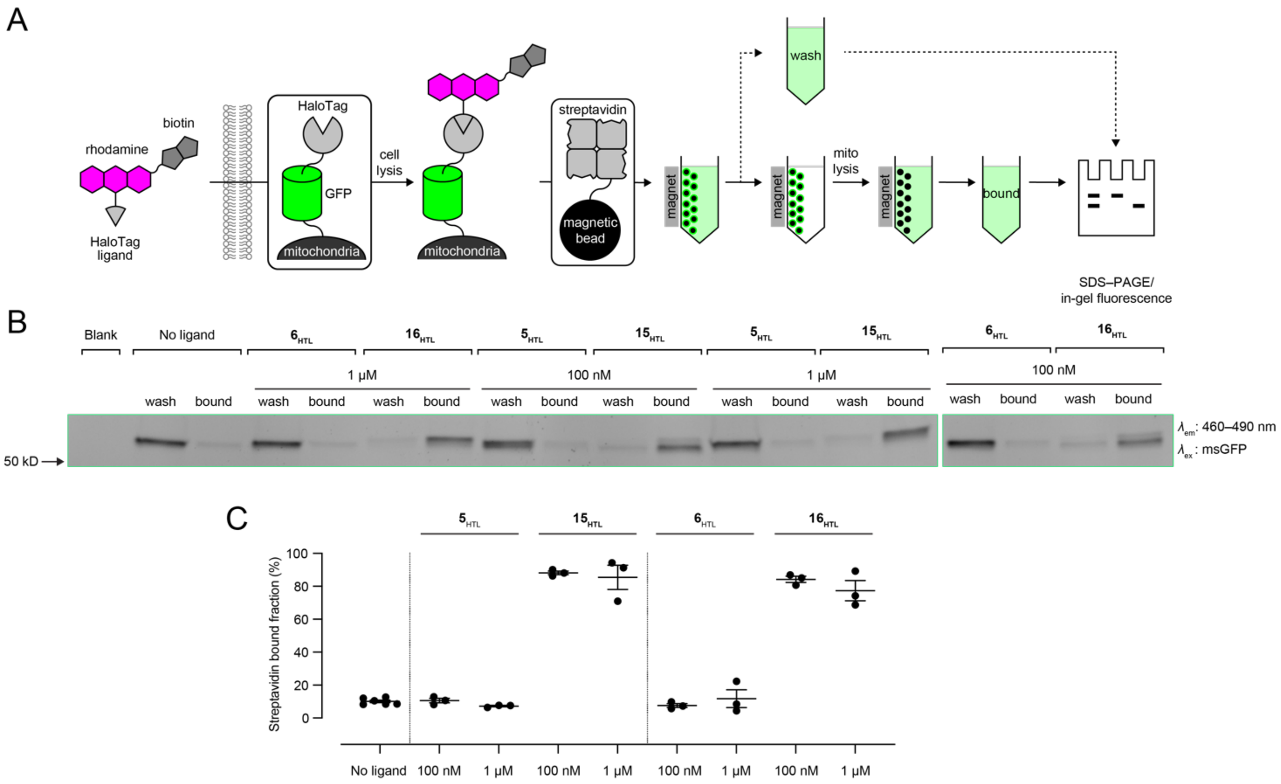


**Figure S6.** Evaluation of biotin-JF$_{646}$-HaloTag ligand (**15$_{\mathbf{HTL}}$**) and biotin-JF$_{635}$-HaloTag ligand (**16$_{\mathbf{HTL}}$**) at 100 nM or 1 µM for affinity purification of mitochondria from HEK293T cells. (*A*) Schematic of the assay to evaluate live-cell intracellular labeling and affinity purification of biotin–HaloTag conjugates. (*B, C*) Representative image for SDS–PAGE/in-gel fluorescence (*B*) and quantification (*C*) showing the amount of msGFP–HaloTag fusion protein bound to streptavidin after labeling without any ligand, biotin-free parent ligands (**5$_{\mathbf{HTL}}$** or **6$_{\mathbf{HTL}}$**), and biotin-containing ligands **15$_{\mathbf{HTL}}$** or **16$_{\mathbf{HTL}}$**. Data from at least three independent trials; solid lines represent mean, and error bars represent ± SEM.

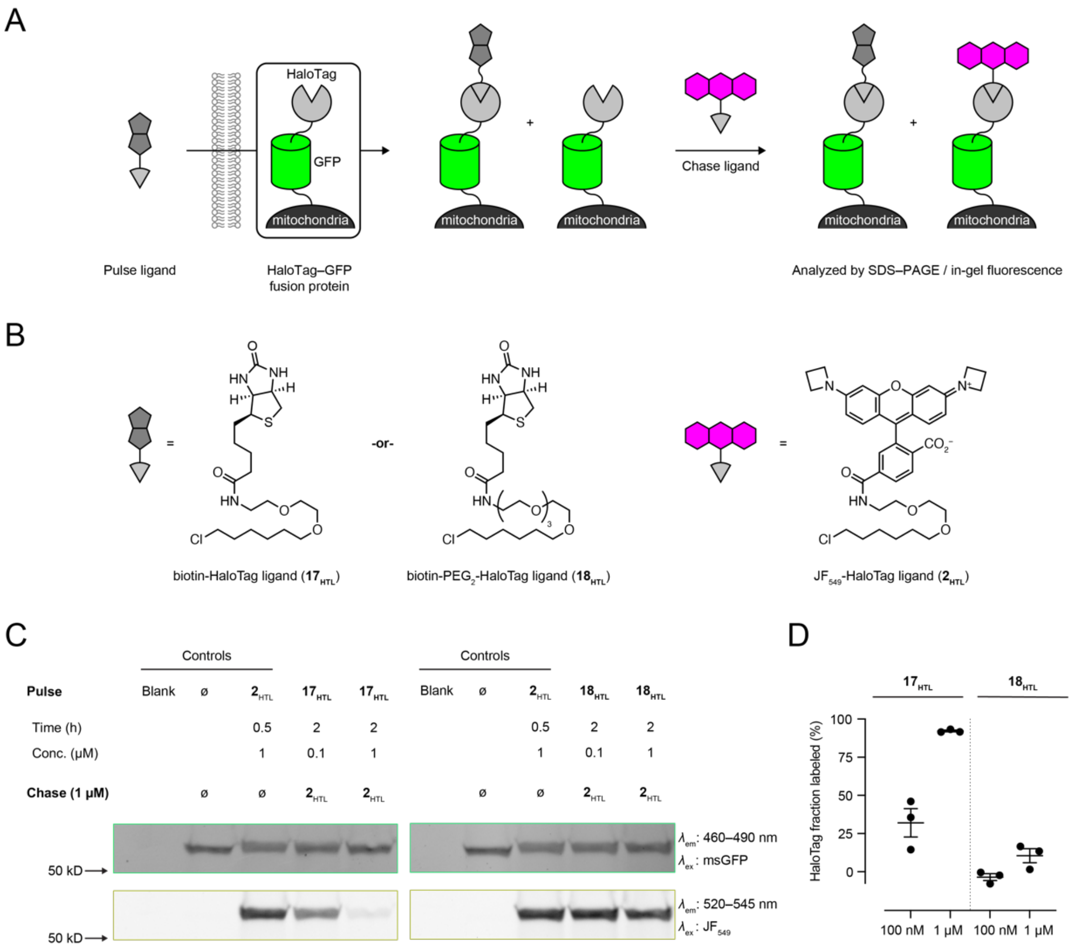


**Figure S7.** Evaluation of cellular permeability and HaloTag labeling of biotin-HaloTag ligand (**$17_{HTL}$**) and biotin-PEG$_2$-HaloTag ligand (**$18_{HTL}$**) in live HEK293T cells expressing msGFP–HaloTag fusion localized to the mitochondrial outer membrane. (*A*) Schematic of the pulse–chase assay to determine the labeling efficiency of **$17_{HTL}$** or **$18_{HTL}$** using the fluorescent JF$_{549}$-HaloTag ligand (**2**$_{HTL}$; chase). (*B*) Chemical structures of **$17_{HTL}$**, **$18_{HTL}$** and **$2_{HTL}$**. (*C, D*) Representative image for SDS-PAGE/in-gel fluorescence analyses for determining labeling efficiency of **$17_{HTL}$** and **$18_{HTL}$**. (*D*) Quantification of fraction of HaloTag–msGFP labeled by 100 nM or 1 μM pulse ligand after 2 h incubation. Data from at least three independent trials; solid lines represent mean, and error bars represent ± SEM.

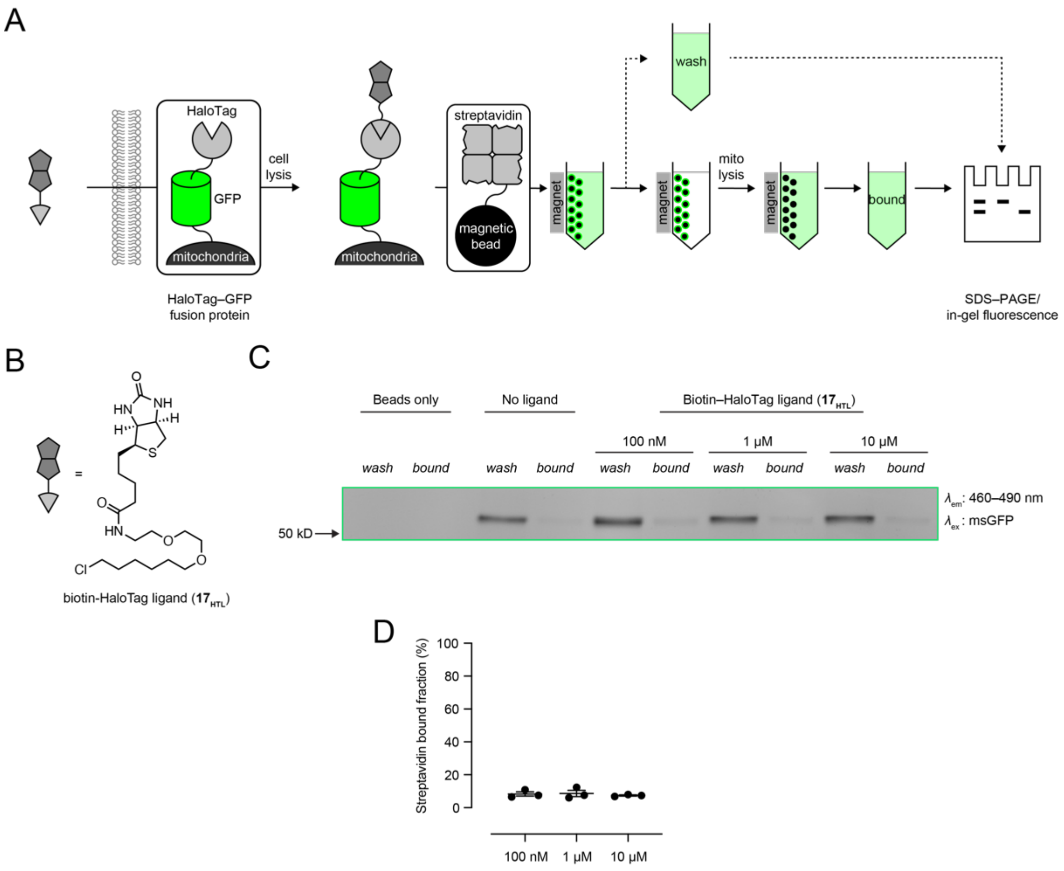


**Figure S8.** Evaluation of commercial biotin-HaloTag ligand (**17**$_{\mathbf{HTL}}$) for affinity purification of mitochondria from HEK293T cells. (*A*) Schematic of the assay to determine the streptavidin purification efficiency of biotin–HaloTag conjugates. (*B*) Chemical structure of **17**$_{\mathbf{HTL}}$. (*C, D*) SDS–PAGE/in-gel fluorescence (*C*) and quantification (*D*) showing the amount of msGFP–HaloTag fusion protein bound to streptavidin after labeling without any ligand or different concentrations of **17**$_{\mathbf{HTL}}$. Data from at least three independent trials; solid lines represent mean, and error bars represent ± SEM.

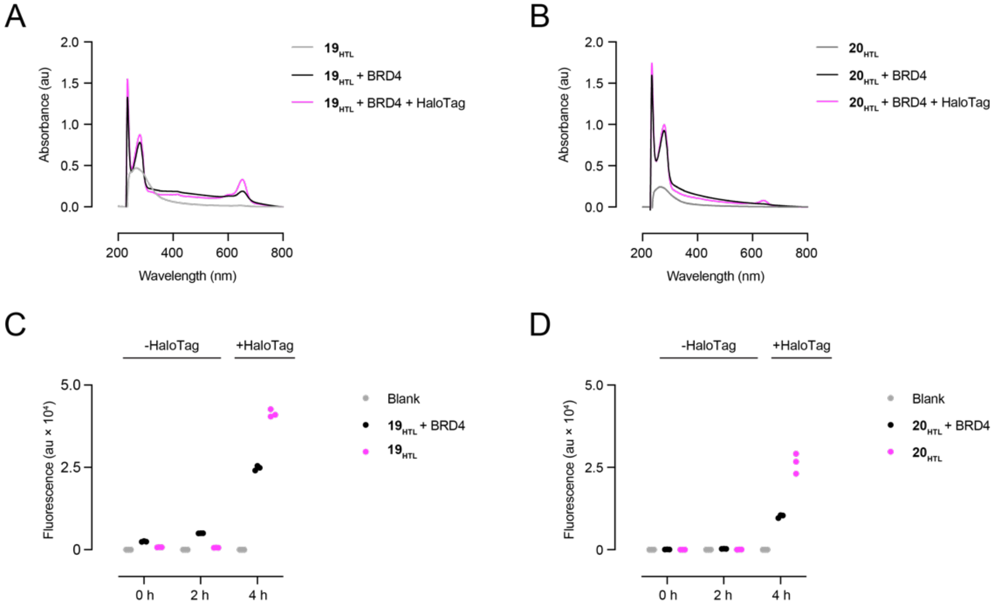


**Figure S9.** Spectroscopic measurements of incubation of (*S*)-JQ1-JF$_{646}$-HaloTag ligand (**19$_{HTL}$**) and (*S*)-JQ1-JF$_{635}$-HaloTag ligand (**20$_{HTL}$**) with purified BRD4. Absorbance spectra of 5 µM (*A*) **19$_{HTL}$** and (*B*) **20$_{HTL}$** upon incubation with 10 µM BRD4 and 10 µM HaloTag. End-point fluorescence measurements of (*C*) **19$_{HTL}$** and (*D*) **20$_{HTL}$** upon incubation with BRD4 and HaloTag.

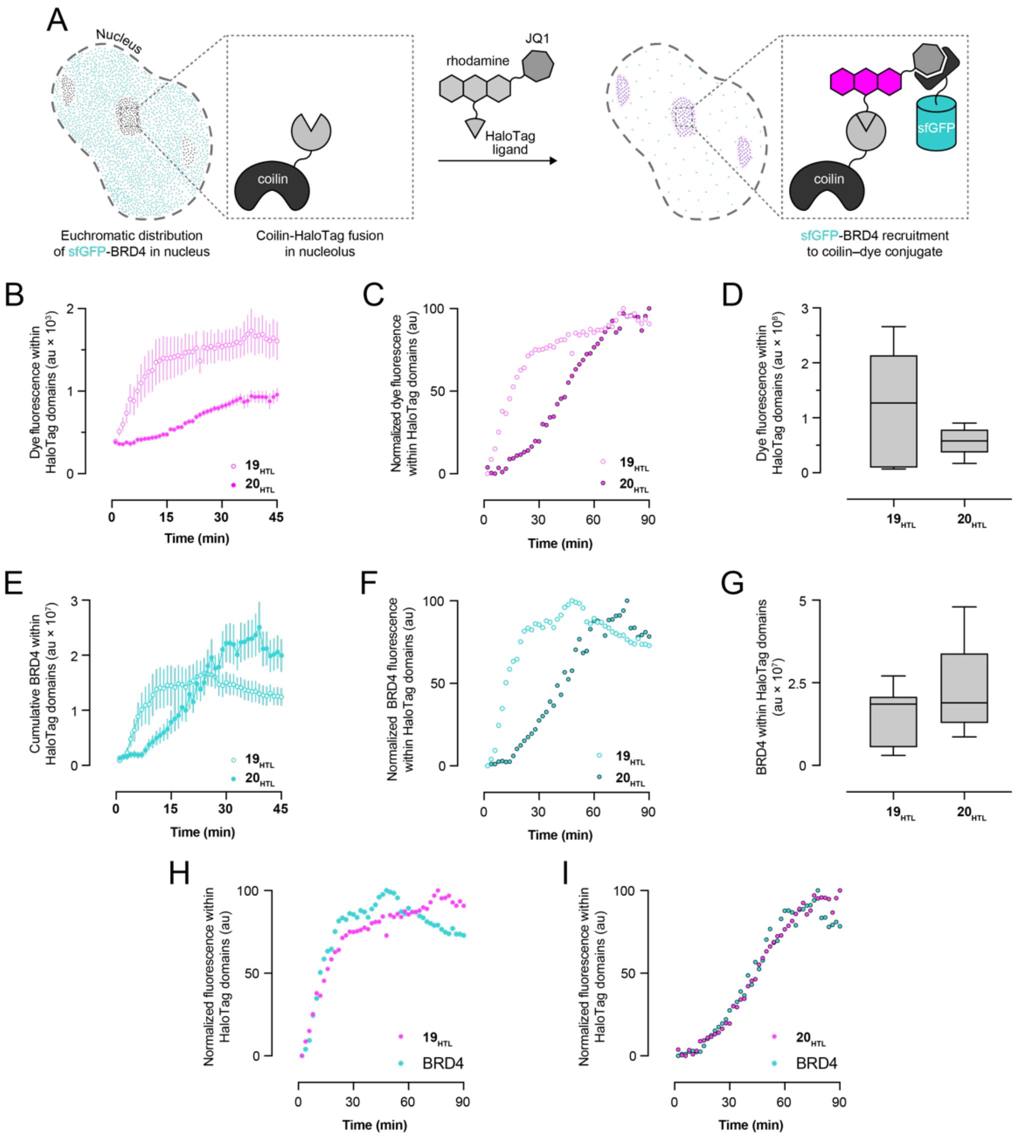


**Figure S10.** Comparison of (*S*)-JQ1-JF$_{646}$-HaloTag ligand (**19$_{\mathbf{HTL}}$**) and (*S*)-JQ1-JF$_{635}$-HaloTag ligand (**20$_{\mathbf{HTL}}$**) for BRD4 translocation using time course lattice light sheet microscopy in Neuro2A cells (n > 10 nuclei) expressing coilin–HaloTag and sfGFP–BRD4. (*A*) Schematic illustrating the assay to manipulate BRD4 localization. (*B*) Extracted rhodamine fluorescence (mean ± SEM) within HaloTag domains upon incubating cells with **19$_{\mathbf{HTL}}$** or **20$_{\mathbf{HTL}}$**. (*C*) Normalized rhodamine fluorescence within HaloTag domains upon incubating cells with **19$_{\mathbf{HTL}}$** or **20$_{\mathbf{HTL}}$**. Only mean values are shown for clarity. (*D*) Box and Whisker plot of rhodamine fluorescence within HaloTag domains. (*E*) Extracted cumulative BRD4 fluorescence (mean ± SEM) within HaloTag domains upon incubating cells with **19$_{\mathbf{HTL}}$** or **20$_{\mathbf{HTL}}$**. (*F*) Normalized BRD4 fluorescence within HaloTag domains upon incubating cells with **19$_{\mathbf{HTL}}$** or **20$_{\mathbf{HTL}}$**. Only mean values are shown for clarity. (*G*) Box and Whisker plot of cumulative BRD4 fluorescence within HaloTag domains. (*H, I*) Normalized rhodamine and BRD4 fluorescence upon incubating cells with **19$_{\mathbf{HTL}}$** (*H*) or **20$_{\mathbf{HTL}}$** (*I*).

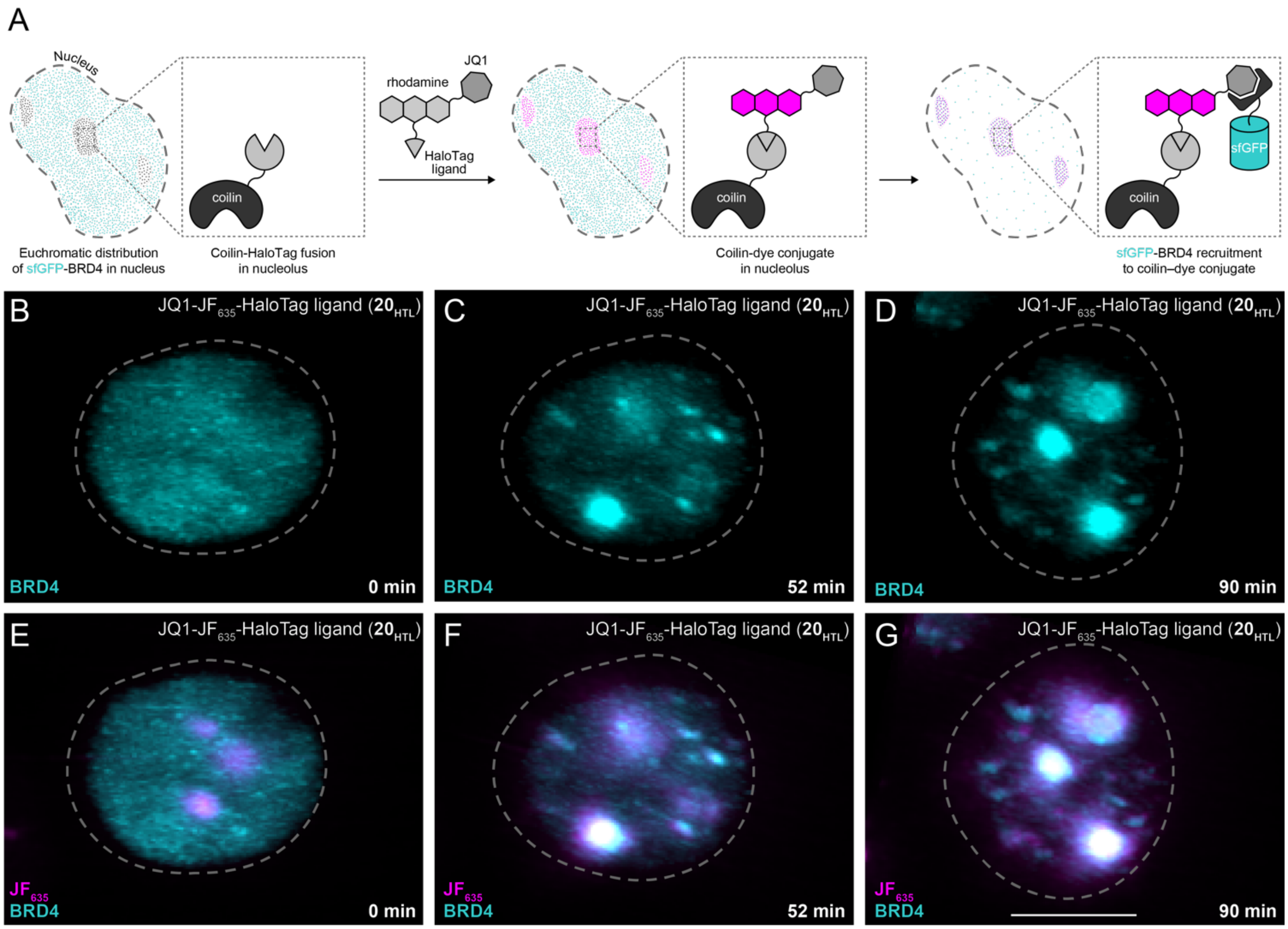


**Fig. S11.** Evaluation of BRD4 recruitment to coilin–HaloTag cells incubated with (*S*)-JQ1-JF$_{635}$-HaloTag ligand (**20**$_{\mathbf{HTL}}$). (*A*) Schematic illustrating the assay to manipulate BRD4 localization. (*B*–*G*) Maximum intensity projections from LLSM of live Neuro2a cells expressing sfGFP–BRD4 and coilin–HaloTag at time 0 min, 52 min, and 90 min after incubation with **20**$_{\mathbf{HTL}}$. (*B*–*D*) Extracted fluorescence signal from BRD4 alone. (*E*–*G*) Fluorescence signal from BRD4 and JF$_{635}$. The dashed line represents the nuclear boundary determined through histone H2B–mCherry expression. Scale bar: 5 µm.

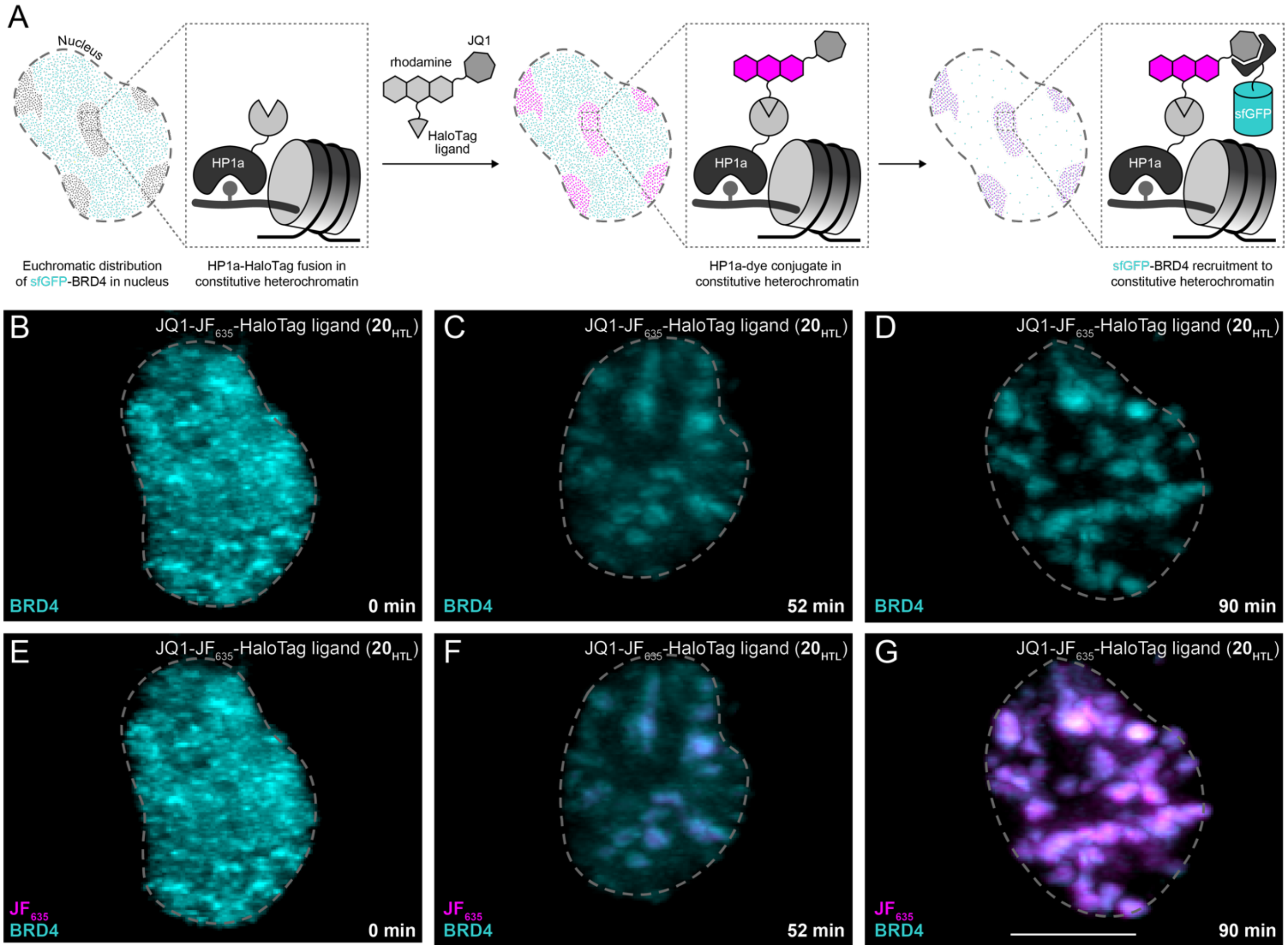


**Fig. S12.** Evaluation of BRD4 recruitment to HP1a–HaloTag cells incubated with (*S*)-JQ1-JF$_{635}$-HaloTag ligand (**20**$_{\mathbf{HTL}}$). (*A*) Schematic illustrating the assay to manipulate BRD4 localization. (*B*–*G*) Maximum intensity projections from LLSM of live Neuro2a cells expressing sfGFP–BRD4 and HP1a–HaloTag at time 0 min, 52 min, and 90 min after incubation with **20**$_{\mathbf{HTL}}$. (*B*–*D*) Extracted fluorescence signal from BRD4 alone. (*E*–*G*) Fluorescence signal from BRD4 and JF$_{635}$. The dashed line represents the nuclear boundary determined through histone H2B–mCherry expression. Scale bar: 5 µm.

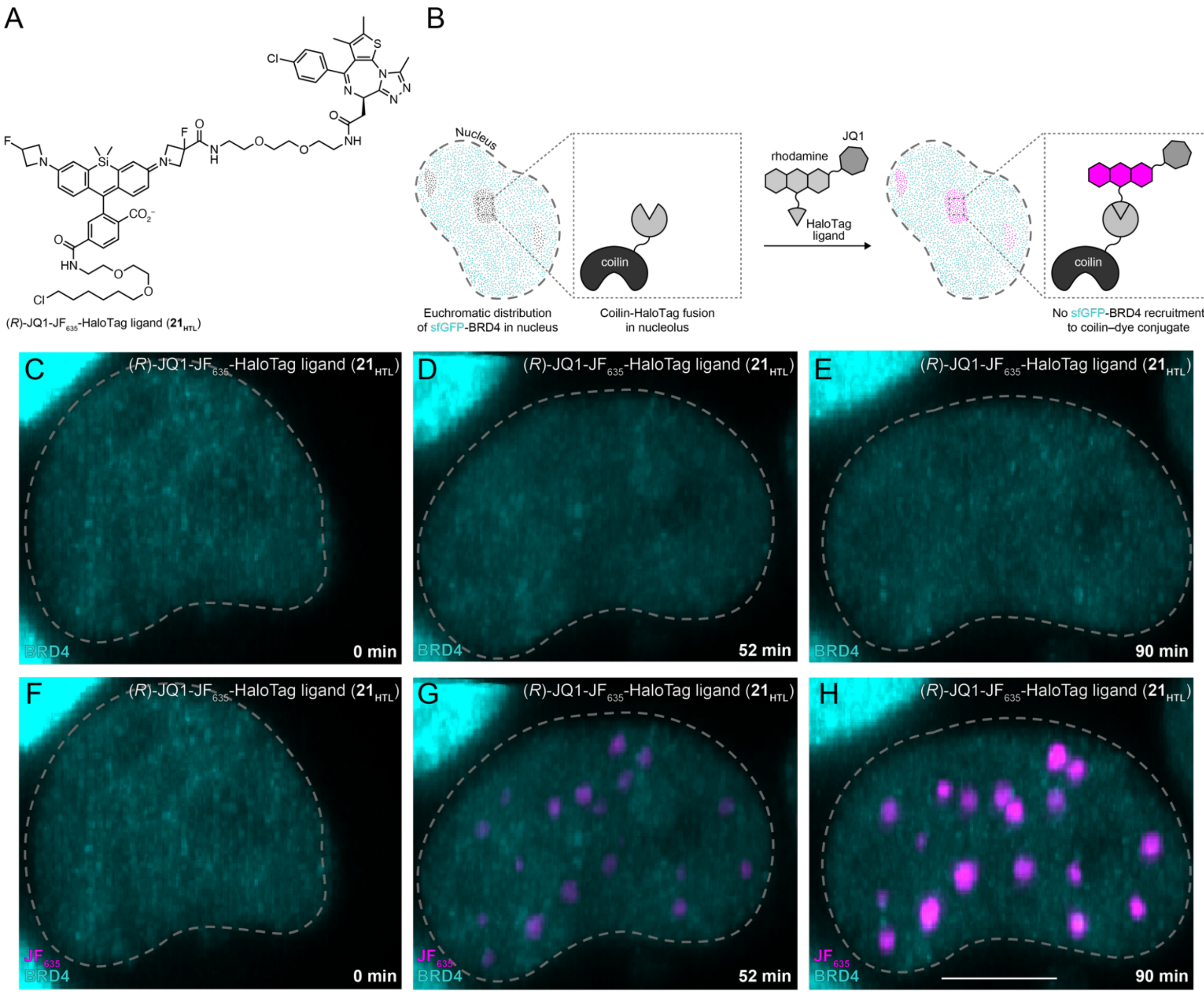


**Fig. S13.** Evaluation of BRD4 recruitment to coilin–HaloTag cells incubated with (*R*)-JQ1-JF$_{635}$-HaloTag ligand (**21**$_{\text{HTL}}$). (*A*) Chemical structure of **21**$_{\text{HTL}}$. (*B*) Schematic illustrating the assay to manipulate BRD4 localization. (*C*–*H*) Maximum intensity projections from LLSM of live Neuro2a cells expressing sfGFP–BRD4 and coilin–HaloTag at time 0 min, 52 min, and 90 min after incubation with **21**$_{\text{HTL}}$. (*C*–*E*) Extracted fluorescence signal from BRD4 alone. (*F*–*H*) Fluorescence signal from BRD4 and JF$_{635}$. The dashed line represents the nuclear boundary determined through histone H2B–mCherry expression. Scale bar: 5 µm.

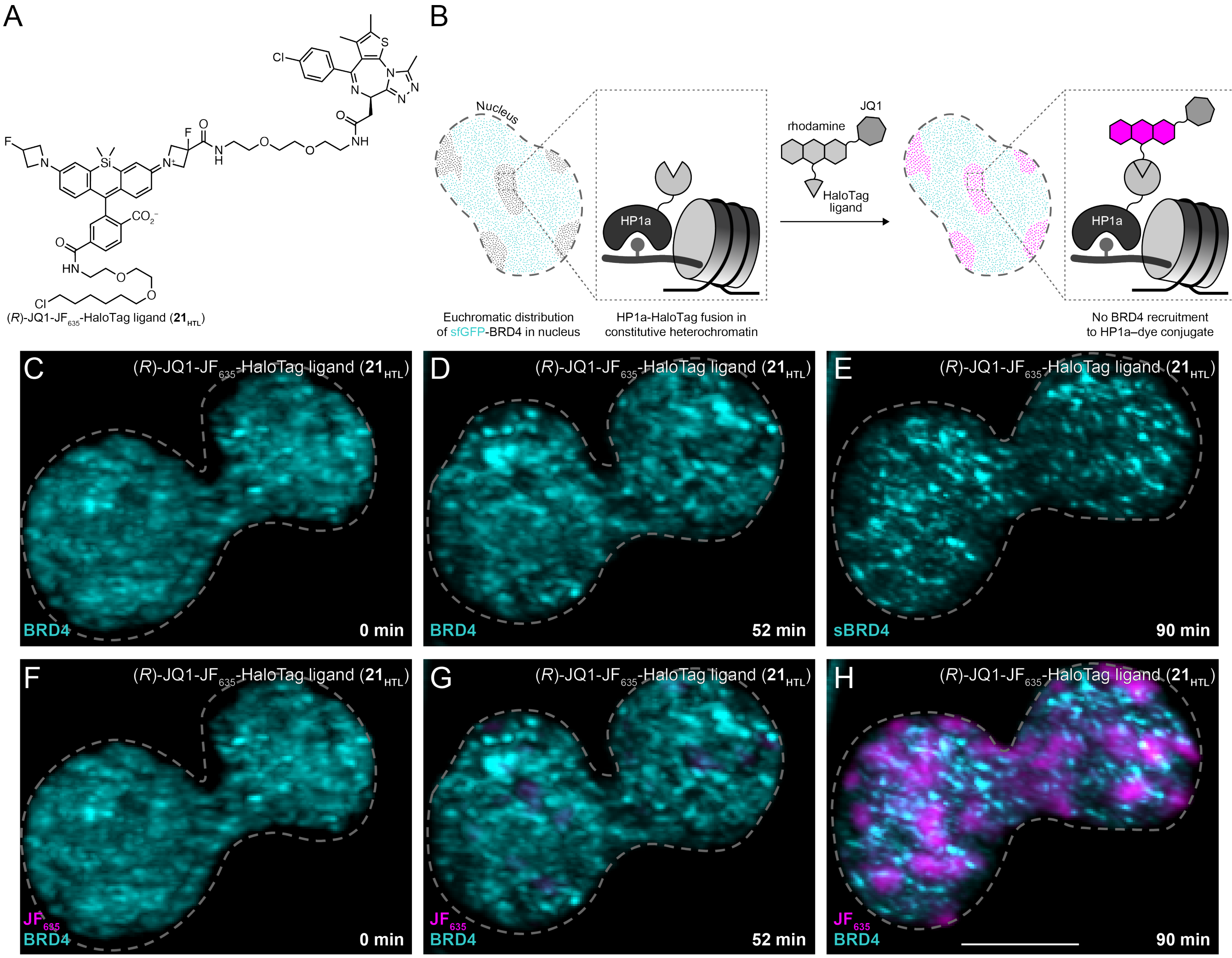


**Fig. S14.** Evaluation of BRD4 recruitment to HP1a–HaloTag cells incubated with (*R*)-JQ1-JF$_{635}$-HaloTag ligand (**21$_{HTL}$**). (*A*) Chemical structure of **21$_{HTL}$**. (*B*) Schematic illustrating the assay to manipulate BRD4 localization. (*C–H*) Maximum intensity projections from LLSM of live Neuro2a cells expressing sfGFP–BRD4 and HP1a–HaloTag at time 0 min, 52 min, and 90 min after incubation with **21$_{HTL}$**. (*C–E*) Extracted fluorescence signal from BRD4 alone. (*F–H*) Fluorescence signal from BRD4 and JF$_{635}$. The dashed line represents the nuclear boundary determined through histone H2B–mCherry expression. Scale bar: 5 µm.

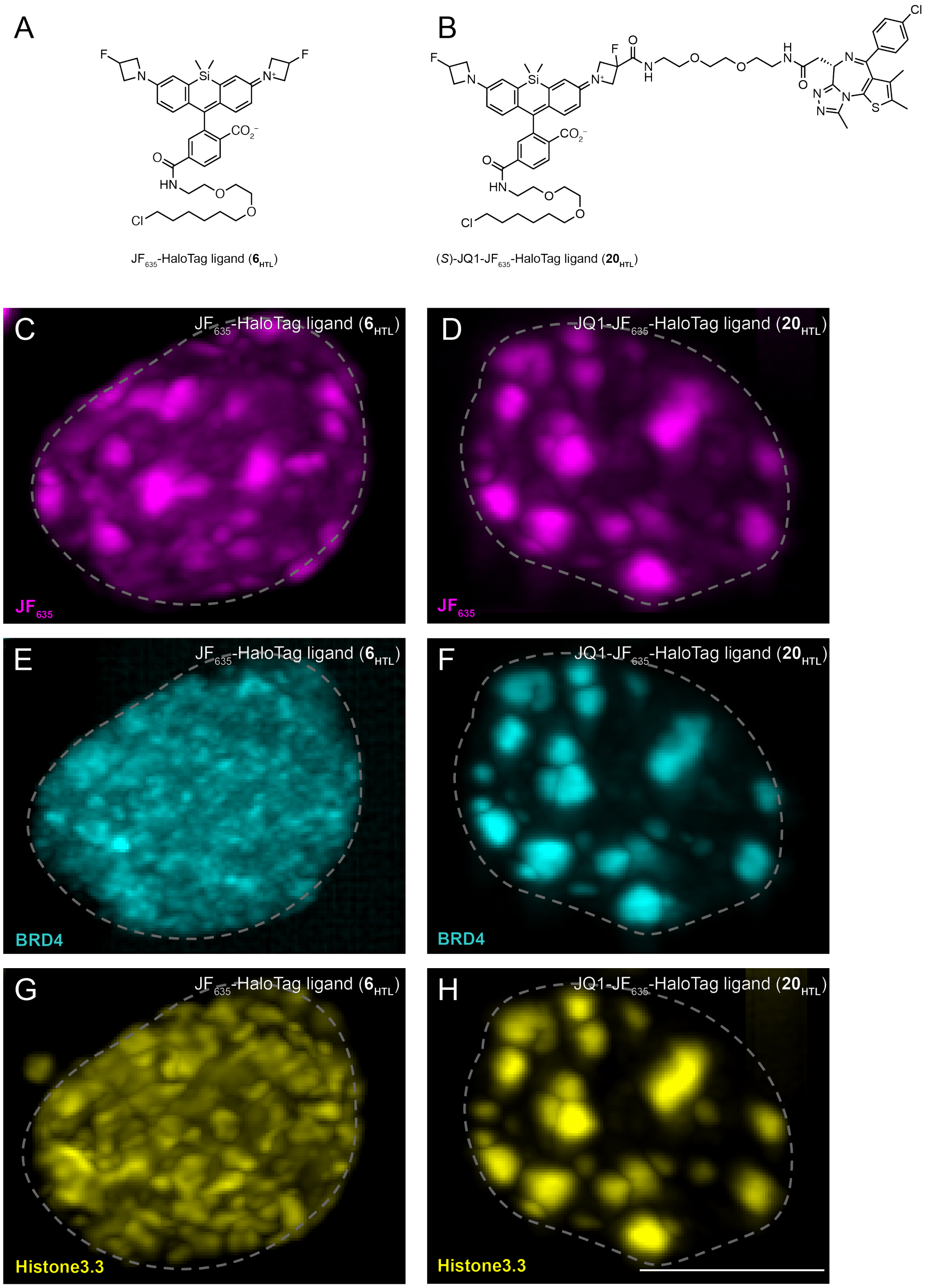


**Fig. S15.** Evaluation of Histone3.3 recruitment to HP1a–HaloTag cells incubated with $JF_{635}$-HaloTag ligand (**6**$_{HTL}$) or (*S*)-JQ1-$JF_{635}$-HaloTag ligand (**20**$_{HTL}$). (*A–B*) Chemical structures of **6**$_{HTL}$ and **20**$_{HTL}$. (*C–H*) Maximum intensity projections from LLSM of live Neuro2a cells expressing HP1a–HaloTag, sfGFP–BRD4 and Histone3.3–SNAP tag after incubation with **6**$_{HTL}$ or **20**$_{HTL}$. (*C, D*) Extracted fluorescence signal from $JF_{635}$ alone. (*E, F*) Extracted fluorescence signal from BRD4 alone. (*G, H*) Extracted fluorescence signal from Histone3.3–SNAP tag labeled with $JF_{554}$-SNAP tag ligand (**22**$_{HTL}$). The dashed line represents the nuclear boundary determined through histone H2B–mCherry expression. Scale bar: 5 μm.

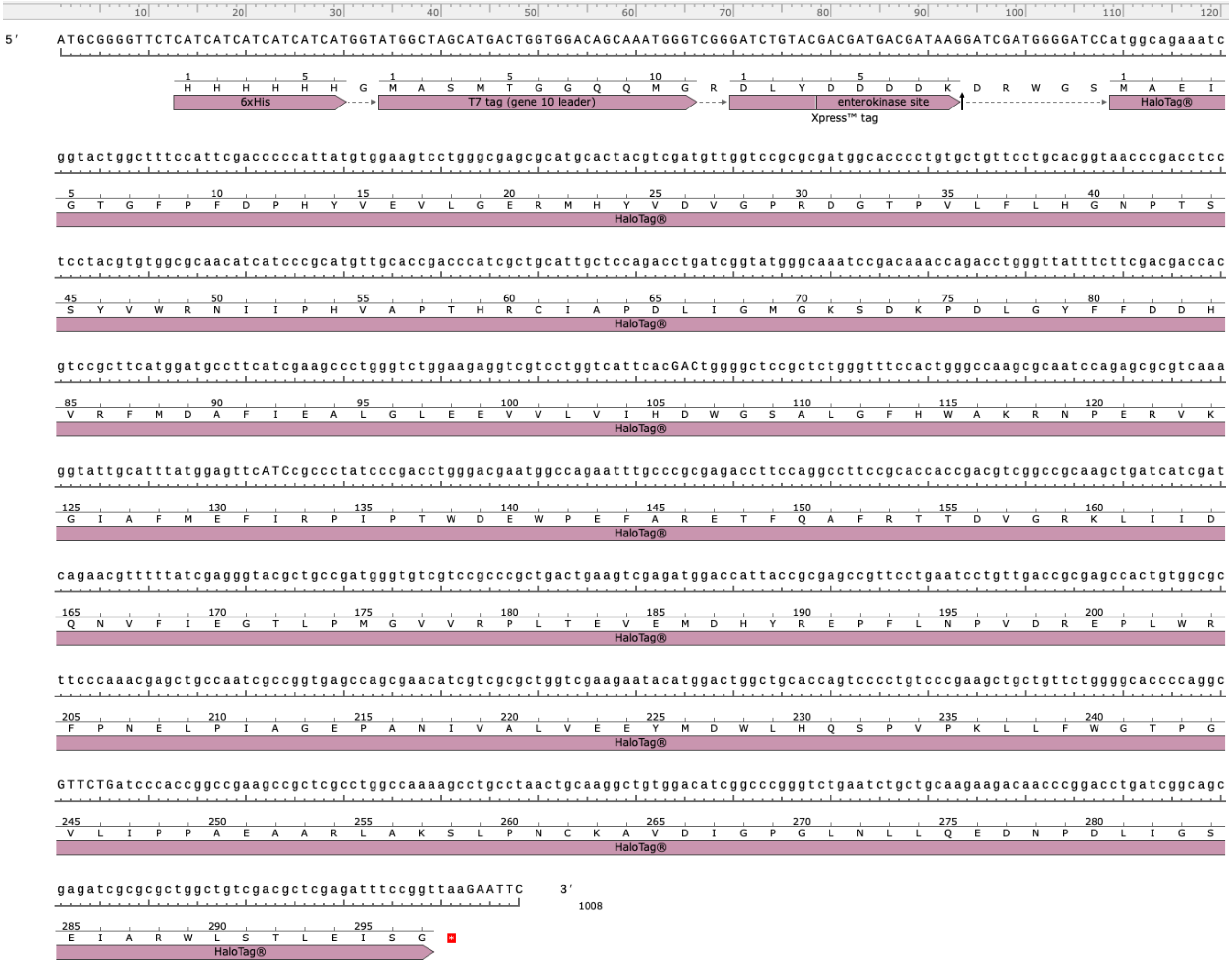


**Fig. S16.** DNA and amino acid sequences of HaloTag used for *in vitro* experiments.

**Scheme S1**

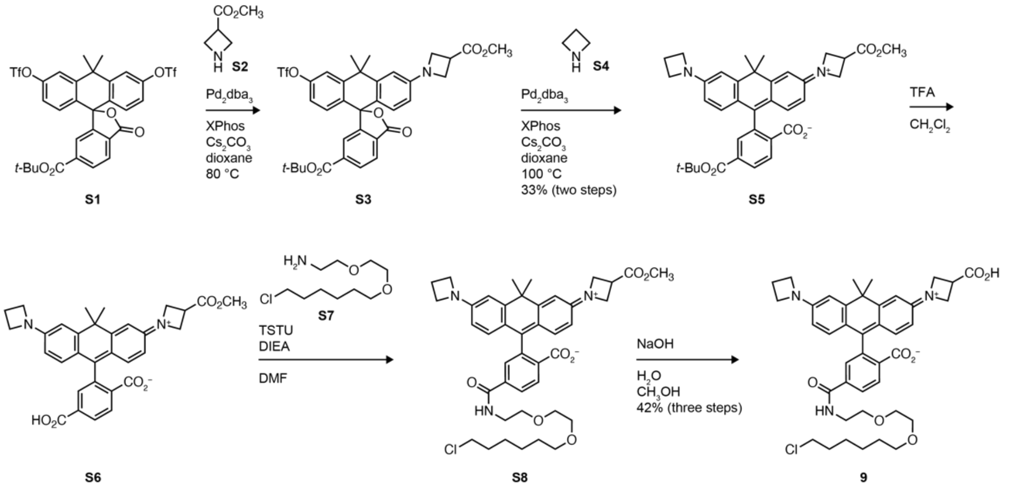


**Scheme S2**

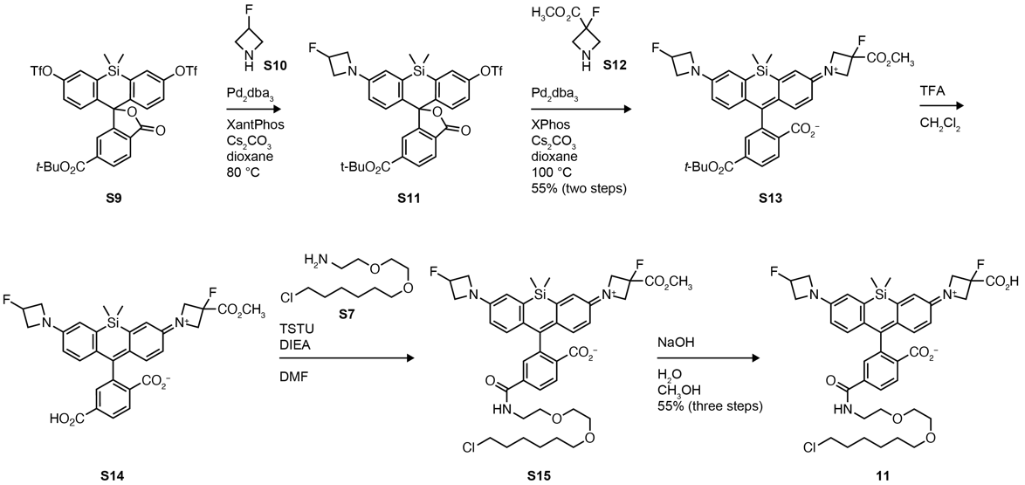

**Scheme S3**

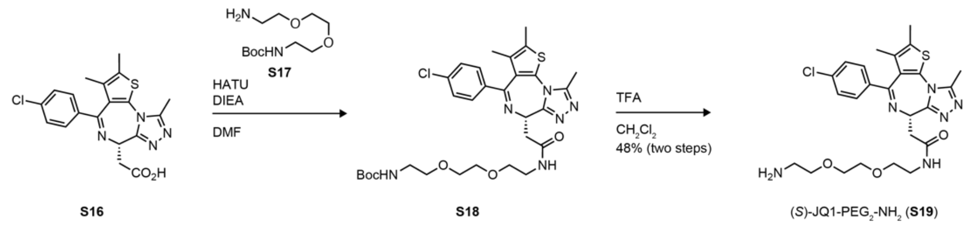


**Scheme S4**

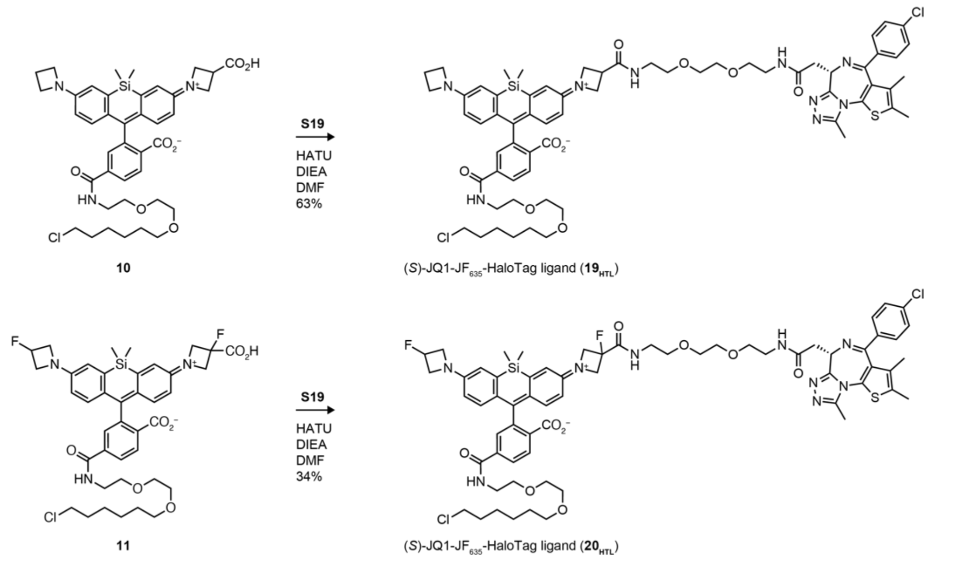


**Scheme S5**

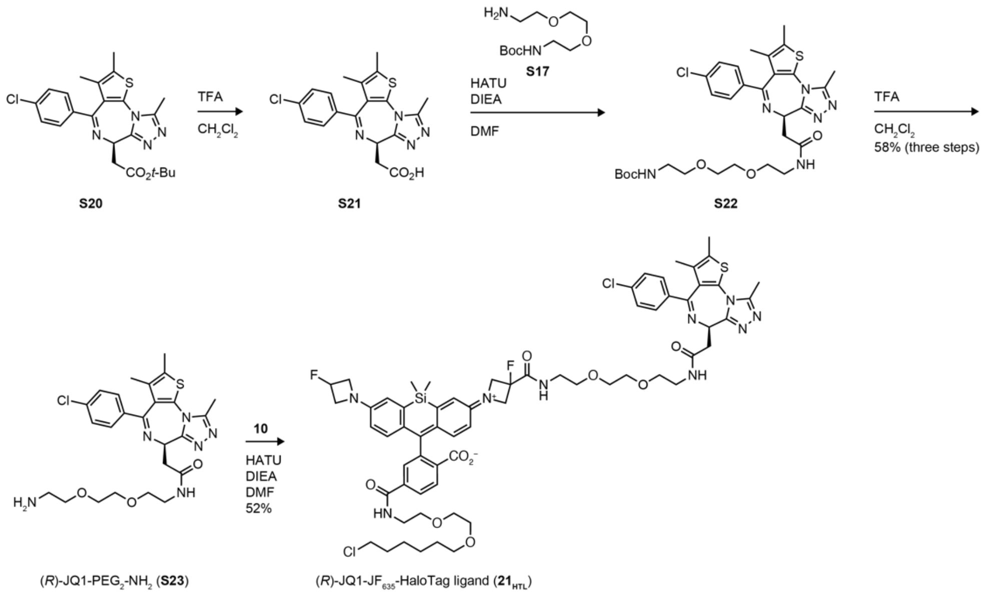

#### SPECTROSCOPY AND CELL BIOLOGY METHODS

**HaloTag protein purification.** The bacterial expression vector pRSET-A (Invitrogen) was used to recombinantly express HaloTag (HT7; Promega).[1] The soluble 6×His-Tagged HaloTag protein was affinity purified by immobilized metal affinity chromatography (IMAC) on a 5-mL Fast Flow HiTrap Sepharose 6 column (Cytiva) with a 0–200 mM imidazole elution gradient using an Avant Protein Purification System (ÄKTA). $A_{280}$ peak fractions were pooled, concentrated by a spin concentrator, and dialyzed 3× into tris-buffered saline (TBS). The amino acid sequence of HaloTag (HT7) expressed from pRSET-A is noted below. The annotated DNA and amino acid sequences are shown in **Figure S16**.

MRGSHHHHHHGMASMTGGQQMGRDLYDDDDKDRWGSMAEIGTGFPFDPHYVEVLG
ERMHYVDVGPRDGTPVLFLHGNPTSSYVWRNIIPHVAPTHRCIAPDLIGMGKSDKPDLG
YFFDDHVRFMDAFIEALGLEEVVLVIHDWGSALGFHWAKRNPERVKGIAFMEFIRPIPT
WDEWPEFARETFQAFRTTDVGRKLIIDQNVFIEGTLPMGVVRPLTEVEMDHYREPFLNP
VDREPLWRFPNELPIAGEPANIVALVEEYMDWLHQSPVPKLLFWGTPGVLIPPAEAARL
AKSLPNCKAVDIGPGLNLLQEDNPDLIGSEIARWLSTLEISG

**1-Photon spectroscopy of HaloTag ligands and HaloTag conjugates.** Compounds **$13_{HTL}$–$16_{HTL}$**, **$19_{HTL}$**, and **$20_{HTL}$** were prepared as stock solutions in DMSO and diluted such that the final DMSO concentration did not exceed 1% v/v. All measurements were taken at ambient temperature (22 ± 2 °C). Absorption spectra were recorded on a Cary Model 100 spectrometer (Agilent) using 1-cm path length 1.0-mL quartz microcuvettes from Starna Cells. Fluorescence spectra were recorded on a Cary Eclipse fluorometer (Varian) using 1-cm path length 3.5-mL quartz cuvettes (Starna Cells). As before,[2-4] HaloTag protein was used as a 100 μM solution in 1× TBS. HaloTag ligands **$13_{HTL}$–$16_{HTL}$**, **$19_{HTL}$**, and **$20_{HTL}$** (5 μM) were dissolved in 10 mM HEPES, pH 7.3, containing 0.1 mg/mL CHAPS. An aliquot of HaloTag protein (1.5 equiv) was added, and the resulting mixture was incubated until a consistent absorbance signal was observed (60–120 min). To measure the fold-increase of absorbance upon HaloTag binding, a "no HaloTag" control experiment was performed where an equivalent volume of TBS blank was added in place of the protein. Reported values for extinction coefficient ($\varepsilon$) are averages of at least two measurements.

**1-Photon spectroscopy of JQ1-rhodamine-HaloTag ligands and Bromodomain-containing protein 4 (BRD4).** Compounds **$19_{HTL}$** and **$20_{HTL}$** were prepared as stock solutions in DMSO and diluted such that the final DMSO concentration did not exceed 1% v/v. All measurements were taken at ambient temperature (22 ± 2 °C). Absorption spectra were recorded on a Cary Model 100 spectrometer (Agilent) using 1-cm path length 1.0-mL quartz microcuvettes from Starna Cells. Fluorescence spectra were recorded on a Cary Eclipse fluorometer (Varian) using 1-cm path length 3.5-mL quartz cuvettes (Starna Cells). Commercially obtained recombinant BRD4 (#31880, Active Motif), containing the catalytic core amino acids 44–168, was used as a 100 μM solution.

HaloTag ligands **19$_{HTL}$** and **20$_{HTL}$** (5 μM) were dissolved in 10 mM HEPES, pH 7.3, containing 0.1 mg/mL CHAPS. An aliquot of BRD4 protein (1.5 equiv) was added, and the absorbance signal of the resulting mixture was recorded at different time points (60–120 min). A "no BRD4" control experiment was also performed where an equivalent volume of TBS blank was added in place of the protein. Reported values for extinction coefficient ($\varepsilon$) are averages of at least two measurements.

**Determination of $K_{L-Z}$.** The lactone−zwitterion equilibrium constant ($K_{L-Z}$) was calculated as described previously[2-4] using equation 1:

$$K_{\mathrm{L\text{-}Z}} = \frac{\frac{\varepsilon_{\mathrm{dw}}}{\varepsilon_{\mathrm{max}}}}{\left(1-\frac{\varepsilon_{\mathrm{dw}}}{\varepsilon_{\mathrm{max}}}\right)} \quad (1)$$

where $\varepsilon_{dw}$ is the extinction coefficient of the dyes in a 1:1 (v/v) dioxane:water solvent mixture containing 0.01% (v/v) triethylamine; this dioxane−water mixture was chosen to give a large range of $K_{L-Z}$ values,[3] and the triethylamine additive ensures the rhodamines are in the net neutral form. The $\varepsilon_{max}$ is the maximal extinction coefficient, measured in 0.1% (v/v) trifluoroacetic acid in 2,2,2-trifluoroethanol (TFE).

**Quantum yield determination.** All reported absolute fluorescence quantum yield values ($\Phi_f$) were measured in our laboratory under identical conditions using a Quantaurus-QY spectrometer (model C11374, Hamamatsu). This instrument uses an integrating sphere to determine photons absorbed and emitted by a sample. Measurements were performed using dilute samples (absorbance < 0.1), and self-absorption corrections were performed using the instrument software.[5] Reported values are averages of at least two measurements.

**Determination of log$D_{7.4}$.** The log of distribution coefficients at pH 7.4 was determined in octanol–phosphate-buffered saline (PBS) using the miniaturized shake flask setup described previously.[6, 7] Briefly, 150 mL each of octanol and PBS pH 7.4 were stirred vigorously for 12–16 h. The layers were allowed to stand for at least 24 h for phase separation. The separated layers were collected and served as PBS saturated with octanol (PBS*) and octanol saturated with PBS (octanol*). Stock solutions (1 mM or 5 mM) of ligands were prepared in DMSO and diluted in PBS* to obtain 1 μM or 10 μM (3–5 mL) of the standard (std). This standard was used to prepare appropriate dilutions using octanol*, in triplicate, in 2-mL glass vials. The leftover standard was used to determine the standard concentration ($C_{std}$). The vials were vortexed for 1 min and shaken horizontally at 700 rpm for 2 h. The vials were then shaken upright at 150 rpm for 2 h to allow the phases to separate. The octanol* layer was pipetted out, and the PBS* layer was used for further analyses. The concentrations of ligands in the standard and PBS* ($C_{PBS*}$) were determined via fluorescence measurements of 200 μL of PBS* or std in 96-well plates on the Cytation 5 Multimode Plate Reader (BioTek). For Si-rhodamine containing ligands, 2–20 μL of trifluoroacetic acid was added to each well of the 96-well plate prior to the fluorescence measurements. Reported values are averages of at least two measurements. The $\log D_{7.4}$ was obtained as described previously using equation 2:

$$\log D_{7.4} = \log\left(\left(\frac{C_{\mathrm{std}}}{C_{\mathrm{PBS*}}} \times r - 1\right)\frac{V_{\mathrm{PBS*}}}{V_{\mathrm{octanol*}}}\right) \quad (2)$$

where $C_{\mathrm{std}}$ is the standard concentration in PBS, $C_{\mathrm{PBS*}}$ is the concentration in PBS after partitioning, $r$ is the dilution factor of the standard solution, $V_{\mathrm{PBS*}}$ is the volume of PBS used in the partitioning, and $V_{\mathrm{octanol*}}$ is the volume of octanol used in the partitioning.

**Plasmid construction for mitochondria imaging and affinity purification.** HaloTag, amplified from the pHTC HaloTag CMV-neo Vector (Promega; G7711) and monomeric-superfolder green fluorescent protein (msGFP; lab stock), was fused to the mitochondrial targeting signal from OMP25,[8] interspacing flexible GS(GSS)$_4$ linkers between each domain by PCR Splicing by Overlap Extension (SOE). This construct was subcloned into hSynapsin promoter bearing a copy of FUGW.[9] FUGW was a gift from David Baltimore (Addgene plasmid # 14883; RRID: Addgene_14883).[10] This construct was designated as pF(UG) hSyn HaloTag-TEV-4×GS-msGFP-mito.

**HEK293T cell culture.** HEK293T cells (ATCC) were passaged before 80% confluency by trypsinization (Corning; 25-053-CI) and trituration. The cell suspension was then plated on glass coverslips (Warner instruments; 64-0734, CS-18R17) coated with poly-D-lysine (Thermofisher; ICN10269491) and cultured in Dulbecco's Modified Eagle Medium (DMEM; ThermoFisher; 11965-118) supplemented with 10% v/v fetal bovine serum (FBS, Atlanta Biological) and penicillin/streptomycin (pen/strep; ThermoFisher; MT-30-001-CI) at 37 °C in a humidified 5% v/v $CO_2$ environment following ATCC guidelines. These HEK293T cells were tested for mycoplasma contamination using the Universal Mycoplasma Detection Kit (ATCC; 30-1012K) and validated using Short Tandem Repeat profiling by ATCC (ATCC; 135-XV) within the previous year or were from newly purchased stock from ATCC. HEK293T cells were transfected with pF(UG) hSyn HaloTag-TEV-4×GS-msGFP-mito using a standard calcium phosphate protocol[11] or using lipofectamine LTX reagent (ThermoFisher; 15338100). We note that this plasmid is optimized for lentivirus-mediated expression, but robust expression in HEK293T cells was observed.

**HEK293T labeling and fluorescence microscopy.** One day after HEK293T cell transfection, the GFP fluorescence signal was confirmed, and HaloTag ligand labeling experiments were conducted. For pulse–chase experiments, HEK293T cells were incubated with biotin-HaloTag ligand (**17**$_{\mathbf{HTL}}$; #G8281, Promega) at either 100 nM or 10 μM for 1 hour at 37 °C and chased with 100 nM of $JF_{549}$-HaloTag ligand (**2**$_{\mathbf{HTL}}$). We also evaluated a commercial biotin-HaloTag ligand with a longer PEG linker (**$18_{HTL}$**, #G8591, Promega) and confirmed this is not cell-permeant, as indicated in the product information[12] and from a previous publication.[13] For labeling with biotin-JF-HaloTag ligands **$13_{HTL}$–$16_{HTL}$**, cells were incubated with 100 nM or 1 μM ligand for 0.5–2 h at 37 °C. For imaging, the coverslips containing HEK293T cells were fixed with 4% paraformaldehyde in PBS at 37 °C for 10 min, washed with PBS (3×), and mounted on glass slides with ProLong Glass Antifade Mountant with NucBlue$^{TM}$ Stain (ThermoFisher; P36981). These cells were imaged using a Zeiss LSM 880 with Airyscan and a plan-apochromatic 63×/1.40 oil objective in Fast Airyscan mode. The same image acquisition settings were used for each trial and condition. Images were then processed with automatic Airyscan deconvolution settings. For quantitation of labeling intensity, ROIs were drawn surrounding individual HEK cells, and the average fluorescence ratio (JF ligand/msGFP) from multiple cells in each condition was measured; no background subtraction was necessary.

**U2OS cell culture, labeling, and fluorescence microscopy.** U2OS cells (ATCC) were cultured in Dulbecco's modified Eagle medium (DMEM, phenol red-free; Life Technologies) supplemented with 10% (v/v) fetal bovine serum (FBS, Life Technologies), 1 mM GlutaMAX (Life Technologies) and maintained at 37 °C in a humidified 5% (v/v) $CO_2$ environment. These cell lines undergo regular mycoplasma testing by the Janelia Cell Culture Facility. U2OS cells stably expressing an integrated HaloTag–histone H2B fusion protein (U2OS.H2B.HaloTag) were used for nuclear imaging and imaged 18–24 h post-plating. For other subcellular targets, U2OS cells were transiently transfected using nucleofection (Lonza) with plasmids constitutively expressing the following fusion proteins: a C-terminal transmembrane anchoring domain from platelet-derived growth factor receptor (PDGFR) fused to the HaloTag protein (HaloTag–PDGFR; for extracellular display); a HaloTag–TOMM20 fusion protein (outer mitochondrial membrane; Addgene plasmid # 123284; RRID: Addgene_123284); or HaloTag–Sec61β fusion protein (endoplasmic reticulum membrane; Addgene plasmid # 123285; RRID: Addgene_123285). The transiently transfected cells were imaged 18–24 h post-transfection. The stable and transiently transfected U2OS cells were incubated with 100 nM biotin-$JF_{549}$-HaloTag ligand (**$13_{HTL}$**), biotin-$JF_{608}$-HaloTag ligand (**$14_{HTL}$**), biotin-$JF_{646}$-HaloTag ligand (**$15_{HTL}$**), or biotin-$JF_{635}$-HaloTag ligand (**$16_{HTL}$**) for 1 h at 37 °C, washed 3× with dye-free media, then fixed with 4% paraformaldehyde in 0.1 M phosphate buffer for 15 min at 37 °C. Fixed cells were then washed 3× in 1× PBS and incubated with Hoechst 33342 (5μg/mL) for 15 min at 22 °C as a nuclear counterstain. Airyscan imaging was performed on a Zeiss LSM 980 with Airyscan 2 confocal microscope using a Plan APO 63×/1.4 oil DIC M27 objective. The same acquisition settings were used for all constructs labeled with **$13_{HTL}$–$16_{HTL}$**. These single plane images were bulk processed in ZEN Blue (Zeiss) with automatic Airyscan settings.

**Dye loading kinetics.** Live U2OS.H2B.HaloTag stable cells were labeled over a time course of 0–4 h with 200 nM of biotin-$JF_{549}$-HaloTag ligand (**$13_{HTL}$**), biotin-$JF_{608}$-HaloTag ligand (**$14_{HTL}$**), biotin-$JF_{646}$-HaloTag ligand (**$15_{HTL}$**), or biotin-$JF_{635}$-HaloTag ligand (**$16_{HTL}$**) at 37 °C. Cells were then washed 3× with dye-free media, fixed with 4% paraformaldehyde in 0.1 M phosphate buffer, pH 7.4 for 15 min at 37 °C. Confocal imaging was performed on a Leica SP8 with an HC PL APO CS2 20×/0.75 immersion objective using the tunable white light laser (WLL) to excite dyes at their $\lambda_{abs}$ in constant power mode. Fluorescence was quantified as the average integrated density of background corrected nuclear signals from confocal image stack projections analyzed in FIJI; $n$ = 100 nuclear signals per compound.

**Affinity capture and in-gel fluorescence quantification.** Affinity capture pulldown isolation experiments of biotin-labeled HaloTag fusion proteins were performed on ice-cold isolation buffer (IB) consisting of KCl (0.18 M), ethylene glycol-bis(β-aminoethyl ether)-*N*,*N*,*N′*,*N′*-tetraacetic acid (EGTA; 1 mM), 3-(*N*-morpholino)propanesulfonic acid (MOPS; 5 mM); the pH was adjusted to pH = 7.35 using KOH(aq).[14] StrepTactin Microbeads (IBA Lifesciences, 6-5510-050) were used as capture reagents as these beads had lowest amount of background observed in our hands. These beads were kept at 4 °C and blocked in IB buffer with PMSF and 1% bovine serum albumin (BSA; Jackson ImmunoResearch, 001-000-162) with rotation at 4 °C for 30 min. HEK293T cells were scraped with 200 µL IB and homogenized using 12 strokes with a 27G syringe. The resulting homogenate was spun at 4 °C at 800g for 5 min to pellet large cellular debris. 50 µL of freshly blocked StrepTactin microbeads were added to the supernatant and incubated for 30 min at 4 °C with slow rotation. After incubation, tubes were placed in DynaMag2 (ThermoFisher, 12321D) magnetic racks and washed twice with IB+PMSF. All wash steps were kept at 4 °C (cold room). The post-bead supernatant and wash volumes were pooled and spun at 15,000g for 12 min. This last step pelleted the unbound mitochondria for final analysis of capture efficiency. The bead fraction and the pooled/pelleted fraction were then solubilized using the lysis buffer containing 1× PBS, 2% SDS, 1% Triton x-100, 10 mM EDTA, and protease inhibitors (2 mM PMSF, aprotinin, leupeptin, and pepstatin A). Samples were then incubated at 50 °C for 3–4 min after the addition of Laemmli sample buffer (SB) (Bio-Rad, 1610747) with B-mercaptoethanol (Bio-Rad, 1610710). For protein detection, 10–20 µL of the sample was subjected to SDS-PAGE, using 10% Tris-glycine gels (TGX) (Bio-Rad, 5671034) with Tris-Glycine buffer (Bio-Rad, 1610732). The affinity capture pulldown efficiency was monitored using in-gel fluorescence. The msGFP signal from HaloTag-TEV-4×GS-msGFP-mito after cell lysis and SDS-PAGE was analyzed, in-gel, using a Bio-Rad Chemidoc MP imager (Bio-Rad) through GFP fluorescence excitation and emission filters. Data were quantified by densitometry using Fiji.[15] Capture efficiency was calculated comparing the total GFP signal recovered from StrepTactin beads vs the signal recovered from the high speed spin of the bead supernatant and pooled wash volumes.

**Quantification of HaloTag labeling efficiency using in-gel fluorescence.** HEK293T cells were cultured, incubated and transfected with pF(UG) hSyn HaloTag-TEV-4×GS-msGFP-mito using lipofectamine LTX reagent as described above. To examine saturation of HaloTag, we screened for fluorescence plateau using 1 µM of $JF_{479}$-HaloTag ligand, $JF_{549}$-HaloTag ligand and $JF_{608}$-HaloTag ligand at 15, 30, 60, 90, 120, and 150 minutes. Only $JF_{549}$-HaloTag ligand and $JF_{608}$-

HaloTag ligand reached a fluorescence plateau, and they were chosen as saturating counter stains or 'chase' dyes for examining the labeling efficiency of biotin-JF-HaloTag ligands (**$13_{HTL}$–$16_{HTL}$**). Next, we developed an in-gel assay to independently quantitate labeling efficiencies of **$13_{HTL}$–$16_{HTL}$**. We used $JF_{549}$-HaloTag ligand as chase for evaluating biotin-$JF_{608}$-HaloTag ligands (**$14_{HTL}$**), biotin-$JF_{646}$-HaloTag ligands (**$15_{HTL}$**), and biotin-$JF_{635}$-HaloTag ligands (**$16_{HTL}$**). We used $JF_{608}$-HaloTag ligand as chase for evaluating biotin-$JF_{549}$-HaloTag ligands (**$13_{HTL}$**). Each experiment consisted of testing the biotin variants **$13_{HTL}$–$16_{HTL}$** at 100 nM and 1 μM with an incubation time of 2 h and a chase labeling incubation of 1 μM for 0.5 h. The 0.5 h chase labeling period was included with the 2 h total incubation. Each experiment consisted of untransfected cells, A) transduced, but no dye, B) non-biotinylated variant of primary label at 1 μM for 0.5 h, C) non-biotinylated version of the chase JF dye at 1 μM for 0.5 h, D) 100 nM of the non-biotinylated version of the primary label for 2 h, E) 100 nM of the biotinylated version of primary label for 2 h, F) 1 μM of the non-biotinylated version of the primary label for 2 h, G) 1 μM of the biotinylated version of primary label for 2 h, and H) 1 μM of the non-biotinylated version of the chase JF dye at 1 μM for 2 h (same as C but for 2 h vs 0.5 h; independent verification of chase saturation using 0.5 h incubation). Samples D-G were incubated with a non-biotinylated chase dye at 1 μM for 0.5 h. Cells were washed and protein lysates were harvested using Bio-Rad SDS-PAGE products as described above. Specifically, cells were lysed using protein lysis buffer as described above, and adding XT Sample Buffer (Bio-Rad, 1610791) and XT Reducing Agent (Bio-Rad, 1610792). Samples were heated at 72 ºC for 5 mins. Samples were then loaded onto a 4–12% Criterion XT Bis-Tris gel (Bio-Rad, 3450124), and ran at 150 V for approximately 1 h. Gels were then imaged on a Chemidoc MP imager (Bio-Rad) using orange, red, and far red filters. Bands representing HaloTag-ligand were quantified by densitometry using Fiji.[15] These data were collected and analyzed in MS Excel and Graphpad Prism. For labeling efficiency calculations, bands representing saturating chase dye fluorescence (*e.g.*, $JF_{549}$) were compared to saturating primary dye fluorescence. This workflow will reveal the approximate amount of HaloTag that did not react to a primary dye during their initial incubation.

The equation is as follows: 1 - ($JF_{549}$ chase dye fluorescence / $JF_{549}$ primary dye fluorescence), where chase dye fluorescence is after incubation with a red JF-biotin dye, and primary dye fluorescence means cells were incubated with $JF_{549}$ only. Both incubations occurring at a saturating concentration.

**Neuro2a cell culture, labeling, and lattice light sheet microscopy (LLSM)**. Neuro2a cells (ATCC, CCL-131) were cultured in Eagle's Minimum Essential Medium (MEM, phenol red-free; Life Technologies) supplemented with 10% (v/v) fetal bovine serum (FBS), and maintained at 37 °C in a humidified 5% (v/v) $CO_2$ environment. Cells were transiently transfected using nucleofection (Lonza) with plasmids constitutively expressing Histone2B–mCherry, BRD4–sfEGFP, and either Coilin–HaloTag or Heterochromatin Protein 1a (HP1a)–HaloTag. In some cases, H2B–mCherry was replaced by Histone 3.3 to assess the locations of transcriptionally active chromatin. All inserts cDNAs were synthesized by Genscript and subcloned into the EcoRI and

NotI sites of the pCIG2 expression vector. 18–24 h post-transfection, the transiently transfected cells seeded on 5-mm coverslips were mounted in custom-fabricated sample holders and loaded on an LLSM built by Intelligent Imaging Innovations. Prior to imaging, the medium in the LLSM sample chamber was equilibrated with 8 nM (*i.e.*, 1/10 of the IC50 of free (*S*)-JQ1) or 20 nM 8 nM (*i.e.*, 1/4 of the IC50 of free (*S*)-JQ1) $\mathbf{19_{HTL}}$, $\mathbf{20_{HTL}}$, or $\mathbf{21_{HTL}}$, and full 3D volumes at multiple stage positions were imaged at 2 minute intervals for 90 minutes total.

**LLSM imaging processing.** Raw image data volumes were deskewed and rotated, then empirically deconvolved using a Richardson-Lucy maximum likelihood algorithm using experimentally measured PSFs for each color channel for 30 iterations. Channel alignment corrections were performed using Advanced Normalization Tools R (ANTsR) registration run in "translation" mode using 2σ Gaussian-blurred versions of the images, and then transforms were applied to the deconvolved images. Next, whole nuclei were segmented using thresholds greater than 2–3x the noise floor. These segmentation masks were then used to isolate each nucleus for further quantitative analyses. For instance, HaloTag labeled domains were next defined by taking the signal intensity greater than 2x the median of the total signal intensity of HP1a within each nucleus. The amount of BRD4 signal was then assessed in- versus outside Halo-tag labeled domains throughout the 90-minute time-lapse acquisition. For visualization improvements of the time-lapse volumes, we used the photobleach correction ImageJ plugin run in histogram matching mode to correct photobleaching of the H2B–mCherry channel. In addition, we used ANTsR to register all subsequent timepoints of each nuclei to its initial timepoint position.

**Data availability.** Other data supporting this study's findings are available from the corresponding author upon request.

#### GENERAL SYNTHETIC ORGANIC CHEMISTRY METHODS

Commercial reagents were the highest quality available and used as received. Solvents for reactions were of anhydrous grade, purchased in septum-sealed bottles, and stored under an inert atmosphere. Reactions were conducted in round-bottomed flasks or septum-sealed crimp-top microwave reaction vials (Biotage) containing Teflon-coated magnetic stir bars. All reactions were conducted under an inert atmosphere of Ar(g) and protected from light using Al foil unless otherwise noted. Heating of reaction mixtures was achieved through aluminum blocks on top of a stirring hotplate equipped with an electronic contact thermometer. Reactions were monitored either by thin layer chromatography (TLC) on precoated TLC glass plates (silica gel 60 F254, 250 µm thickness) or by tandem liquid chromatography–mass spectrometry (LC–MS; Shimadzu LCMS 2020, Phenomenex Kinetex 30 × 2.1 mm 2.6 µm C18 column, 1–10 µL injection, 5–98% $CH_3CN/H_2O$ linear gradient with constant 0.1% v/v $HCO_2H$, 6 min run, 1 mL/min flowrate, ESI, positive ion mode). TLC plates were visualized either by UV illumination or by developing the TLC with ceric ammonium molybdate or $KMnO_4$.

Reaction products were purified either by flash chromatography on Biotage Isolera automated purification system using prepacked silica gel columns and/or by preparative high-pressure liquid chromatography (HPLC; Agilent 1200, Phenomenex Gemini–NX 150 × 30 mm 10 µm C18 110 Å column, 42 mL/min flowrate) under the indicated solvent gradient conditions. Analytical HPLC analyses were performed on a LC-MS system (Agilent 1200, Phenomenex Gemini–NX 150 × 4.6 mm 5 µm C18 110 Å column, 1 mL/min flowrate) or an analytical HPLC (Shimadzu UFLC, Phenomenex Gemini–NX 150 × 4.6 mm 5 µm C18 110 Å column, 1 mL/min flowrate) under the indicated conditions. All the HPLC systems are fitted with a diode array detector. High-resolution mass spectrometry was obtained from the High Resolution Mass Spectrometry Facility at the University of Iowa. NMR spectra were recorded on Bruker Avance 400 MHz spectrometer and processed through MestReNova. Deuterated solvents were used as purchased. $^{1}H$ and $^{13}C$ chemical shifts (δ) were referenced to TMS or residual solvent peaks. $^{19}F$ chemical shifts were referenced to $CFCl_3$. Data for $^{1}H$ NMR spectra are reported as follows: chemical shift (δ ppm), multiplicity (s = singlet, d = doublet, t = triplet, q = quartet, p = pentet (quintet), dd = doublet of doublets, dt = doublet of triplets, m = multiplet, br = broad signal), coupling constant (Hz), and integration. Data for $^{13}C$ NMR spectra are reported by chemical shift (δ ppm) with hydrogen multiplicity (C, CH, $CH_2$, $CH_3$) information obtained from DEPT spectra.

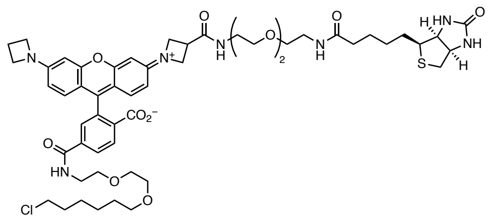

**Biotin-JF$_{549}$-HaloTag ligand (13$_{HTL}$).** The TFA salt of 3″-carboxy-JF$_{549}$ -HaloTag ligand[16] (**8**; 10.0 mg, 8.5 µmol, 1 equiv) was dissolved in anhydrous DMF (2 mL). To this solution were added $Et_3N$ (12.0 µL, 85 µmol, 10 equiv), DSC (5.5 mg, 21 µmol, 2.5 equiv), and a catalytic amount of DMAP (~0.05 mg). The reaction mixture was stirred for 90 min at ambient temperature, after which biotin-PEG$_2$-NH$_2$ (**12**, 32.0 mg, 85 µmol, 10 equiv) was added. The reaction mixture was further stirred for 16 h at ambient temperature. The solvent was removed under reduced pressure, and the product was purified by preparative HPLC using a 5–95% $CH_3CN/H_2O$ linear gradient with constant 0.1% v/v trifluoroacetic acid (TFA). Product-containing fractions were combined and lyophilized to obtain **13$_{HTL}$** as a red solid (TFA salt, 4.5 mg, 51%). $^1$H NMR ($CD_3OD$, 400 MHz) δ 8.39 (d, *J* = 8.2 Hz, 1H), 8.21 (dd, *J* = 8.2, 1.8 Hz, 1H), 7.81 (d, *J* = 1.8 Hz, 1H), 7.08 (d, *J* = 9.1 Hz, 2H), 6.63 (ddd, *J* = 9.2, 5.3, 2.1 Hz, 2H), 6.59 (d, *J* = 2.1 Hz, 1H), 6.53 (d, *J* = 2.1 Hz, 1H), 4.48 – 4.39 (m, 3H), 4.38 – 4.23 (m, 7H), 3.74 – 3.51 (m, 19H), 3.48 – 3.41 (m, 4H), 3.36 (t, *J* = 5.5 Hz, 2H), 3.22 – 3.14 (m, 1H), 2.89 (dd, *J* = 12.8, 5.0 Hz, 1H), 2.66 (dd, *J* = 12.8, 3.0 Hz, 1H), 2.56 (p, *J* = 7.4 Hz, 2H), 2.21 (t, *J* = 7.4 Hz, 2H), 1.77 – 1.55 (m, 6H), 1.53–1.47 (m, 2H), 1.45 – 1.30 (m, 6H). $^{13}$C NMR ($CD_3OD$, 101 MHz) δ 176.12 (C), 173.82 (C), 167.92 (C), 167.34 (C), 166.03 (C), 160.59 (C), 158.93 (C), 158.63 (C), 158.18 (C), 157.68 (C), 139.40 (C), 135.58 (C), 134.75 (C), 132.84 (CH), 132.44 (CH), 132.33 (CH), 130.37 (CH), 130.05 (CH), 115.15 (C), 114.96 (C), 113.94 (CH), 113.49 (CH), 95.61 (CH), 95.18 (CH), 72.14 ($CH_2$), 71.31 ($CH_2$), 71.24 ($CH_2$), 71.14 ($CH_2$), 70.61 ($CH_2$), 70.46 ($CH_2$), 70.35 ($CH_2$), 63.33 (CH), 61.61 (CH), 56.97 (CH), 55.32 ($CH_2$), 52.98 ($CH_2$), 45.75 ($CH_2$), 41.21 ($CH_2$), 41.06 ($CH_2$), 40.63 ($CH_2$), 40.28 ($CH_2$), 36.74 ($CH_2$), 34.62 (CH), 33.72 ($CH_2$), 30.46 ($CH_2$), 29.76 ($CH_2$), 29.50 ($CH_2$), 27.69 ($CH_2$), 26.84 ($CH_2$), 26.44 ($CH_2$), 16.81 ($CH_2$). $^{19}$F NMR ($CD_3OD$, 376 MHz) δ −75.35. Analytical HPLC: $t_R$= 11.2 min, 98.0% purity (10–95% MeCN/$H_2O$ linear gradient over 20 min with constant 0.1% v/v TFA, 1 mL/min flow rate, detection at 550 nm). HRMS (ESI) calculated for $C_{54}H_{71}N_7O_{11}SCl$ $[M+H]^+$ = 1060.4615, found 1060.4623.

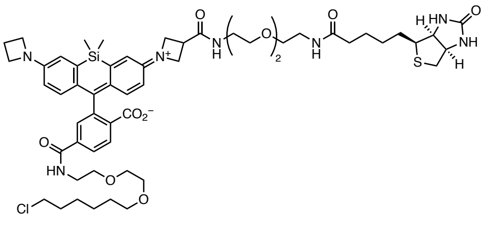

**Biotin-JF$_{646}$-HaloTag ligand (15$_{HTL}$).** The TFA salt of 3″-carboxy-JF$_{646}$ -HaloTag ligand[16] (**10**; 57.0 mg, 66 μmol, 1 equiv) was dissolved in DMF (2 mL). To this solution were added $Et_3N$ (96.0 μL, 661 μmol, 10 equiv), DSC (42.3 mg, 165.2 μmol, 2.5 equiv), and a catalytic amount of DMAP (~0.05 mg). The reaction mixture was stirred for 90 min at ambient temperature, after which biotin-PEG$_2$-NH$_2$ (**12**, 99.0 mg, 264 μmol, 4 equiv) was added. The reaction mixture was stirred overnight at ambient temperature. The solvent was removed under reduced pressure, and the product was purified by preparative HPLC using a 30–50% $CH_3CN/H_2O$ linear gradient with a constant 0.1% v/v TFA. Product-containing fractions were combined and lyophilized to obtain **15$_{HTL}$** as a blue solid (TFA salt, 43 mg, 58.1%). $^1$H NMR ($CD_3OD$, 400 MHz) δ 8.24 (d, $J$ = 8.2 Hz, 1H), 8.10 (dd, $J$ = 8.2, 1.7 Hz, 1H), 7.69 (d, $J$ = 1.6 Hz, 1H), 6.92 (d, $J$ = 2.5 Hz, 2H), 6.88 (dd, $J$ = 9.2, 7.9 Hz, 2H), 6.35 (dd, $J$ = 9.2, 2.6 Hz, 2H), 4.47 – 4.42 (m, 1H), 4.41 – 4.18 (m, 9H), 3.67 – 3.49 (m, 19H), 3.45 – 3.40 (m, 4H), 3.35 (t, $J$ = 5.5 Hz, 2H), 3.20 – 3.12 (m, 1H), 2.88 (dd, $J$ = 12.8, 5.0 Hz, 1H), 2.66 (d, $J$ = 12.7 Hz, 1H), 2.51 (p, $J$ = 7.6 Hz, 2H), 2.20 (t, $J$ = 7.4 Hz, 2H), 1.76 – 1.55 (m, 6H), 1.53 – 1.47 (m, 2H), 1.46 – 1.29 (m, 6H), 0.60 (s, 3H), 0.55 (s, 3H). $^{13}$C NMR ($CD_3OD$, 400 MHz) δ 176.11 (C), 174.04 (C), 168.44 (C), 168.22 (C), 166.04 (C), 154.06 (C), 153.57 (C), 139.38 (C), 133.58 (C), 131.20 (CH), 130.41 (CH), 130.25 (CH), 129.00 (CH), 119.57 (CH), 118.98 (CH), 113.39 (CH), 113.08 (CH), 72.12 (CH$_2$), 71.30 (CH$_2$), 71.20 (CH$_2$), 71.13 (CH$_2$), 70.61 (CH$_2$), 70.47 (CH$_2$), 70.36 (CH$_2$), 63.34 (CH), 61.60 (CH), 56.97 (CH), 55.39 (CH$_2$), 53.32 (CH$_2$), 45.74 (CH$_2$), 41.14 (CH$_2$), 41.05 (CH$_2$), 40.59 (CH$_2$), 40.27 (CH$_2$), 36.74 (CH$_2$), 34.90 (CH), 33.71 (CH$_2$), 30.44 (CH$_2$), 29.75 (CH$_2$), 29.49 (CH$_2$), 27.68 (CH$_2$), 26.83 (CH$_2$), 26.42 (CH$_2$), 17.04 (CH$_2$), -0.74 (CH$_3$), -1.66 (CH$_3$). Five aromatic quaternary carbons were not observed. $^{19}$F NMR ($CD_3OD$, 376 MHz) δ −75.35. Analytical HPLC: $t_R$= 11.9 min, 97.0% purity (10–95% MeCN/$H_2O$ linear gradient over 20 min with constant 0.1% v/v TFA, 1 mL/min flow rate, detection at 254 nm). HRMS (ESI) calculated for $C_{56}H_{76}N_7O_{10}SClSiNa$ [M+Na]$^+$ = 1124.4724**,** found 1124.4730.

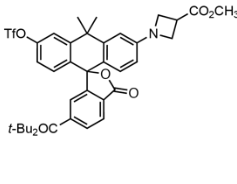

**methyl 1-(6'-(*tert*-butoxycarbonyl)-10,10-dimethyl-3'-oxo-3-(((trifluoromethyl)sulfonyl)oxy) -3'*H*,10*H*-spiro[anthracene-9,1'-isobenzofuran]-6-yl)azetidine-3-carboxylate** (S3)**.** An oven-dried microwave reaction vial was charged with 6-*tert*-butoxycarbonyl carbofluorescein ditriflate[2] (**S1**, 722.6 mg, 1 mmol, 1 equiv), $Pd_2dba_3$ (46 mg, 50 μmol, 0.05 equiv), XPhos (72 mg, 150 μmol,

0.15 equiv), methyl azetidine-3-carboxylate hydrochloride (**S2**, 182 mg, 1.2 mmol, 1.2 equiv), and $Cs_2CO_3$ (912 mg, 2.8 mmol, 2.8 equiv). The vial was sealed and backfilled with Ar(g) (3×), after which anhydrous dioxane (10 mL) was added. The resulting mixture was stirred at 80 °C for 3 h. The reaction was cooled and filtered through a pad of celite using EtOAc. The filtrate was concentrated under reduced pressure. Purification by $SiO_2$ gel chromatography (50 g $SiO_2$ column, 0–50% EtOAc/hexanes, linear gradient) provided monoazetidine compound **S3** as a light green foamy solid (220 mg) and unreacted starting material (168 mg). LC–MS (ESI): calculated for $C_{34}H_{33}F_3NO_9S$ $[M+H]^+$ = 688.18, found 688.18. This material was used immediately in the subsequent synthetic step.

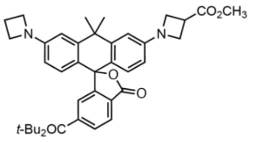


**methyl 1-(3-(azetidin-1-yl)-6'-(*tert*-butoxycarbonyl)-10,10-dimethyl-3'-oxo-3'*H*,10*H*-spiro [anthracene-9,1'-isobenzofuran]-6-yl)azetidine-3-carboxylate (*i.e.*, 3″-methoxycarbonyl-6-*tert*-butoxycarbonyl-JF$_{608}$; S5).** An oven-dried microwave reaction vial was charged with **S3** (220 mg, 320 µmol, 1 equiv), $Pd_2dba_3$ (29 mg, 32 µmol, 0.1 equiv), XPhos (46 mg, 96 µmol, 0.3 equiv), azetidine (**S4**, 108 µL, 91 mg, 1.6 mmol, 5 equiv), and $Cs_2CO_3$ (292 mg, 896 µmol, 2.8 equiv). The vial was sealed and backfilled with Ar(g) (3×), after which anhydrous dioxane (10 mL) was added. The resulting mixture was stirred at 100 °C for 4 h, then cooled to room temperature and filtered through a pad of celite using EtOAc. The filtrate was concentrated under reduced pressure. Purification by $SiO_2$ gel chromatography (50 g $SiO_2$ column, 0–100% EtOAc/hexanes, linear gradient) provided **S5** as a green-blue solid (153 mg, 33.4% over two steps). $^1H$ NMR ($CDCl_3$, 400 MHz) δ 8.15 (dd, *J* = 8.0, 1.3 Hz, 1H), 8.01 (d, *J* = 8.0 Hz, 1H), 7.61 (t, *J* = 1.0 Hz, 1H), 6.63 – 6.51 (m, 4H), 6.23 (ddd, *J* = 8.8, 5.1, 2.3 Hz, 2H), 4.15 – 4.03 (m, 4H), 3.92 (t, *J* = 7.2 Hz, 4H), 3.75 (s, 3H), 3.58 (tt, *J* = 8.6, 6.1 Hz, 1H), 2.38 (p, *J* = 7.2 Hz, 2H), 1.83 (s, 3H), 1.73 (s, 3H), 1.53 (s, 9H). $^{13}C$ NMR (101 MHz, $CDCl_3$) δ 173.21 (C), 170.08 (C), 164.51 (C), 155.51 (C), 152.37 (C), 151.63 (C), 146.92 (C), 146.67 (C), 137.85 (C), 130.17 (CH), 130.13 (C), 128.99 (CH), 128.91 (CH), 125.05 (CH), 124.87 (CH), 120.85 (C), 119.77 (C), 110.77 (CH), 110.62 (CH), 108.43 (CH), 108.05 (CH), 82.36 (C), 54.55 ($CH_2$), 54.52 ($CH_2$), 52.38 ($CH_3$), 52.36 ($CH_2$), 38.49 (C), 35.43 ($CH_3$), 33.54 ($CH_3$), 32.81 (CH), 28.16 ($CH_3$), 16.94 ($CH_2$). HRMS (ESI) calculated for $C_{36}H_{39}N_2O_6$ $[M+H]^+$ = 595.2803, found 595.2804.

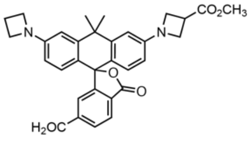

**6-carboxy-3″-methoxycarbonyl-JF$_{608}$ (S6).** Compound **S5** (32 mg, 54 µmol) was dissolved in $CH_2Cl_2$ (2 mL). To this solution was added TFA (0.4 mL), and the resulting blue solution was stirred at ambient temperature for 7 h. The reaction was concentrated under reduced pressure, yielding **S6** as a blue solid. HRMS (ESI): calculated for $C_{32}H_{31}N_2O_6$ $[M+H]^+$ = 539.2177, found 539.2182. This material was used immediately in the subsequent synthetic step.

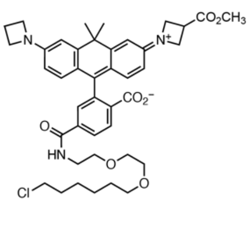


**3″-methoxycarbonyl-JF$_{608}$ -HaloTag ligand (S8).** Compound **S6** was dissolved in DMF (2 mL). To this solution were added DIEA (96 µL, 538 µmol, 10 equiv) and TSTU (24 mg, 81 µmol, 1.5 equiv). The reaction mixture was stirred for 10 min at ambient temperature, after which HaloTag($O_2$)-$NH_2$ (**S7**, 21 mg, 81 µmol, 1.5 equiv) was added. The reaction mixture was stirred for 16 h at ambient temperature. The solvent was removed under reduced pressure to yield **S8** as a blue solid. LC–MS (ESI) calculated for $C_{42}H_{51}ClN_3O_7$ $[M+H]^+$ = 744.34, found 744.40. This material was used immediately in the subsequent synthetic step.

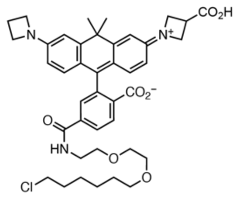


**3″-carboxy-JF$_{608}$ -HaloTag ligand (9).** NaOH (1 M, 108 µL) was added to a solution of **S8** in $CH_3OH$ (4 mL). The resulting solution was stirred for 14 h. The reaction mixture was acidified to pH ~1 using 1M HCl. This crude product was purified by preparative HPLC using a 25–75% $CH_3CN/H_2O$ linear gradient with a constant 0.1% v/v TFA. Product-containing fractions were combined and lyophilized to obtain **9** as a blue solid (TFA salt, 19 mg, 41.8% over three steps). $^1H$ NMR ($CDCl_3$, 400 MHz) δ 8.33 (d, *J* = 8.3 Hz, 1H), 8.15 (dd, *J* = 8.2, 1.8 Hz, 1H), 7.73 (d, *J* = 1.8 Hz, 1H), 6.94 (t, *J* = 9.0 Hz, 2H), 6.87 (d, *J* = 2.3 Hz, 1H), 6.84 (d, *J* = 2.2 Hz, 1H), 6.41 (t, *J* = 2.1 Hz, 1H), 6.39 (t, *J* = 2.1 Hz, 1H), 4.47 (t, *J* = 9.5 Hz, 2H), 4.41 – 4.30 (m, 6H), 3.72 – 3.67 (m, 1H), 3.66 – 3.59 (m, 5H), 3.58 – 3.55 (m, 3H), 3.52 (t, *J* = 6.6 Hz, 2H), 3.43 (t, *J* = 6.5 Hz, 2H), 2.55 (p, *J* = 7.6 Hz, 2H), 1.82 (s, 3H), 1.75 – 1.68 (m, 5H), 1.54 – 1.46 (m, 2H), 1.44 – 1.37 (m, 2H), 1.37 – 1.30 (m, 2H). $^{13}C$ NMR (101 MHz, $CDCl_3$) δ 175.37 (C), 168.10 (C), 167.62 (C), 158.20 (C), 157.25 (C), 156.98 (C), 156.31 (C), 140.23 (C), 139.03 (C), 137.85 (CH), 137.16 (CH), 134.71 (C), 132.24 (CH), 129.97 (CH), 129.35 (CH), 122.05 (C), 121.98 (C), 112.26 (CH), 111.82

(CH), 110.00 (CH), 109.79 (CH), 72.12 ($CH_2$), 71.20 ($CH_2$), 71.13 ($CH_2$), 70.36 ($CH_2$), 55.25 ($CH_2$), 53.06 ($CH_2$), 45.73 ($CH_2$), 42.70 (C), 41.15 ($CH_2$), 35.36 ($CH_3$), 33.91 (CH), 33.71 ($CH_2$), 32.27 ($CH_3$), 30.44 ($CH_2$), 27.68 ($CH_2$), 26.43 ($CH_2$), 16.83 ($CH_2$). HRMS (ESI) calculated for $C_{41}H_{49}ClN_3O_7$ $[M+H]^+$ = 730.3254, found 730.3255.

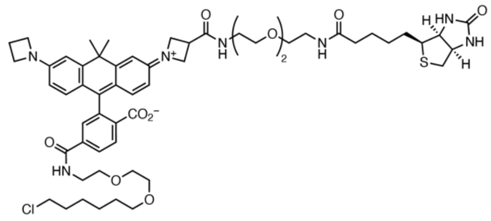

**Biotin-JF$_{608}$-HaloTag ligand (14$_{HTL}$).** The TFA salt of **9** (19 mg, 22.5 µmol, 1 equiv) was dissolved in DMF (3 mL). To this solution were added DIEA (40 µL, 225 µmol, 10 equiv), biotin-PEG$_2$-NH$_2$ (**12**, 9 mg, 33.7 µmol, 1.5 equiv), and HATU (12 mg, 33.8 µmol, 1.5 equiv). The reaction mixture was stirred for 18 h at ambient temperature. The solvent was removed under reduced pressure, and the product was purified by preparative HPLC using a 30–60% $CH_3CN/H_2O$ linear gradient with a constant 0.1% v/v TFA. Product-containing fractions were combined and lyophilized to obtain **14$_{HTL}$** as a blue solid (TFA salt, 19.8 mg, 73.3%). $^1$H NMR ($CD_3OD$, 400 MHz) δ 8.34 (d, $J$ = 8.3 Hz, 1H), 8.15 (dd, $J$ = 8.3, 1.8 Hz, 1H), 7.75 (d, $J$ = 1.8 Hz, 1H), 6.97 (d, $J$ = 6.8 Hz, 1H), 6.95 (d, $J$ = 6.8 Hz, 1H), 6.87 (d, $J$ = 2.2 Hz, 1H), 6.84 (d, $J$ = 2.2 Hz, 1H), 6.42 (t, $J$ = 2.4 Hz, 1H), 6.39 (t, $J$ = 2.5 Hz, 1H), 4.50 – 4.31 (m, 9H), 4.27 (dd, $J$ = 7.9, 4.4 Hz, 1H), 3.67 – 3.51 (m, 19H), 3.43 (t, $J$ = 6.4 Hz, 4H), 3.36 (t, $J$ = 5.5 Hz, 2H), 3.18 (dt, $J$ = 8.9, 5.3 Hz, 1H), 2.89 (ddd, $J$ = 12.8, 5.0, 1.0 Hz, 1H), 2.66 (dt, $J$ = 12.8, 1.2 Hz, 1H), 2.56 (p, $J$ = 7.6 Hz, 2H), 2.21 (t, $J$ = 7.3 Hz, 2H), 1.82 (s, 3H), 1.73– 1.63 (m, 6H), 1.67 – 1.56 (m, 3H), 1.51 – 1.47 (m, 2H), 1.46 – 1.31 (m, 6H). $^{13}$C NMR ($CD_3OD$, 101 MHz) δ 176.13 (C), 173.93 (C), 168.08 (C), 167.44 (C), 166.05 (C), 158.51 (C), 157.65 (C), 157.03 (C), 156.37 (C), 139.48 (C), 138.91 (C), 138.11 (CH), 137.51 (CH), 134.86 (C), 132.53 (CH), 130.26 (CH), 129.33 (CH), 122.03 (C), 121.99 (C), 112.21 (CH), 111.83 (CH), 109.98 (CH), 109.78 (CH), 72.12 ($CH_2$), 71.30 ($CH_2$), 71.20 ($CH_2$), 71.13 ($CH_2$), 70.60 ($CH_2$), 70.45 ($CH_2$), 70.36 ($CH_2$), 63.33 (CH), 61.60 (CH), 56.99 (CH), 55.24 ($CH_2$), 53.02 ($CH_2$), 45.75 ($CH_2$), 42.83 (C), 41.15 ($CH_2$), 41.06 ($CH_2$), 40.61 ($CH_2$), 40.27 ($CH_2$), 36.73 ($CH_2$), 35.40 ($CH_3$), 34.63 ($CH_3$), 33.71 ($CH_2$), 32.23 (CH), 30.45 ($CH_2$), 29.77 ($CH_2$), 29.50 ($CH_2$), 27.68 ($CH_2$), 26.84 ($CH_2$), 26.43 ($CH_2$), 16.80 ($CH_2$). $^{19}$F NMR ($CD_3OD$, 376 MHz) δ −75.45. HRMS (ESI) calculated for $C_{57}H_{77}N_7O_{10}SCl$ $[M+H]^+$ = 1086.5136, found 1086.5136.

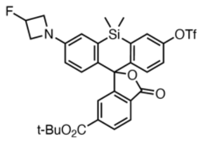

***tert*-butyl 3-(3-fluoroazetidin-1-yl)-5,5-dimethyl-3'-oxo-7-(((trifluoromethyl)sulfonyl)oxy)-3'*H*,5*H*-spiro[dibenzo[*b*,*e*]siline-10,1'-isobenzofuran]-6'-carboxylate (**S11**).** An oven-dried microwave reaction vial was charged with 6-*tert*-butoxycarbonyl-Si-fluorescein ditriflate[2] (**S9**, 370 mg, 500 µmol, 1 equiv), $Pd_2dba_3$ (46 mg, 50 µmol, 0.1 equiv), XantPhos (87 mg, 150 µmol, 0.3 equiv), 3-fluoroazetidine hydrochloride (**S10**, 62 mg, 550 µmol, 1.1 equiv), and $Cs_2CO_3$ (381 mg, 1.17 mmol, 2.4 equiv). The vial was sealed and backfilled with Ar(g) (3×), after which anhydrous dioxane (3 mL) was added. The resulting mixture was stirred at 80 °C for 2.5 h. The reaction was cooled and filtered through a pad of celite using $CH_2Cl_2$. The filtrate was concentrated under reduced pressure. Purification by $SiO_2$ gel chromatography (50 g $SiO_2$ column, 0–25% EtOAc/hexanes, linear gradient) provided monoazetidine compound **S11** as a light green foamy solid (210 mg). LC–MS (ESI): calculated for $C_{31}H_{29}F_4NO_7SSi$ $[M+H]^+$ = 663.14, found 664.15. This material was used immediately in the subsequent synthetic step.

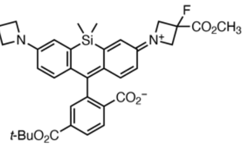


**4-(*tert*-butoxycarbonyl)-2-(3-(3-fluoro-3-(methoxycarbonyl)azetidin-1-ium-1-ylidene)-7-(3-fluoroazetidin-1-yl)-5,5-dimethyl-3,5-dihydrodibenzo[*b*,*e*]silin-10-yl)benzoate (*i.e.*, 3″-methoxycarbonyl-6-*tert*-butoxycarbonyl-$JF_{635}$; S13).** An oven-dried microwave reaction vial was charged with **S11** (210 mg, 316 µmol, 1 equiv), $Pd_2dba_3$ (29 mg, 32 µmol, 0.1 equiv), XPhos (55 mg, 95 µmol, 0.3 equiv), methyl 3-fluoroazetidine-3-carboxylate hydrochloride (**S12,** 161 mg, 949 µmol, 3 equiv), and $Cs_2CO_3$ (495 mg, 1.52 mmol, 4.8 equiv). The vial was sealed and backfilled with Ar(g) (3×), after which anhydrous dioxane (4 mL) was added. The resulting mixture was stirred at 100 °C for 4 h, then cooled to room temperature and filtered through a pad of celite using $CH_2Cl_2$. The filtrate was concentrated under reduced pressure. Purification by $SiO_2$ gel chromatography (50 g $SiO_2$ column, 0–40% EtOAc/hexanes, linear gradient) provided **S13** as a light green solid (177 mg, 55% over two steps). $^1$H NMR ($CDCl_3$, 400 MHz) δ 8.13 (dd, $J$ = 8.0, 1.3 Hz, 1H), 7.98 (dd, $J$ = 8.0, 0.7 Hz, 1H), 7.83 (t, $J$ = 1.0 Hz, 1H), 6.91 (dd, $J$ = 8.7, 7.2 Hz, 2H), 6.72 (dd, $J$ = 5.6, 2.7 Hz, 2H), 6.36 (ddd, $J$ = 8.7, 5.0, 2.7 Hz, 2H), 5.46 – 5.46 (m, 0.5H), 5.36 – 5.31 (m, 0.5H), 4.42 – 4.34 (m, 2H), 4.24 – 4.13 (m, 4H), 4.04 – 4.00 (m, 1H), 3.98 – 3.94 (m, 1H), 3.87 (s, 3H), 1.55 (s, 9H), 0.68 (s, 3H), 0.60 (s, 3H). $^{13}$C NMR (101 MHz, $CDCl_3$) δ 170.11 (C), 168.67 (d, $^2J_{CF}$ = 28.0 Hz, C), 164.32 (C), 155.11 (C), 149.90 (C), 149.05 (C), 137.37 (C), 136.40 (C), 136.16 (C), 133.99 (C), 133.37 (C), 130.07 (CH), 128.89 (C), 127.75 (CH), 127.73 (CH), 125.79 (CH), 125.00 (CH), 116.28 (CH), 116.23 (CH), 113.35 (CH), 113.22 (CH), 91.21 (C), 88.28 (d, $^1J_{CF}$ = 219.9 Hz, C), 82.76 (d, $^1J_{CF}$ = 204.9 Hz, CH), 82.43 (C), 61.22 (d, $^2J_{CF}$ = 25.1 Hz, $CH_2$), 59.55 (d, $^2J_{CF}$ = 23.8 Hz, $CH_2$), 53.25 ($CH_3$), 28.19 ($CH_3$), 0.11 ($CH_3$), −0.69 ($CH_3$). HRMS (ESI) calculated for $C_{35}H_{37}F_2N_2O_6Si$ $[M+H]^+$ = 647.2383, found 647.2376.

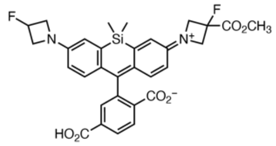

**6-carboxy-3″-methoxycarbonyl-JF$_{635}$ (S14).** Compound **S13** (16 mg, 25 µmol, 1 equiv) was dissolved in $CH_2Cl_2$ (2 mL). To this solution was added TFA (0.4 mL), and the resulting blue solution was stirred at ambient temperature for 4 h. The reaction was concentrated under reduced pressure, yielding **S14** as a blue solid. LC–MS (ESI): calculated for $C_{31}H_{29}F_2N_2O_6Si$ $[M]^+$ = 591.18, found 591.25. This material was used immediately in the subsequent synthetic step.

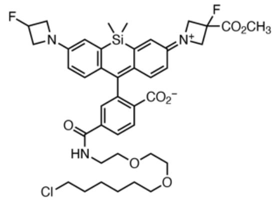

**3″-methoxycarbonyl-JF$_{635}$ -HaloTag ligand (S15).** Compound **S14** was dissolved in DMF (2 mL). To this solution were added *N*,*N*-Diisopropylethylamine (DIEA; 43 µL, 247 µmol, 10 equiv) and *N*,*N*,*N'*,*N'*-tetramethyl-*O*-(N-succinimidyl)uronium tetrafluoroborate (TSTU; 12.3 mg, 41 µmol, 1.65 equiv). The reaction mixture was stirred for 10 min at ambient temperature, after which HaloTag($O_2$)-$NH_2$ (**S7**, 8 mg, 31 µmol, 1.25 equiv) was added. The reaction mixture was stirred for 16 h at ambient temperature. The solvent was removed under reduced pressure to yield **S15** as a blue solid. LC–MS (ESI) calculated for $C_{41}H_{49}ClF_2N_3O_7Si$ $[M]^+$ = 796.30, found 796.20. This material was used immediately in the subsequent synthetic step.

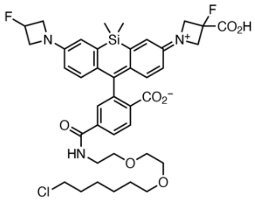

**3″-carboxy-JF$_{635}$ -HaloTag ligand (**11**).** Compound **S15** was dissolved in a mixture of THF:$CH_3OH$ (1:1 v/v; 4 mL). An aqueous solution of NaOH (1 M, 266 µL) was added, and the resulting solution and stirred for 7 h. The reaction mixture was diluted with $H_2O$ (2 mL) and acidified to pH ~1 using an aqueous solution of HCl (1 M). The mixture was extracted with EtOAc (4×2 mL), and the combined organics were concentrated under reduced pressure. This crude product was purified by preparative HPLC using a 5–95% $CH_3CN/H_2O$ linear gradient with a constant 0.1% v/v TFA. Product-containing fractions were combined and lyophilized to obtain **11** as a blue solid (TFA salt, 13.6 mg, 61% over three steps). $^1$H NMR ($CDCl_3$, 400 MHz) δ 8.02–8.0 (m, 1H), 7.92–7.90 (m, 1H), 7.72 (s, 1H), 7.20 (t, $J$ = 1.0 Hz, 1H), 6.80 (dd, $J$ = 8.7, 1.4 Hz, 2H), 6.72 (dd, $J$ = 12.6, 2.6 Hz, 2H), 6.62 (d, $J$ = 2.6 Hz, 1H), 6.31 (dd, $J$ = 8.8, 2.6 Hz, 1H), 6.26 (dd, $J$ = 8.9, 2.6 Hz, 1H), 5.52–5.49 (m, 0.5H), 5.35–5.33 (m, 0.5H), 4.39–4.24 (m, 4H), 4.18–4.01 (m, 4H), 3.64–3.62 (m, 6H), 3.58–3.55 (m, 2H), 3.49 (t, $J$ = 6.6 Hz, 2H), 3.42 (t, $J$ = 6.7 Hz, 2H), 1.75–1.67 (m, 2H), 1.44 (p, $J$ = 6.9 Hz, 2H), 1.43–1.35 (m, 2H), 1.33–1.27 (m, 2H), 0.53 (s, 3H), 0.46 (s, 3H). $^{13}$C NMR (101 MHz, $CDCl_3$) δ 169.99 (d, $^2J_{CF}$ = 28.3 Hz, C), 169.44 (C), 167.06 (C), 152.36 (C), 150.32 (C), 149.73 (C), 139.34 (C), 138.98 (C), 138.85 (C), 132.95 (C), 132.62 (C), 130.27 (C), 129.77 (C), 127.77 (CH), 127.20 (CH), 124.83 (CH), 117.29 (CH), 116.92 (CH), 113.42 (CH), 113.03 (CH), 87.73 (d, $^1J_{CF}$ = 220.4 Hz, C), 82.49 (d, $^1J_{CF}$ = 205.0 Hz, CH), 71.40 ($CH_2$), 70.22 ($CH_2$), 69.99 ($CH_2$), 69.48 ($CH_2$), 61.29 (d, $^2J_{CF}$ = 25.2 Hz, $CH_2$), 59.74 (d, $^2J_{CF}$ = 24.9 Hz, $CH_2$), 45.14 ($CH_2$), 40.35 ($CH_2$), 32.57 ($CH_2$), 29.35 ($CH_2$), 26.72 ($CH_2$), 25.41 ($CH_2$), 0.13 ($CH_3$), −1.43 ($CH_3$). One aromatic quaternary carbon was not observed. HRMS (ESI) calculated for $C_{40}H_{47}ClF_2N_3O_7Si$ $[M+H]^+$ = 782.2834, found 782.2823.

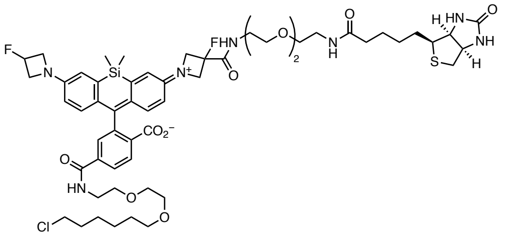

**Biotin-JF$_{635}$-HaloTag ligand (16$_{HTL}$).** The TFA salt of **11** (61.9 mg, 69 µmol, 1 equiv) was dissolved in DMF (2 mL). To this solution were added DIEA (121 µL, 690 µmol, 10 equiv), *N*-(3-dimethylaminopropyl)-*N'*-ethylcarbodiimide hydrochloride (EDC·HCl; 17.2 mg, 90 µmol, 1.3 equiv), and 1-[bis(dimethylamino)methylene]-1*H*-1,2,3-triazolo[4,5-*b*]pyridinium 3-oxid hexafluorophosphate (HATU; 34.2 mg, 90 µmol, 1.3 equiv). The reaction mixture was stirred at ambient temperature for 10 min, after which biotin-PEG$_2$-NH$_2$ (**12**, 27.4 mg, 104 µmol, 1.5 equiv) was added. The reaction mixture was stirred for a further 16 h at ambient temperature. The solvent was removed under reduced pressure, and the product was purified by preparative HPLC using a 40–55% $CH_3CN/H_2O$ linear gradient with a constant 0.1% v/v TFA. Product-containing fractions were combined and lyophilized to obtain **16$_{HTL}$** as a blue solid (TFA salt, 42.4 mg, 48.9%). $^1$H NMR ($CD_3OD$, 400 MHz) δ 7.97 (q, $J$ = 8.0 Hz, 2H), 7.59 (d, $J$ = 1.6 Hz, 1H), 6.78 (dd, $J$ = 14.7, 2.6 Hz, 2H), 6.72 (dd, $J$ = 8.9, 4.7 Hz, 2H), 6.30 (ddd, $J$ = 11.9, 8.9, 2.6 Hz, 2H), 5.40 (tt, $J$ = 6.1, 3.3 Hz, 0.5H), 5.26 (tt, $J$ = 6.1, 3.3 Hz, 1H), 4.38 – 4.25 (m, 3H), 4.25 – 4.10 (m, 5H), 4.05 – 3.88 (m, 2H), 3.54 – 3.34 (m, 19H), 3.29 (t, $J$ = 6.5 Hz, 2H), 3.25 (t, $J$ = 5.5 Hz, 2H), 3.02 (td, $J$ = 8.9, 4.1 Hz, 1H), 2.73 (dt, $J$ = 12.8, 4.7 Hz, 1H), 2.54 (dd, $J$ = 12.7, 1.9 Hz, 1H), 2.09 (t, $J$ = 7.4 Hz, 2H), 1.64 – 1.41 (m, 6H), 1.40 – 1.35 (m, 2H), 1.33 – 1.15 (m, 6H), 0.55 (s, 3H), 0.46 (s, 3H).

$^{13}C$ NMR ($CD_3OD$, 101 MHz, 320 K) δ 176.07 (C), 171.56 (C), 170.37 (d, $^{2}J_{CF}$ = 24.0 Hz, C), 168.66 (C), 165.98 (C), 152.01 (C), 151.24 (C), 141.72 (C), 138.96 (C), 134.26 (C), 133.81 (C), 129.76 (C), 129.36 (CH), 127.01 (CH), 124.76 (CH), 117.54 (CH), 114.38 (CH), 92.34 (d, $^{1}J_{CF}$ = 221.4 Hz, C), 84.29 (d, $^{1}J_{CF}$ = 202.7 Hz, CH), 72.15 ($CH_2$), 71.34 ($CH_2$), 71.32 ($CH_2$), 71.26 ($CH_2$), 71.14 ($CH_2$), 70.68 ($CH_2$), 70.36 ($CH_2$), 70.29 ($CH_2$), 63.36 (CH), 62.29 (d, $^{2}J_{CF}$ = 25.0 Hz, $CH_2$), 61.63 (CH), 60.59 (d, $^{2}J_{CF}$ = 24.1 Hz, $CH_2$), 56.90 (CH), 45.67 ($CH_2$), 41.17 ($CH_2$), 40.98 ($CH_2$), 40.35 ($CH_2$), 40.33 ($CH_2$), 36.77 ($CH_2$), 33.69 ($CH_2$), 30.41 ($CH_2$), 29.73 ($CH_2$), 29.48 ($CH_2$), 27.65 ($CH_2$), 26.77 ($CH_2$), 26.39 ($CH_2$), -0.14 ($CH_3$), -1.19 ($CH_3$). Two aromatic quaternary carbons were not observed. $^{19}F$ NMR ($CD_3OD$, 376 MHz) δ −75.55, −161.83, −180.00. Analytical HPLC: $t_R$= 11.6 min, 99.1% purity (5–95% MeCN/$H_2O$ linear gradient over 15 min with constant 0.1% v/v TFA, 1 mL/min flow rate, detection at 254 nm). HRMS (ESI) calculated for $C_{56}H_{75}N_7O_{10}F_2SClSiNa$ $[M+Na]^+$ = 1160.4536, found 1160.4545.

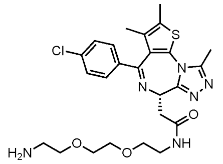

**(*S*)-JQ1-PEG$_2$-amine** (S19)**.** (+)-JQ1-$CO_2H$ (**S16**, 200 mg, 0.5 mmol, 1 equiv) was dissolved in DMF (4 mL). To this solution were added DIEA (448 µL, 1.25 mmol, 5 equiv), HATU (284 mg, 0.75 mmol, 1.5 equiv), and *tert*-butyl (2-(2-(2-aminoethoxy)ethoxy)ethyl)carbamate (**S17**, 186 mg, 0.75 mmol, 1.5 equiv). The reaction mixture was stirred at ambient temperature for 16 h. The solvent was removed under reduced pressure and resuspended in DCM (8 mL). TFA (2.5 mL) was added, and the reaction mixture was stirred at ambient temperature for 8 h. Volatiles were removed under reduced pressure and the product was purified by preparative HPLC using a 30–95% $CH_3CN/H_2O$ linear gradient with a constant 0.1% v/v TFA. Product-containing fractions were combined and lyophilized to obtain **S19** as a yellow sticky solid (5×TFA salt, 198.8 mg, 47.9% over two steps). The molar equivalents of TFA were determined via $^{19}F$ NMR by using fluorobenzene as an internal standard. $^{1}H$ NMR ($CD_3OD$, 400 MHz) δ 7.49 – 7.46 (m, 2H), 7.45 – 7.40 (m, 2H), 4.72 (dd, $J$ = 8.3, 6.0 Hz, 1H), 3.74 – 3.70 (m, 2H), 3.68 (s, 4H), 3.62 (t, $J$ = 5.6 Hz, 2H), 3.54 – 3.34 (m, 4H), 3.13 (t, $J$ = 5.0 Hz, 2H), 2.76 (s, 3H), 2.46 (s, 3H), 1.70 (s, 3H). $^{13}C$ NMR ($CD_3OD$, 101 MHz) δ 172.77 (C), 166.59 (C), 156.89 (C), 152.43 (C), 138.26 (C), 137.80 (C), 133.69 (C), 133.47 (C), 132.16 (C), 131.99 (C), 131.44 (CH), 129.86 (CH), 71.38 ($CH_2$), 71.36 ($CH_2$), 70.65 ($CH_2$), 67.91 ($CH_2$), 54.99 (CH), 40.69 ($CH_2$), 40.40 ($CH_2$), 38.51 ($CH_2$), 14.39 ($CH_3$), 12.94 ($CH_3$), 11.56 ($CH_3$); TFA peaks: 161.02 (q, $^{2}J_{CF}$ = 38.1 Hz, C), 117.34 (q, $^{1}J_{CF}$ = 288.4 Hz, C). HRMS (ESI) calculated for $C_{25}H_{32}ClN_6O_3S$ $[M+H]^+$ = 531.1940, found 531.1939.

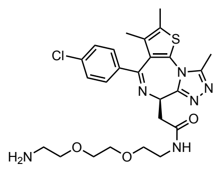

**(*R*)-JQ1-PEG$_2$-amine** (S23)**.** To a solution of (-)-JQ1-CO$_2$$^t$Bu (**S20**, 248.8 mg, 0.54 mmol, 1 equiv) in DCM (10 mL) was added TFA (1 mL). This light orange solution was stirred at ambient temperature for 24 h. The solvent was removed under reduced pressure and the crude was dissolved in DMF (3 mL). To this solution were successively added DIEA (976 µL, 5.44 mmol, 10 equiv), HATU (311 mg, 0.82 mmol, 1.5 equiv), and **S17** (203 mg, 0.82 mmol, 1.5 equiv). The reaction mixture was stirred at ambient temperature for 18 h. The solvent was removed under reduced pressure and the oil was resuspended in DCM (4 mL), and TFA (1 mL) was added. The reaction mixture was stirred at ambient temperature for 18 h. The solvent was removed under reduced pressure and the product was purified by preparative HPLC using a 5–95% $CH_3CN/H_2O$ linear gradient with a constant 0.1% v/v TFA. Product-containing fractions were combined and lyophilized to obtain **S23** as a yellow sticky solid (5×TFA salt, 350.7 mg, 58.5% over three steps). The molar equivalents of TFA were determined via $^{19}$F NMR by using fluorobenzene as an internal standard. $^1$H NMR ($CD_3OD$, 400 MHz) δ 7.49 – 7.44 (m, 2H), 7.43 – 7.42 (m, 2H), 4.71 (dd, $J$ = 8.3, 6.0 Hz, 1H), 3.74 – 3.70 (m, 2H), 3.68 (s, 4H), 3.62 (t, $J$ = 5.6 Hz, 2H), 3.53 – 3.34 (m, 4H), 3.13 (t, $J$ = 5.0 Hz, 2H), 2.75 (s, 3H), 2.46 (s, 3H), 1.70 (s, 3H). $^{13}$C NMR ($CD_3OD$, 101 MHz) δ 172.68 (C), 166.65 (C), 156.81 (C), 152.51 (C), 138.32 (C), 137.64 (C), 133.87 (C), 133.41 (C), 132.22 (C), 131.97 (C), 131.49 (CH), 129.86 (CH), 71.33 ($CH_2$), 70.62 ($CH_2$), 67.89 ($CH_2$), 54.90 (CH), 40.65 ($CH_2$), 40.39 ($CH_2$), 38.38 ($CH_2$), 14.40 ($CH_3$), 12.96 ($CH_3$), 11.55 ($CH_3$); TFA peaks: 161.41 (q, $^2J_{CF}$ = 38.3 Hz, C), 158.94 (q, $^2J_{CF}$ = 41.3 Hz, C), 117.34 (q, $^1J_{CF}$ = 289.0 Hz, C), 116.37 (q, $^1J_{CF}$ = 284.4 Hz, C). HRMS (ESI) calculated for $C_{25}H_{32}ClN_6O_3S$ $[M+H]^+$ = 531.1940, found 531.1945.

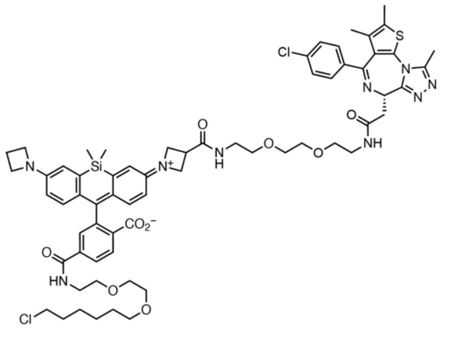

**(*S*)-JQ1-JF$_{646}$-HaloTag ligand** (19$_{HTL}$)**.** The TFA salt of **10** (23 mg, 27 µmol, 1 equiv) was dissolved in DMF (3 mL). To this solution were added DIEA (49 µL, 267 µmol, 10 equiv) and HATU (12.2 mg, 32 µmol, 1.2 equiv). The reaction mixture was stirred at ambient temperature for 5 min, after which **S19** (44.1 mg, 40 µmol, 1.5 equiv) was added. The reaction mixture was stirred for a further 16 h at ambient temperature. The solvent was removed under reduced pressure, and the product was purified by preparative HPLC using a 40–60% $CH_3CN/H_2O$ linear gradient with a constant 0.1% v/v TFA. Product-containing fractions were combined and lyophilized to obtain **$19_{HTL}$** as a blue solid (4xTFA salt, 29 mg, 63.2%). The molar equivalents of TFA were determined via $^{19}F$ NMR by using fluorobenzene as an internal standard. $^1H$ NMR ($CD_3OD$, 400 MHz; rotamers observed) δ 8.27 (dd, *J* = 8.2, 2.9 Hz, 1H), 8.11 (dt, *J* = 8.2, 2.0 Hz, 1H), 7.69 (dt, *J* = 7.3, 3.7 Hz, 1H), 7.41 (dd, *J* = 8.6, 2.9 Hz, 2H), 7.36 (d, *J* = 8.3 Hz, 2H), 6.93 – 6.76 (m, 4H), 6.34 (dd, *J* = 9.3, 2.6 Hz, 1H), 6.15 – 6.04 (m, 1H), 4.64 (dd, *J* = 8.8, 5.5 Hz, 1H), 4.36 – 4.18 (m, 8H), 3.66 – 3.55 (m, 16H), 3.53 – 3.48 (m, 3H), 3.46 – 3.40 (m, 7H), 3.36 – 3.33 (m, 1H), 2.61 (d, *J* = 8.9 Hz, 3H), 2.52 (p, *J* = 7.7 Hz, 2H), 2.43 (d, *J* = 2.9 Hz, 3H), 1.74 – 1.66 (m, 5H), 1.50 (p, *J* = 6.8 Hz, 2H), 1.38 (q, *J* = 7.6 Hz, 2H), 1.33 (dd, *J* = 9.0, 5.8 Hz, 2H), 0.56 (d, *J* = 8.3 Hz, 3H), 0.52 (d, *J* = 3.5 Hz, 3H). $^{13}C$ NMR ($CD_3OD$, 101 MHz; rotamers observed) δ 173.91 (C), 172.76 (C), 168.09 (C), 167.99 (C), 166.21 (C), 156.98 (C), 154.20 (C), 153.51 (C), 152.24 (C), 139.15 (C), 138.06 (C), 137.91 (C), 133.45 (C), 133.36 (C), 132.06 (C), 132.03 (C), 131.94 (C), 131.90 (C), 131.35 (CH), 129.81 (CH), 129.53 (CH), 128.97 (CH), 119.77 (CH), 119.15 (CH), 113.28 (CH), 112.74 (CH), 72.12 ($CH_2$), 71.40 ($CH_2$), 71.34 ($CH_2$), 71.21 ($CH_2$), 71.14 ($CH_2$), 70.61 ($CH_2$), 70.43 ($CH_2$), 70.38 ($CH_2$), 55.28 ($CH_2$), 55.09 (CH), 53.28 ($CH_2$), 45.74 ($CH_2$), 41.16 ($CH_2$), 40.66 ($CH_2$), 40.56 ($CH_2$), 38.71 ($CH_2$), 34.78 (CH), 33.71 ($CH_2$), 30.45 ($CH_2$), 27.68 ($CH_2$), 26.43 ($CH_2$), 16.97 ($CH_2$), 14.44 ($CH_3$), 12.98 ($CH_3$), 11.64 ($CH_3$), -0.81 ($CH_3$), -1.75 ($CH_3$). Five aromatic quaternary carbons were not observed. Analytical HPLC: $t_R$= 13.8 min, 96.5% purity (10–95% $CH_3CN/H_2O$ linear gradient over 20 min with constant 0.1% v/v TFA, 1 mL/min flow rate, detection at 650 nm). HRMS (ESI) calculated for $C_{65}H_{78}N_9O_9SCl_2Si$ $[M+H]^+$ = 1258.4790, found 1258.4790.

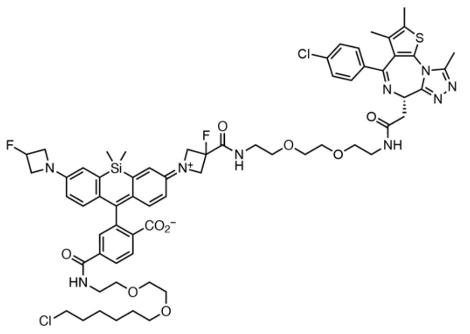

**(*S*)-JQ1-JF$_{635}$-HaloTag ligand (**$20_{HTL}$**).** The TFA salt of **11** (61.6 mg, 69 µmol, 1 equiv) was dissolved in DMF (3 mL). To this solution were added DIEA (121 µL, 109 µmol, 10 equiv), EDC·HCl (25.8 mg, 135 µmol, 2 equiv), HATU (51.3 mg, 135 µmol, 2 equiv). The reaction mixture was stirred at ambient temperature for 10 min, after which **S19** (55.1 mg, 50 µmol, 0.72 equiv) was added. The reaction mixture was stirred for a further 16 h at ambient temperature. The solvent was removed under reduced pressure, and the product was purified by preparative HPLC using a 40–65% $CH_3CN/H_2O$ linear gradient with a constant 0.1% v/v TFA. Product-containing fractions were combined and lyophilized to obtain **$20_{HTL}$** as a blue solid (4xTFA salt, 28.8 mg, 32.9%). The molar equivalents of TFA were determined via $^{19}F$ NMR by using fluorobenzene as an internal standard. $^{1}H$ NMR ($CD_3OD$, 400 MHz, 320 K) δ 8.07 (qd, *J* = 8.3, 3.6 Hz, 2H), 7.67 (d, *J* = 6.4 Hz, 1H), 7.43 (d, *J* = 8.3 Hz, 2H), 7.39 – 7.33 (m, 2H), 6.90 – 6.75 (m, 4H), 6.38 (dd, *J* = 8.9, 2.6 Hz, 1H), 6.30 (td, *J* = 8.7, 2.6 Hz, 1H), 5.53 – 5.46 (m, 0.5H), 5.39 – 5.32 (m, 0.5H), 4.70 – 4.58 (m, 1H; buried under the water peak), 4.48 – 4.27 (m, 4H), 4.26 – 4.14 (m, 2H), 4.14 – 4.00 (m, 2H), 3.65 – 3.56 (m, 12H), 3.55 – 3.50 (m, 4H), 3.50 – 3.33 (m, 10H), 2.66 (d, *J* = 8.3 Hz, 3H), 2.40 (d, *J* = 3.5 Hz, 3H), 1.73 – 1.63 (m, 5H), 1.47 (p, *J* = 6.8 Hz, 2H), 1.37 (p, *J* = 7.0 Hz, 2H), 1.29 (q, *J* = 8.3 Hz, 2H), 0.61 (d, *J* = 5.0 Hz, 3H), 0.53 (d, *J* = 4.9 Hz, 3H). $^{13}C$ NMR ($CD_3OD$, 101 MHz; rotamers observed) δ 172.69 (C), 170.72 (C), 170.20 (d, $^{2}J_{CF}$ = 21.3 Hz, C), 168.40 (C), 166.32 (C), 156.89 (C), 152.46 (C), 152.28 (C), 151.60 (C), 141.04 (C), 138.11 (C), 137.73 (C), 133.54 (C), 133.39 (C), 132.99 (C), 132.12 (C), 131.90 (C), 131.44 (CH), 129.80 (CH), 129.31 (CH), 128.04 (CH), 125.80 (CH), 118.87 (CH), 118.15 (CH), 114.27 (CH), 114.07 (CH), 92.03 (d, $^{1}J_{CF}$ = 221.9 Hz, C), 84.12 (d, $^{1}J_{CF}$ = 202.6 Hz, C), 72.11 ($CH_2$), 71.36 ($CH_2$), 71.26 ($CH_2$), 71.19 ($CH_2$), 71.11 ($CH_2$), 70.66 ($CH_2$), 70.33 ($CH_2$), 70.20 ($CH_2$), 62.30 (d, $^{2}J_{CF}$ = 25.1 Hz, $CH_2$), 60.62 (d, $^{2}J_{CF}$ = 24.5 Hz, $CH_2$), 54.99 (CH), 45.74 ($CH_2$), 41.14 ($CH_2$), 40.52 ($CH_2$), 40.28 ($CH_2$), 38.52 ($CH_2$), 33.68 ($CH_2$), 30.41 ($CH_2$), 27.66 ($CH_2$), 26.39 ($CH_2$), 14.46 ($CH_3$), 12.99 ($CH_3$), 11.64 ($CH_3$), -0.10 ($CH_3$), -1.30 ($CH_3$). TFA peaks: 161.33 (q, $^{2}J_{CF}$ = 37.5 Hz, C), 158.94 (q, $^{2}J_{CF}$ = 41.4 Hz, C), 116.04 (q, $^{1}J_{CF}$ = 285.1 Hz, C). Five aromatic quaternary carbons were not observed. $^{19}F$ NMR ($CD_3OD$, 376 MHz) δ −75.14, -75.62, −161.97, −179.86. Analytical HPLC: $t_R$= 14.7 min, 98.4% purity (30–95% $CH_3CN/H_2O$ linear gradient over 20 min with constant 0.1% v/v TFA, 1 mL/min flow rate, detection at 650 nm). HRMS (ESI) calculated for $C_{65}H_{76}N_9O_9F_2SCl_2Si$ $[M+H]^+$ = 1294.4596, found 1294.4609.

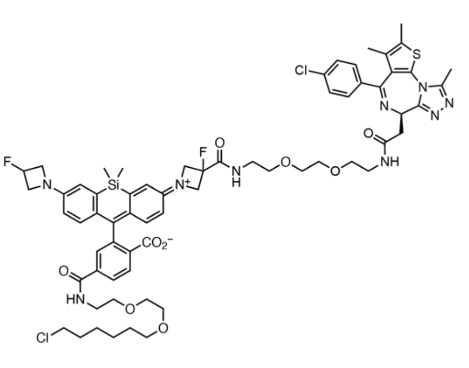

**(*R*)-JQ1-$JF_{635}$-HaloTag ligand ($21_{HTL}$).** The TFA salt of **11** (8.5 mg, 11 μmol, 1 equiv) was dissolved in DMF (2 mL). To this solution were added DIEA (20 μL, 109 μmol, 10 equiv), **S23** (30 mg, 26 μmol, 1.7 equiv), EDC·HCl (6.1 mg, 32 μmol, 3 equiv), and HATU (12.2 mg, 32 μmol, 3 equiv). The reaction mixture was stirred at ambient temperature for 16 h. The solvent was removed under reduced pressure, and the product was purified by preparative HPLC using a 5–95% $CH_3CN/H_2O$ linear gradient with a constant 0.1% v/v TFA. Product-containing fractions were combined and lyophilized to obtain **$21_{HTL}$** as a blue solid (TFA salt, 8.8 mg, 52.5%). $^{1}H$ NMR

($CD_3OD$, 400 MHz) δ 8.36 (br s, 1 H), 8.07 – 8.03 (m, 2H), 7.69 – 7.66 (m, 1H), 7.42 – 7.35 (m, 4H), 6.84 – 6.79 (m, 3H), 6.74 (d, *J* = 8.8 Hz, 1H), 6.40 (ddd, *J* = 8.9, 2.7, 1.6 Hz, 1H), 6.29 – 6.23 (m, 1H), 5.52 – 5.48 (m, 0.5H), 5.39 – 5.34 (m, 0.5H), 4.66 – 4.61 (m, 1H), 4.45 – 3.93 (m, 8H), 3.67 – 3.41 (m, 22H), 3.39 (t, *J* = 6.5 Hz, 2H), 3.29 – 3.27 (m, 2H), 2.65 (2 x s, 3H), 2.41 (2 x s, 3H), 1.73 – 1.60 (m, 5H), 1.46 (p, *J* = 6.8 Hz, 2H), 1.40– 1.33 (m, 2H), 1.31 – 1.27 (m, 2H), 0.62 (2 x s, 3H), 0.53 (2 x s, 3H). Analytical HPLC: $t_R$= 15.7 min, 97.7% purity (5–95% $CH_3CN/H_2O$ linear gradient over 20 min with constant 0.1% v/v TFA, 1 mL/min flow rate, detection at 254 nm). HRMS (ESI) calculated for $C_{65}H_{76}N_9O_9F_2SCl_2Si$ $[M+H]^+$ = 1294.4596, found 1294.4597.

### NMR Spectra and HPLC Traces

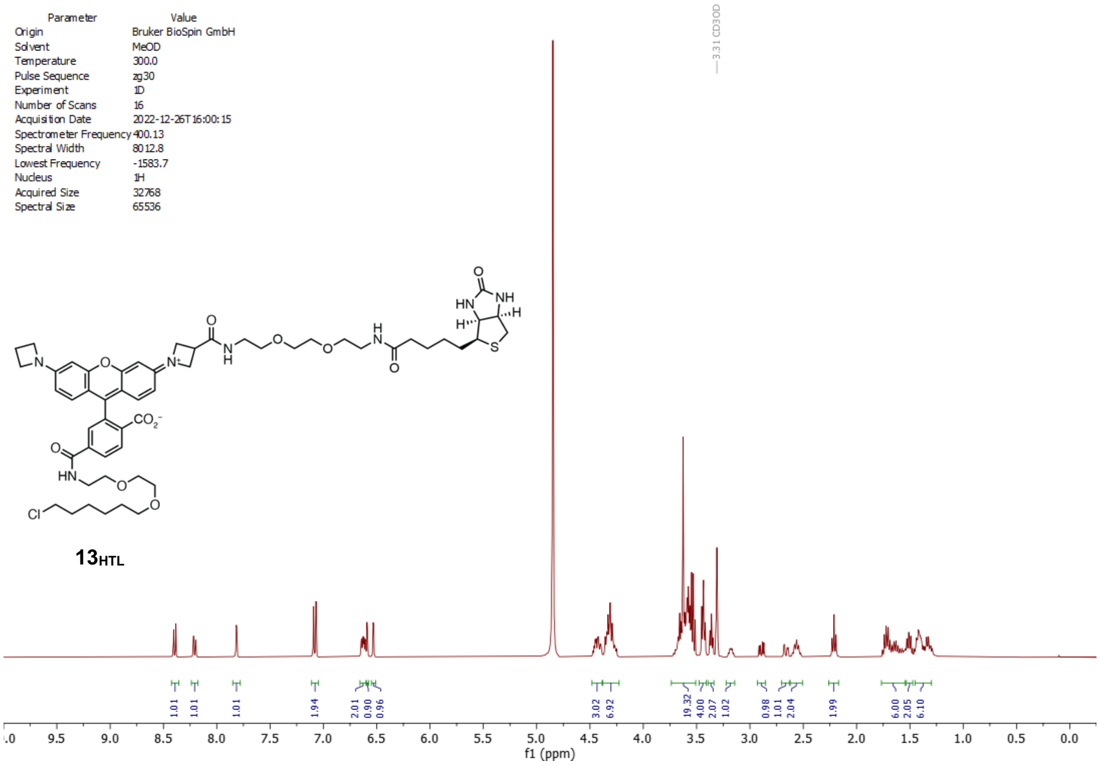


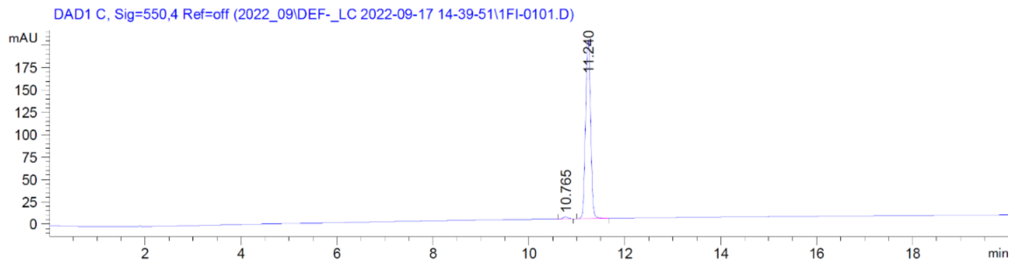


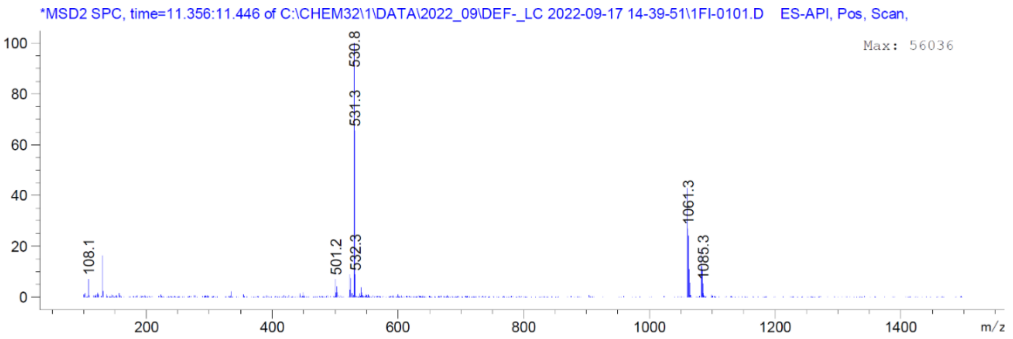

| Parameter | Value |
| --- | --- |
| Origin | Bruker BioSpin GmbH |
| Solvent | MeOD |
| Temperature | 300.0 |
| Pulse Sequence | zgpg30 |
| Experiment | 1D |
| Number of Scans | 4512 |
| Acquisition Date | 2022-12-26T15:53:17 |
| Spectrometer Frequency | 100.62 |
| Spectral Width | 24038.5 |
| Lowest Frequency | -1818.1 |
| Nucleus | 13C |
| Acquired Size | 32768 |
| Spectral Size | 65536 |

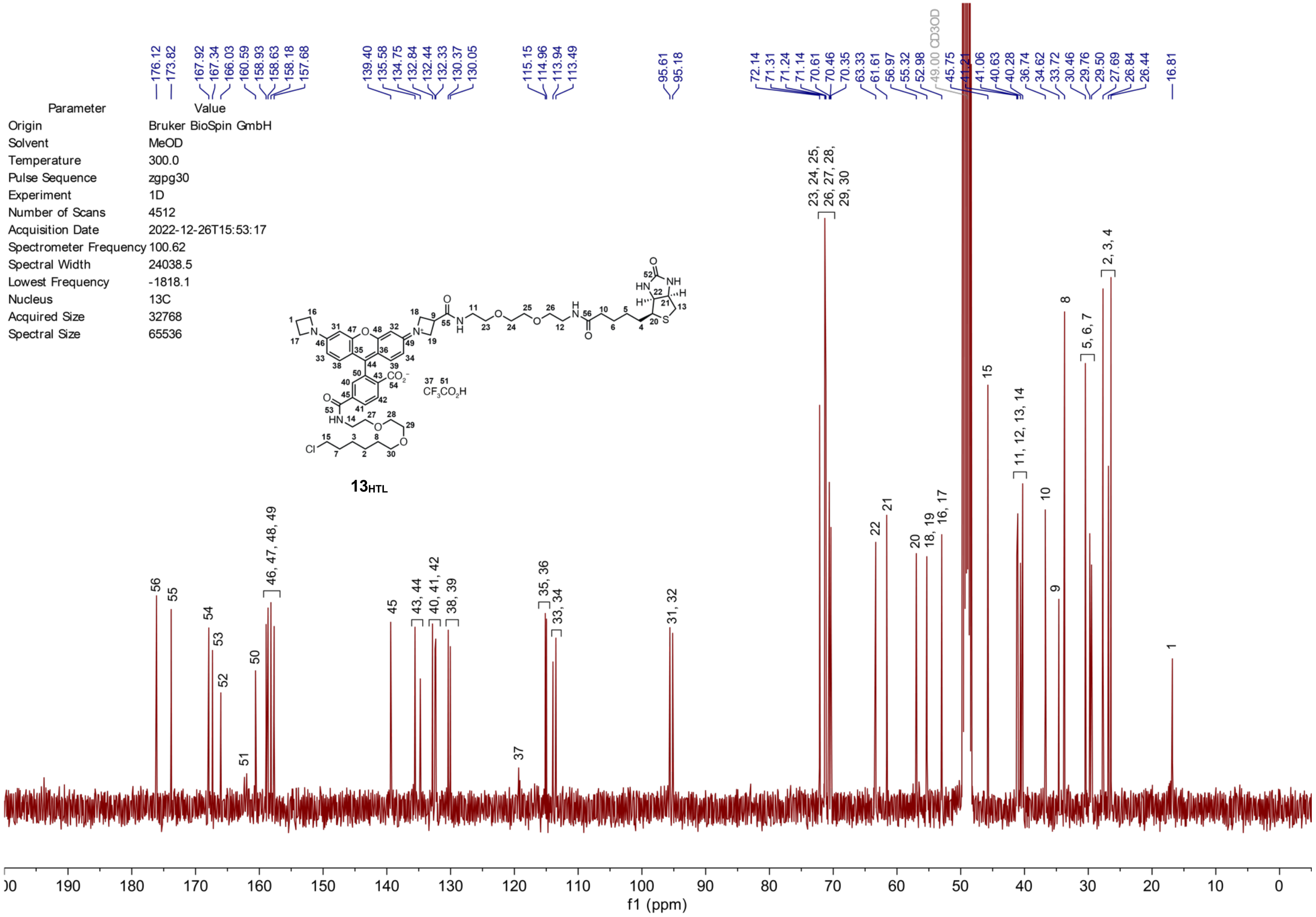

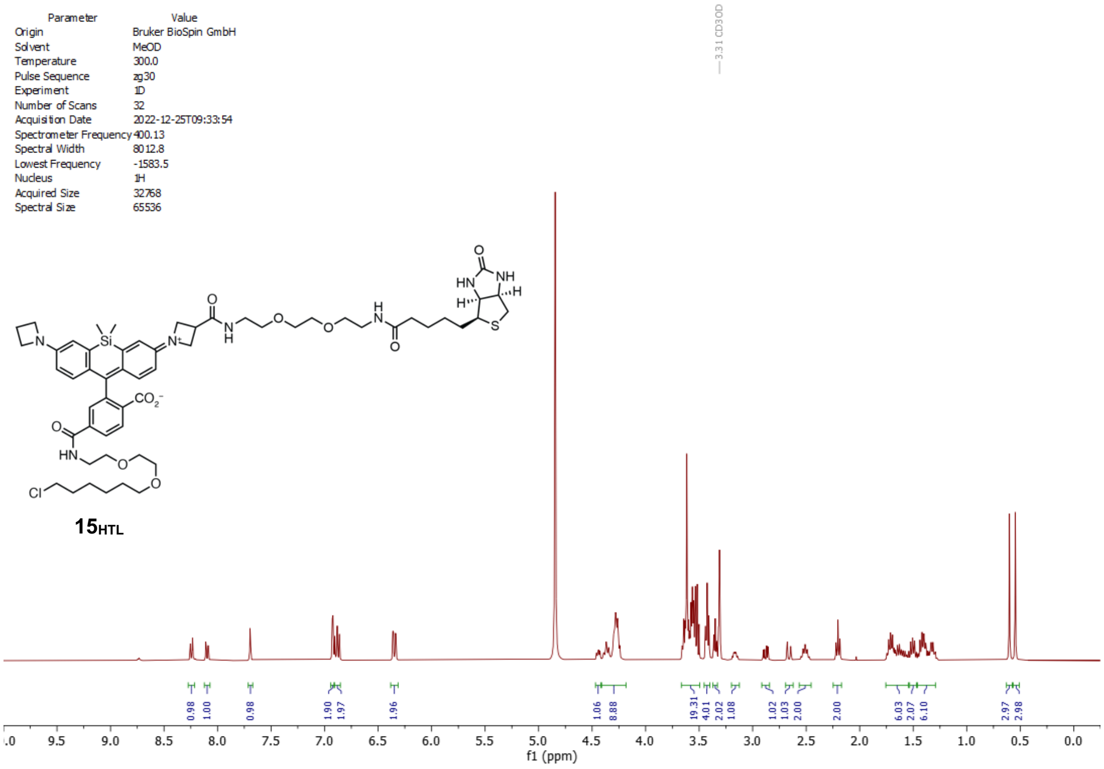
Parameter Value
Origin Bruker BioSpin GmbH
Solvent MeOD
Temperature 300.0
Pulse Sequence zg30
Experiment 1D
Number of Scans 32
Acquisition Date 2022-12-25T09:33:54
Spectrometer Frequency 400.13
Spectral Width 8012.8
Lowest Frequency -1583.5
Nucleus 1H
Acquired Size 32768
Spectral Size 65536
3.31 CD3OD
15HTL
0.98 1.00 0.98 1.90 1.97 1.96 1.06 8.88 19.31 4.01 2.02 1.08 1.02 1.03 2.00 2.00 6.03 2.07 6.10 2.97 2.98
f1 (ppm)

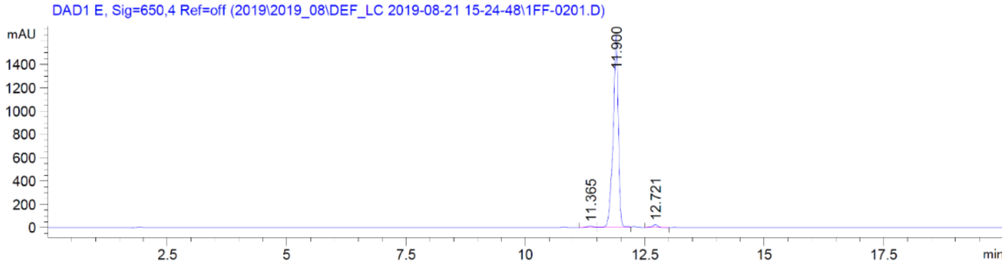
DAD1 E, Sig=650,4 Ref=off (2019\2019_08\DEF_LC 2019-08-21 15-24-48\1FF-0201.D)
mAU
11.900
11.365
12.721
min

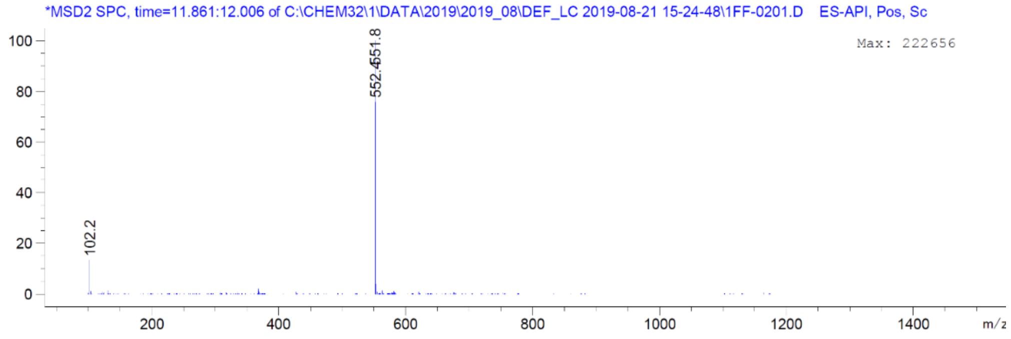
*MSD2 SPC, time=11.861:12.006 of C:\CHEM32\1\DATA\2019\2019_08\DEF_LC 2019-08-21 15-24-48\1FF-0201.D ES-API, Pos, Sc
Max: 222656
552.4551.8
102.2
m/z

| Parameter | Value |
|---|---|
| Origin | Bruker BioSpin GmbH |
| Solvent | MeOD |
| Temperature | 300.0 |
| Pulse Sequence | zgpg30 |
| Experiment | 1D |
| Number of Scans | 19504 |
| Acquisition Date | 2022-12-26T10:50:14 |
| Spectrometer Frequency | 100.62 |
| Spectral Width | 24038.5 |
| Lowest Frequency | -1818.4 |
| Nucleus | 13C |
| Acquired Size | 32768 |
| Spectral Size | 65536 |

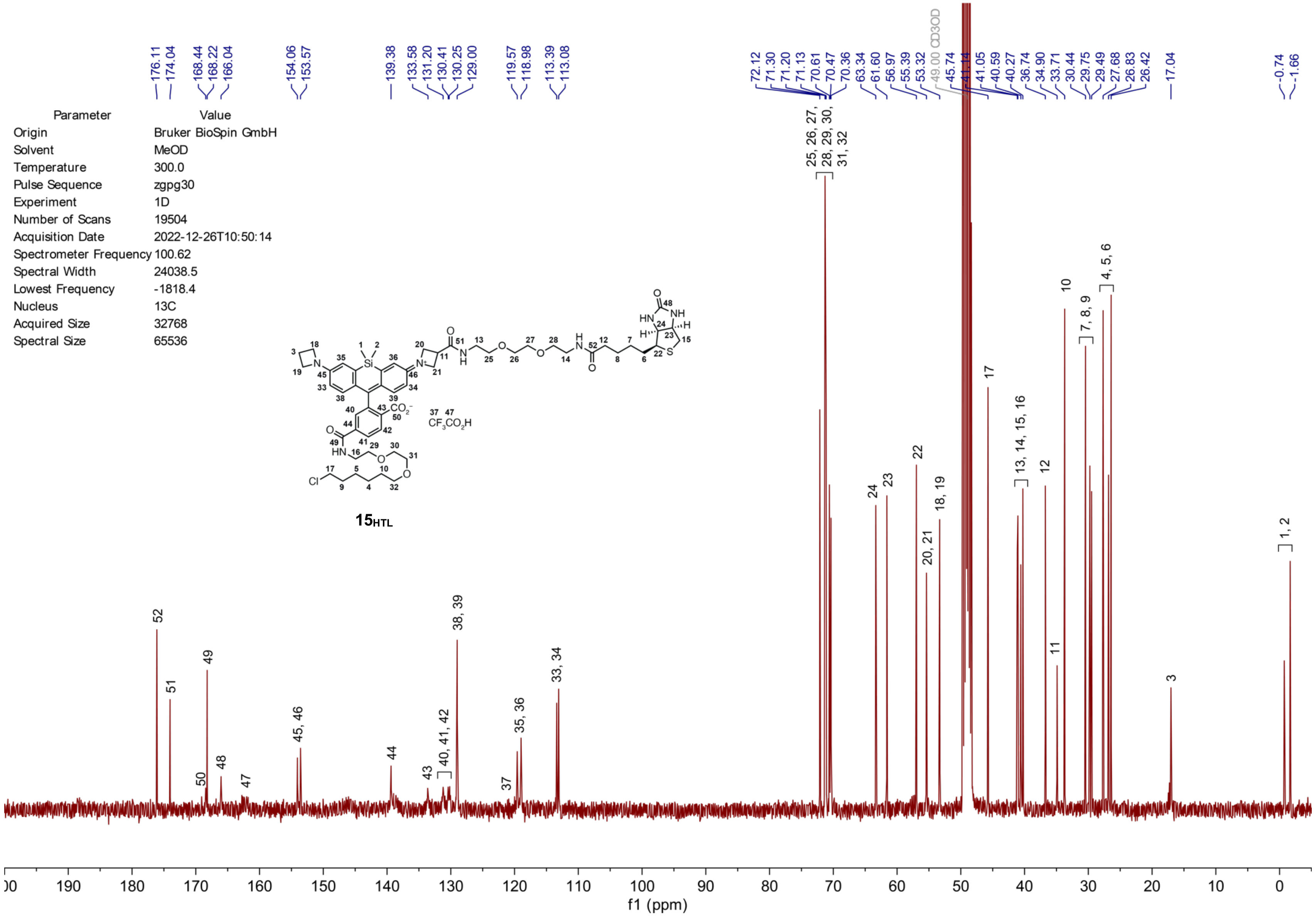

| Parameter | Value |
|---|---|
| Solvent | CDCl3 |
| Temperature | 295.8 |
| Pulse Sequence | zg30 |
| Experiment | 1D |
| Number of Scans | 16 |
| Acquisition Date | 2023-06-29T09:52:15 |
| Spectrometer Frequency | 400.13 |
| Spectral Width | 8012.8 |
| Lowest Frequency | -1545.6 |
| Nucleus | 1H |
| Acquired Size | 32768 |
| Spectral Size | 65536 |

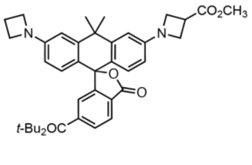


**S3**

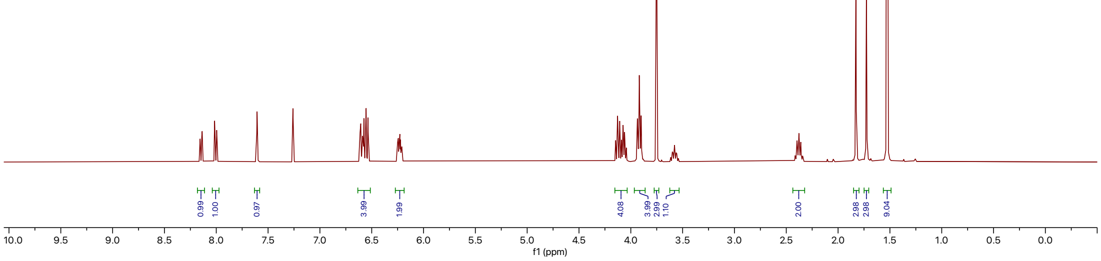

| Parameter | Value |
|---|---|
| Solvent | CDCl3 |
| Temperature | 296.2 |
| Pulse Sequence | zgpg30 |
| Experiment | 1D |
| Number of Scans | 1024 |
| Acquisition Date | 2023-06-29T15:33:45 |
| Spectrometer Frequency | 100.62 |
| Spectral Width | 24038.5 |
| Lowest Frequency | -1947.0 |
| Nucleus | 13C |
| Acquired Size | 32768 |
| Spectral Size | 65536 |

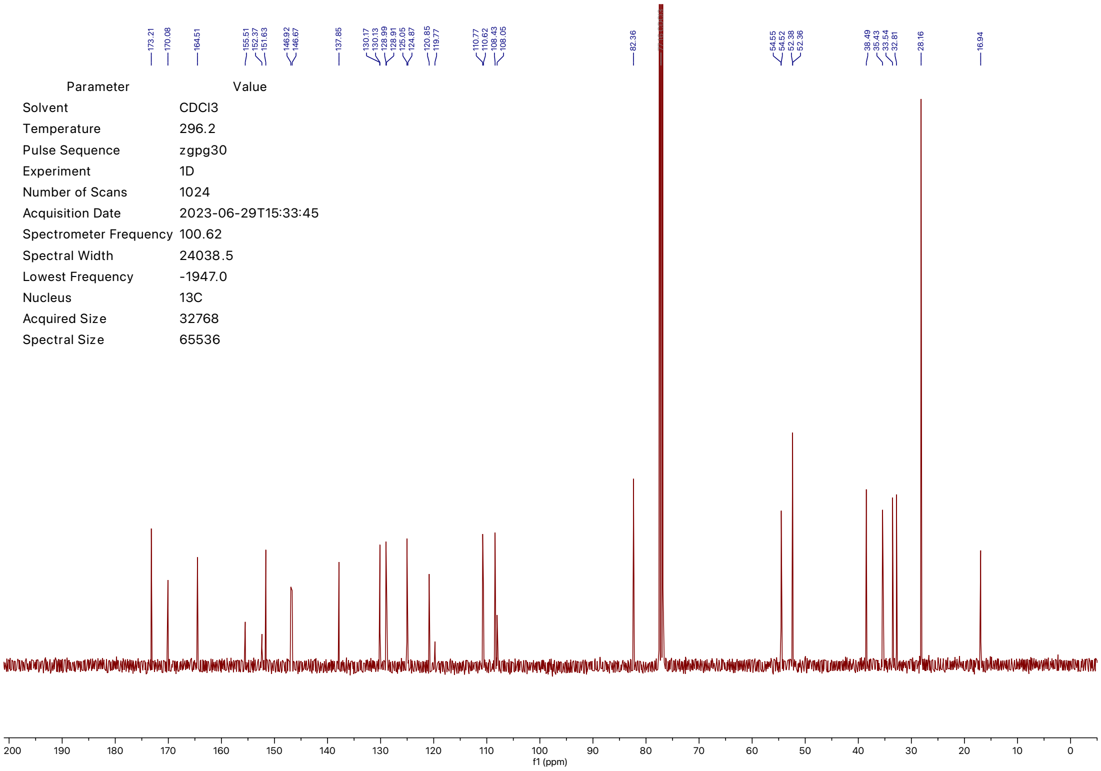

| Parameter | Value |
| --- | --- |
| Solvent | MeOD |
| Temperature | 295.7 |
| Pulse Sequence | zg30 |
| Experiment | 1D |
| Number of Scans | 16 |
| Acquisition Date | 2023-07-17T08:33:56 |
| Spectrometer Frequency | 400.13 |
| Spectral Width | 8012.8 |
| Lowest Frequency | -1543.3 |
| Nucleus | 1H |
| Acquired Size | 32768 |
| Spectral Size | 65536 |

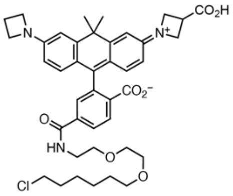


**9**

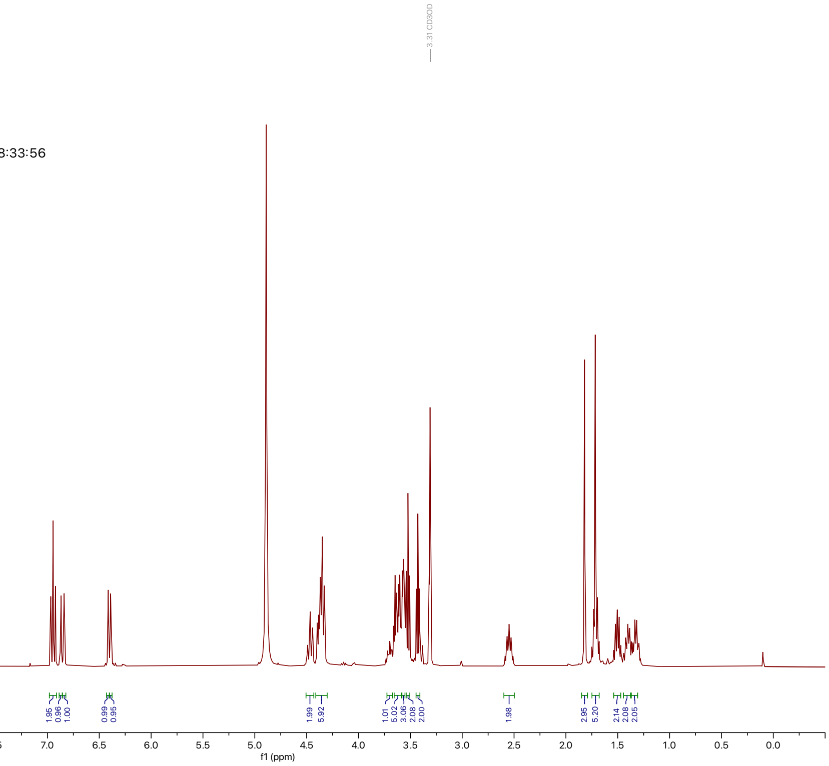


| Parameter | Value |
| --- | --- |
| Solvent | MeOD |
| Temperature | 296.0 |
| Pulse Sequence | zgpg30 |
| Experiment | 1D |
| Number of Scans | 9000 |
| Acquisition Date | 2023-07-17T04:35:34 |
| Spectrometer Frequency | 100.62 |
| Spectral Width | 24038.5 |
| Lowest Frequency | -1818.0 |
| Nucleus | 13C |
| Acquired Size | 32768 |
| Spectral Size | 65536 |

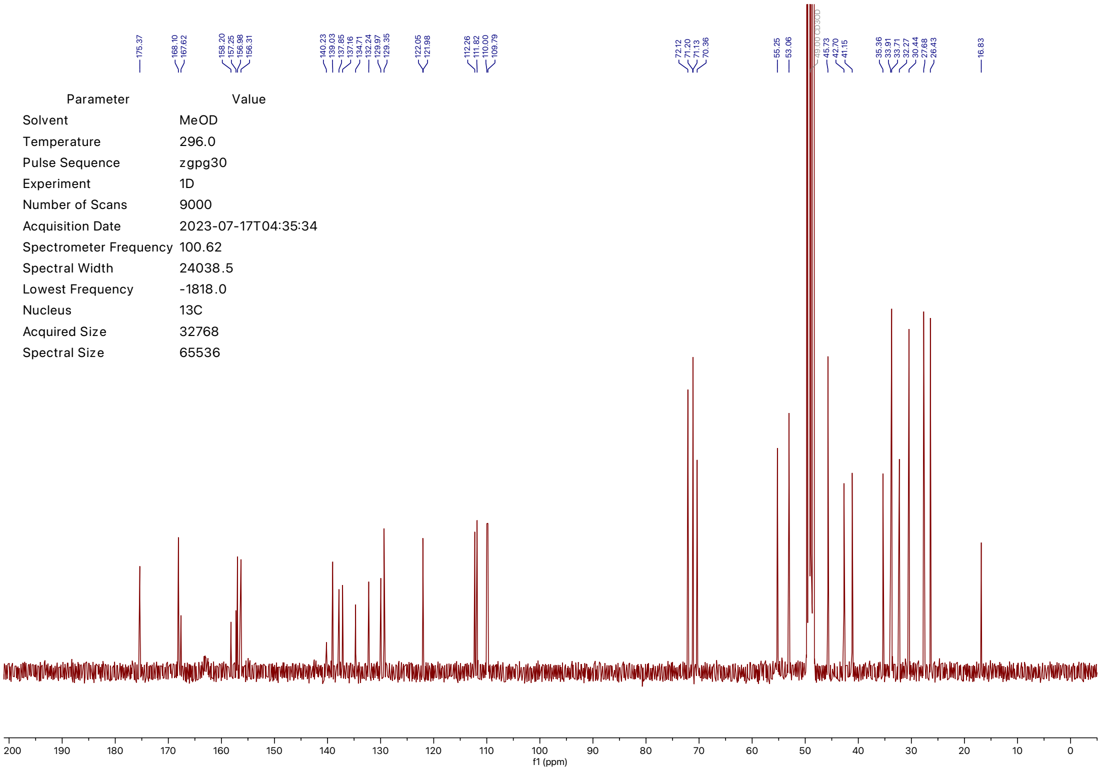

| Parameter | Value |
|---|---|
| Solvent | MeOD |
| Temperature | 295.8 |
| Pulse Sequence | zg30 |
| Experiment | 1D |
| Number of Scans | 16 |
| Acquisition Date | 2023-07-26T10:33:16 |
| Spectrometer Frequency | 400.13 |
| Spectral Width | 8012.8 |
| Lowest Frequency | -1543.3 |
| Nucleus | 1H |
| Acquired Size | 32768 |
| Spectral Size | 65536 |

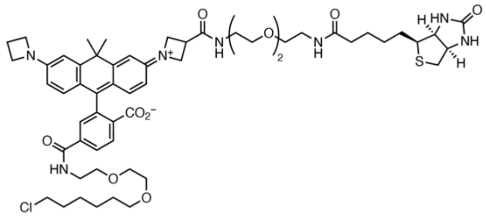


**14**$_{HTL}$

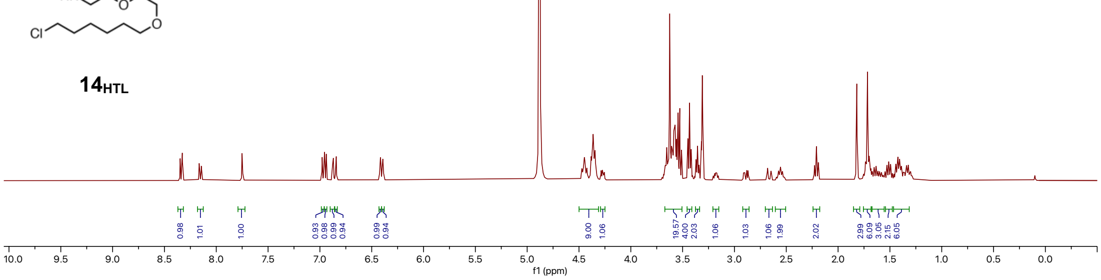

| Parameter | Value |
| --- | --- |
| Solvent | MeOD |
| Temperature | 296.3 |
| Pulse Sequence | zgpg30 |
| Experiment | 1D |
| Number of Scans | 15000 |
| Acquisition Date | 2023-07-26T10:30:33 |
| Spectrometer Frequency | 100.62 |
| Spectral Width | 24038.5 |
| Lowest Frequency | -1818.7 |
| Nucleus | 13C |
| Acquired Size | 32768 |
| Spectral Size | 65536 |

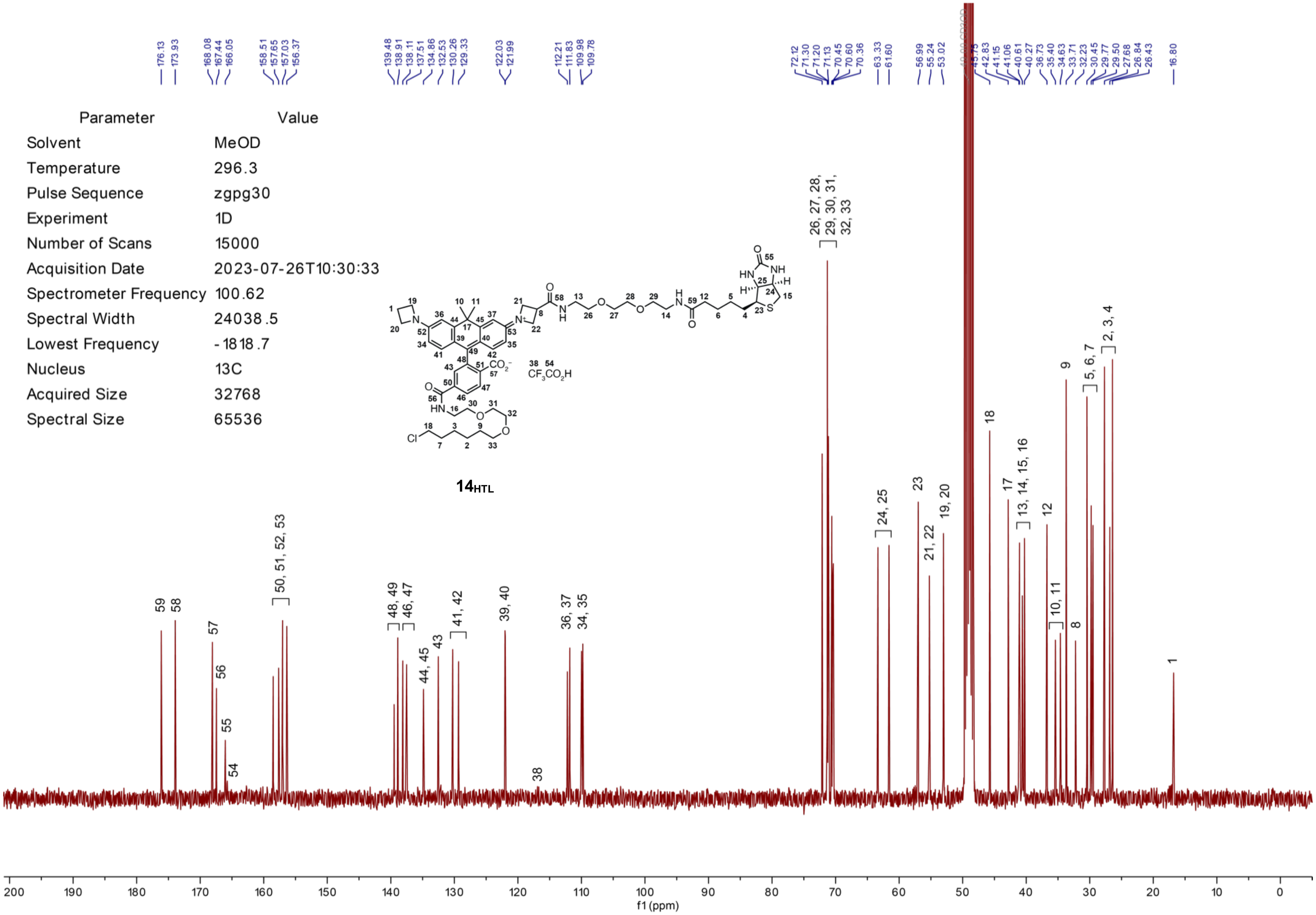

| Parameter | Value |
|---|---|
| Origin | Bruker BioSpin GmbH |
| Solvent | CDCl3 |
| Temperature | 300.0 |
| Pulse Sequence | zg30 |
| Experiment | 1D |
| Number of Scans | 23 |
| Acquisition Date | 2019-09-06T08:55:00 |
| Spectrometer Frequency | 400.13 |
| Spectral Width | 8012.8 |
| Lowest Frequency | -1545.4 |
| Nucleus | 1H |
| Acquired Size | 32768 |
| Spectral Size | 65536 |

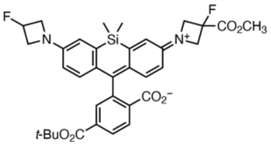


**S13**

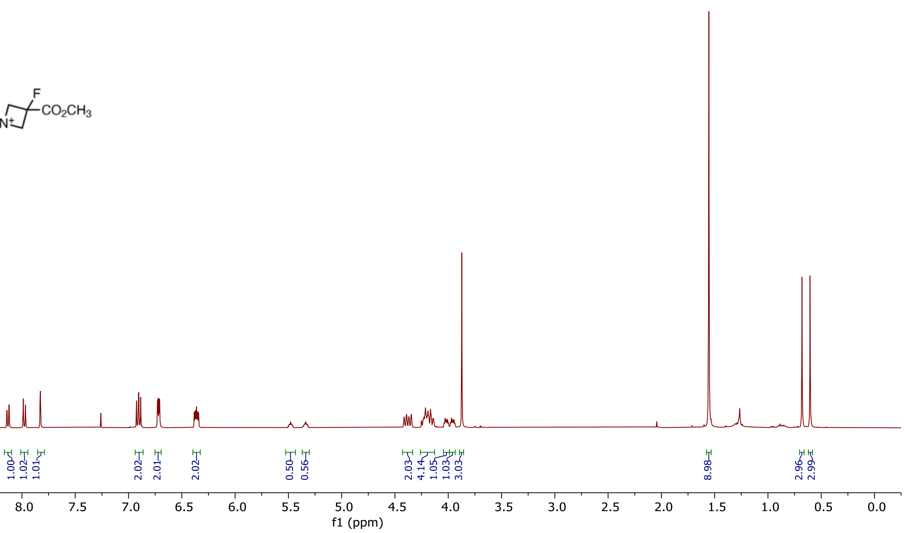


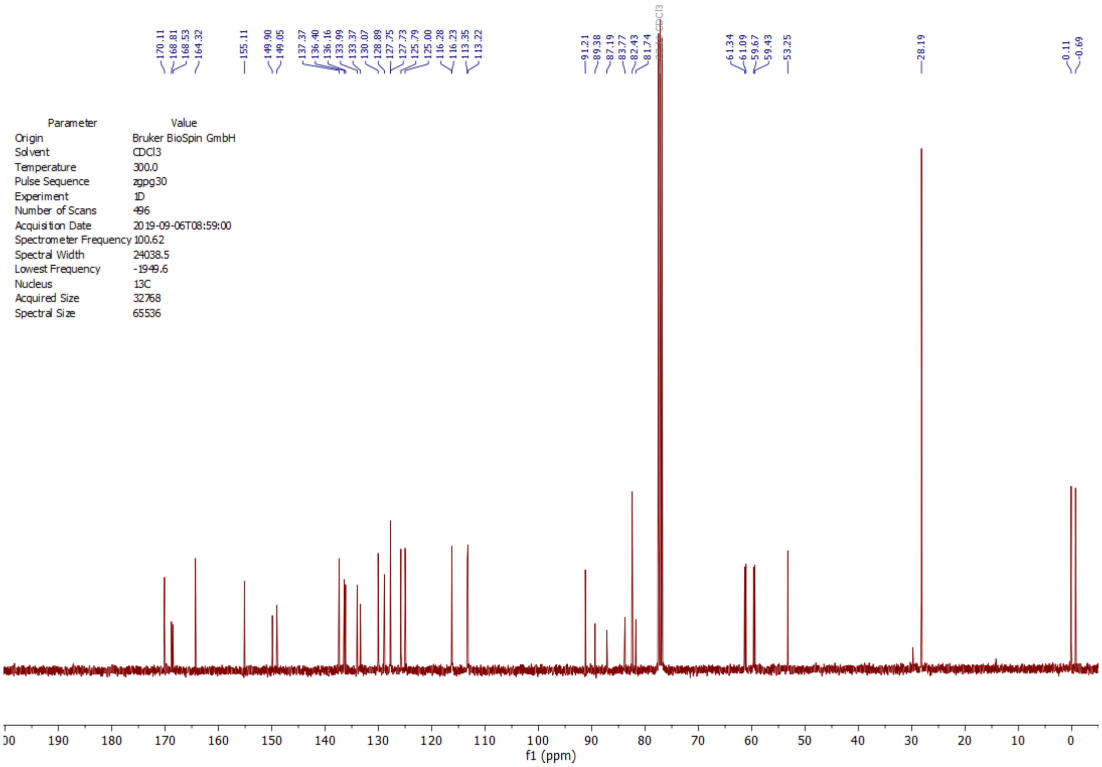


| Parameter | Value |
|---|---|
| Origin | Bruker BioSpin GmbH |
| Solvent | CDCl3 |
| Temperature | 300.0 |
| Pulse Sequence | zgpg30 |
| Experiment | 1D |
| Number of Scans | 496 |
| Acquisition Date | 2019-09-06T08:59:00 |
| Spectrometer Frequency | 100.62 |
| Spectral Width | 24038.5 |
| Lowest Frequency | -1949.6 |
| Nucleus | 13C |
| Acquired Size | 32768 |
| Spectral Size | 65536 |

| Parameter | Value |
|---|---|
| Origin | Bruker BioSpin GmbH |
| Solvent | CDCl3 |
| Temperature | 300.0 |
| Pulse Sequence | zg30 |
| Experiment | 1D |
| Number of Scans | 71 |
| Acquisition Date | 2019-09-08T19:01:00 |
| Spectrometer Frequency | 400.13 |
| Spectral Width | 8012.8 |
| Lowest Frequency | -1545.4 |
| Nucleus | 1H |
| Acquired Size | 32768 |
| Spectral Size | 65536 |

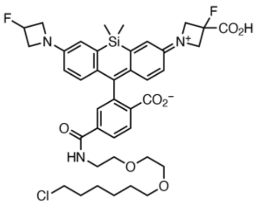


**11**

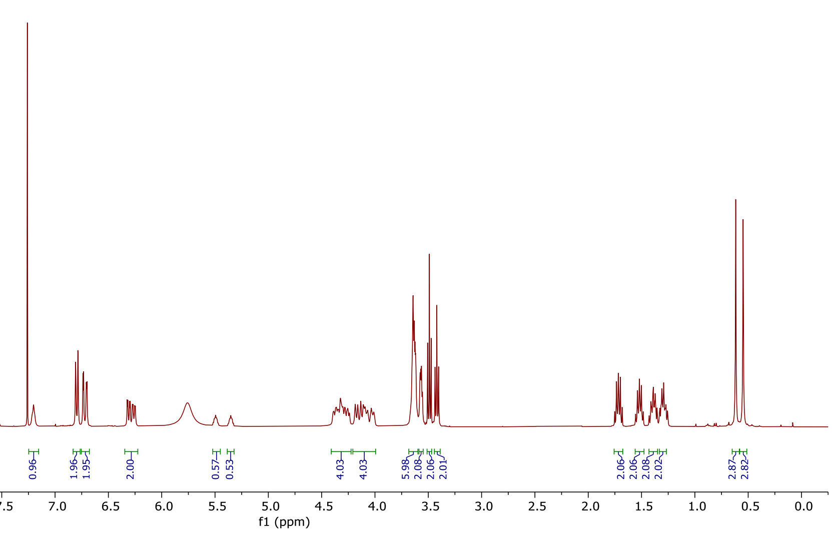


| Parameter | Value |
|---|---|
| Comment | |
| Origin | Bruker BioSpin GmbH |
| Solvent | CDCl3 |
| Temperature | 300.0 |
| Pulse Sequence | zgpg30 |
| Experiment | 1D |
| Number of Scans | 13000 |
| Acquisition Date | 2019-09-09T07:43:00 |
| Spectrometer Frequency | 100.62 |
| Spectral Width | 24038.5 |
| Lowest Frequency | -1944.1 |
| Nucleus | 13C |
| Acquired Size | 32768 |
| Spectral Size | 65536 |

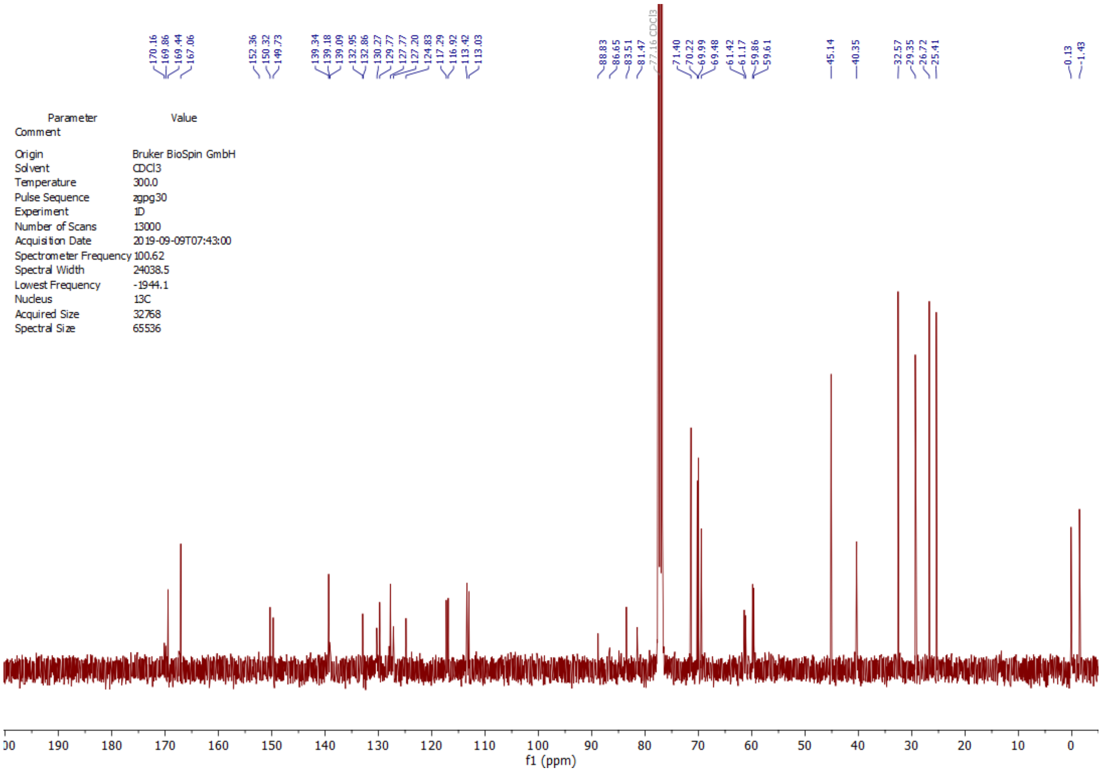

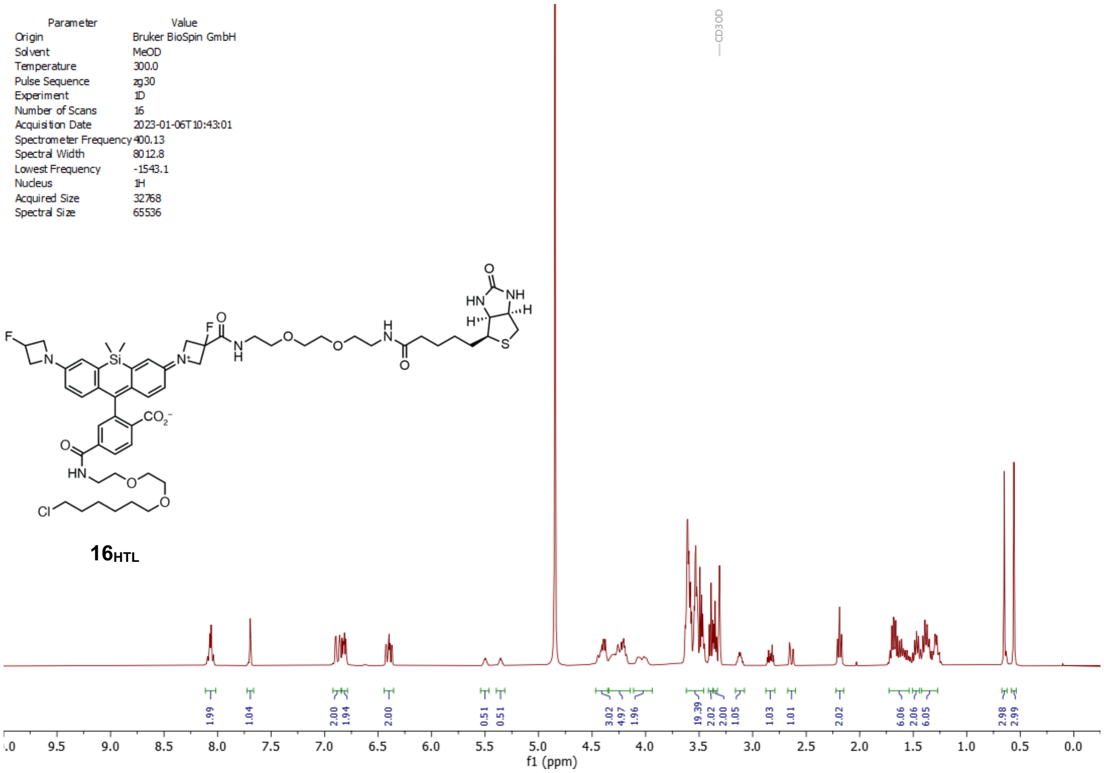

Parameter Value
Origin Bruker BioSpin GmbH
Solvent MeOD
Temperature 300.0
Pulse Sequence zg30
Experiment 1D
Number of Scans 16
Acquisition Date 2023-01-06T10:43:01
Spectrometer Frequency 400.13
Spectral Width 8012.8
Lowest Frequency -1543.1
Nucleus 1H
Acquired Size 32768
Spectral Size 65536
CD3OD
f1 (ppm)


**16HTL**

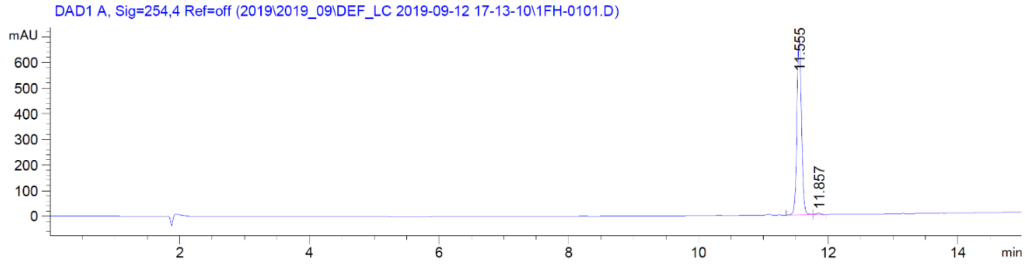

DAD1 A, Sig=254,4 Ref=off (2019\2019_09\DEF_LC 2019-09-12 17-13-10\1FH-0101.D)
mAU
11.555
11.857
min


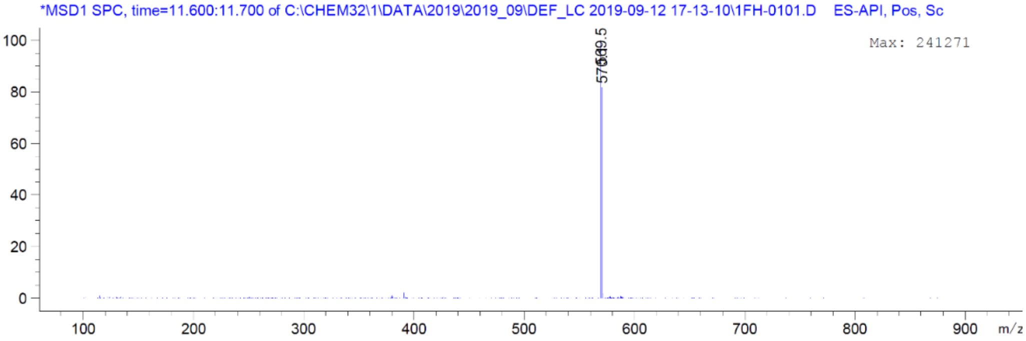

*MSD1 SPC, time=11.600:11.700 of C:\CHEM32\1\DATA\2019\2019_09\DEF_LC 2019-09-12 17-13-10\1FH-0101.D ES-API, Pos, Sc
Max: 241271
m/z

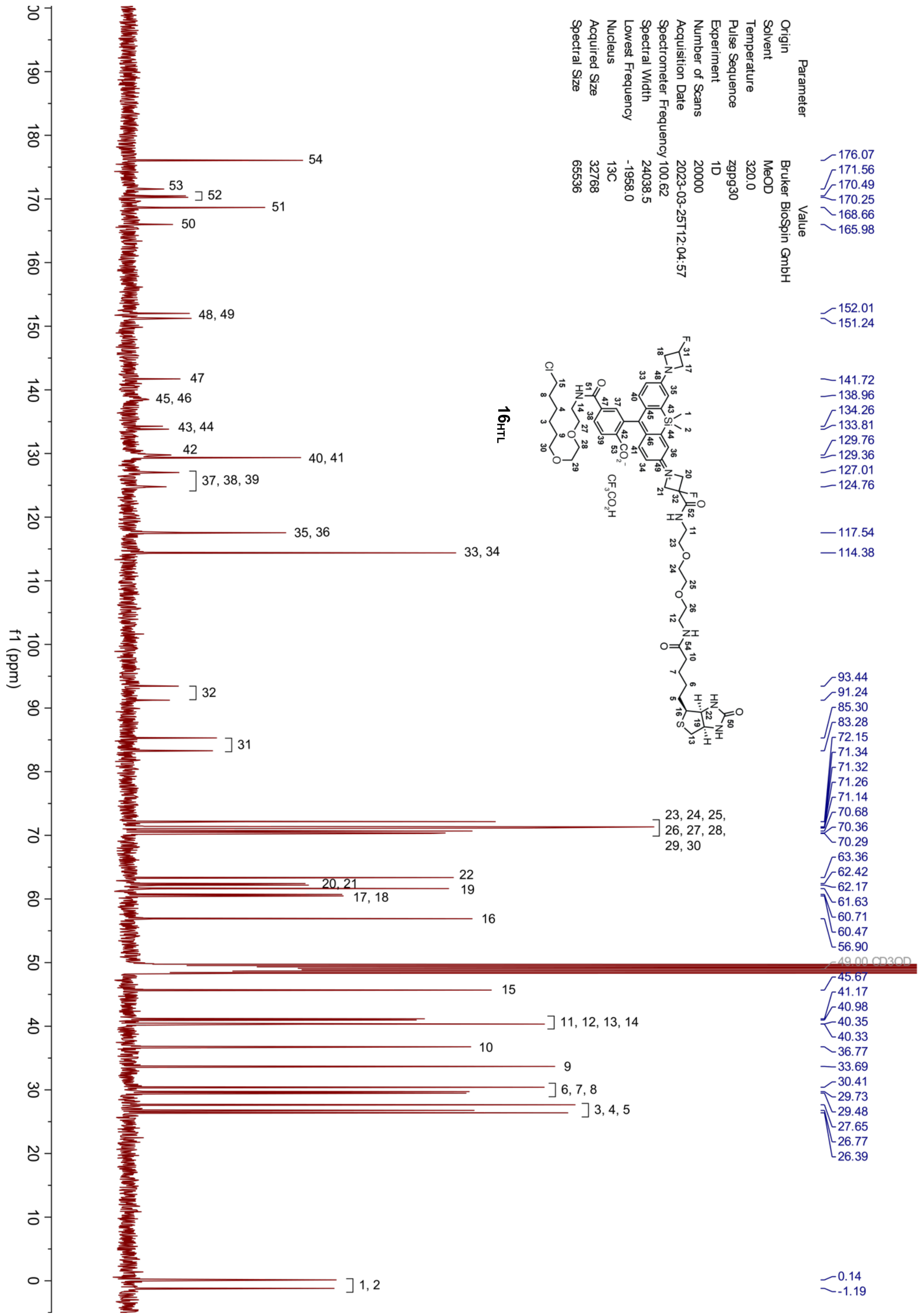

Parameter Value
Origin Bruker BioSpin GmbH
Solvent MeOD
Temperature 320.0
Pulse Sequence zgpg30
Experiment 1D
Number of Scans 20000
Acquisition Date 2023-03-25T12:04:57
Spectrometer Frequency 100.62
Spectral Width 24038.5
Lowest Frequency -1958.0
Nucleus 13C
Acquired Size 32768
Spectral Size 65536
16HTL
CF3CO2H
176.07
171.56
170.49
170.25
168.66
165.98
152.01
151.24
141.72
138.96
134.26
133.81
129.76
129.36
127.01
124.76
117.54
114.38
93.44
91.24
85.30
83.28
72.15
71.34
71.32
71.26
71.14
70.68
70.36
70.29
63.36
62.42
62.17
61.63
60.71
60.47
56.90
49.00 CD3OD
45.67
41.17
40.98
40.35
40.33
36.77
33.69
30.41
29.73
29.48
27.65
26.77
26.39
0.14
-1.19
54
53
52
51
50
48, 49
47
45, 46
43, 44
42
40, 41
37, 38, 39
35, 36
33, 34
32
31
23, 24, 25, 26, 27, 28, 29, 30
22
20, 21
19
17, 18
16
15
11, 12, 13, 14
10
9
6, 7, 8
3, 4, 5
1, 2
f1 (ppm)

| Parameter | Value |
|---|---|
| Origin | Bruker BioSpin GmbH |
| Solvent | MeOD |
| Temperature | 295.3 |
| Pulse Sequence | zg30 |
| Experiment | 1D |
| Number of Scans | 128 |
| Acquisition Date | 2021-07-05T12:18:00 |
| Spectrometer Frequency | 400.13 |
| Spectral Width | 8012.8 |
| Lowest Frequency | -1543.2 |
| Nucleus | 1H |
| Acquired Size | 32768 |
| Spectral Size | 65536 |

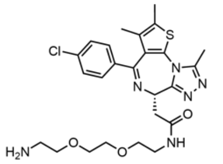

**S19**

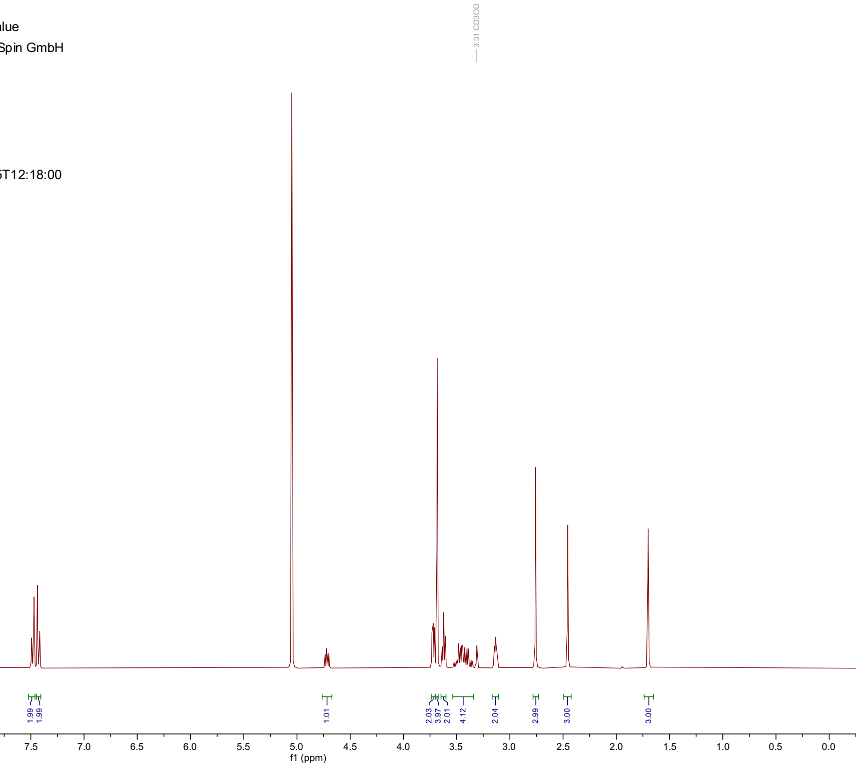


| Parameter | Value |
|---|---|
| Origin | Bruker BioSpin GmbH |
| Solvent | CDCl3 |
| Temperature | 296.1 |
| Pulse Sequence | zgpg30 |
| Experiment | 1D |
| Number of Scans | 1024 |
| Acquisition Date | 2023-06-20T12:28:32 |
| Spectrometer Frequency | 100.62 |
| Spectral Width | 24038.5 |
| Lowest Frequency | -2213.3 |
| Nucleus | 13C |
| Acquired Size | 32768 |
| Spectral Size | 65536 |

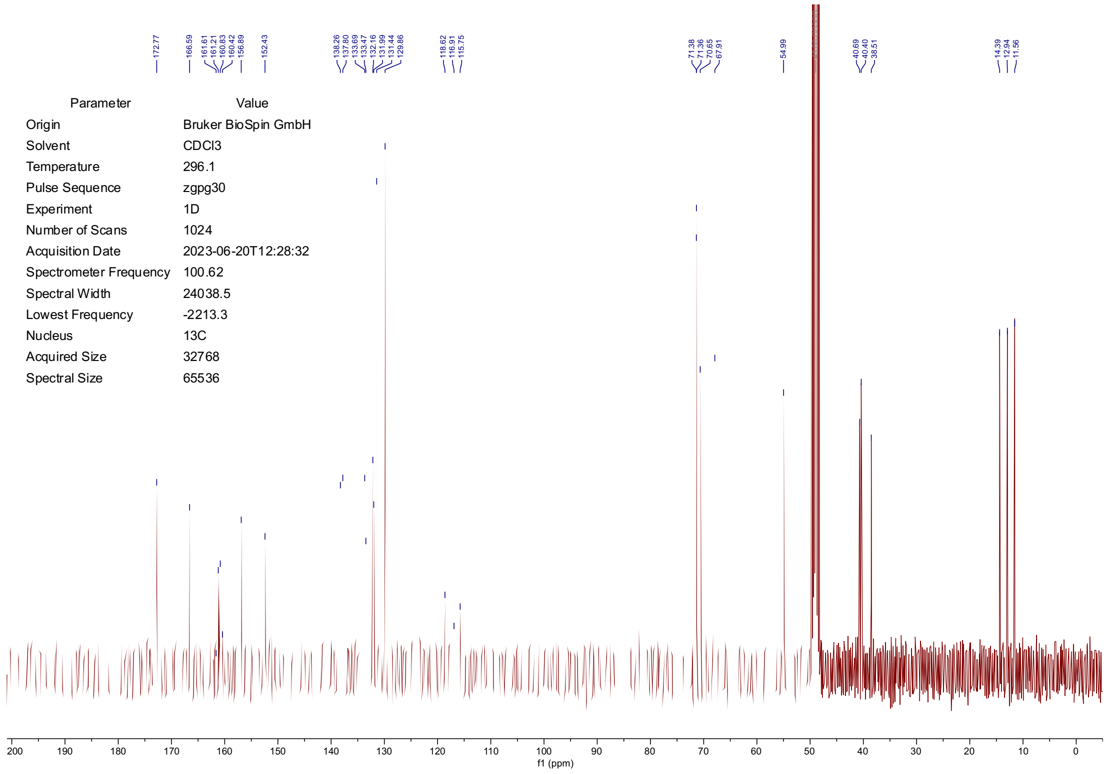

| Parameter | Value |
|---|---|
| Origin | Bruker BioSpin GmbH |
| Solvent | MeOD |
| Temperature | 295.4 |
| Pulse Sequence | zg30 |
| Experiment | 1D |
| Number of Scans | 128 |
| Acquisition Date | 2021-10-15T18:05:00 |
| Spectrometer Frequency | 400.13 |
| Spectral Width | 8012.8 |
| Lowest Frequency | -1543.0 |
| Nucleus | 1H |
| Acquired Size | 32768 |
| Spectral Size | 65536 |

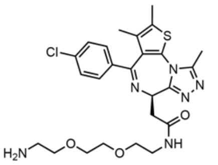


**S23**

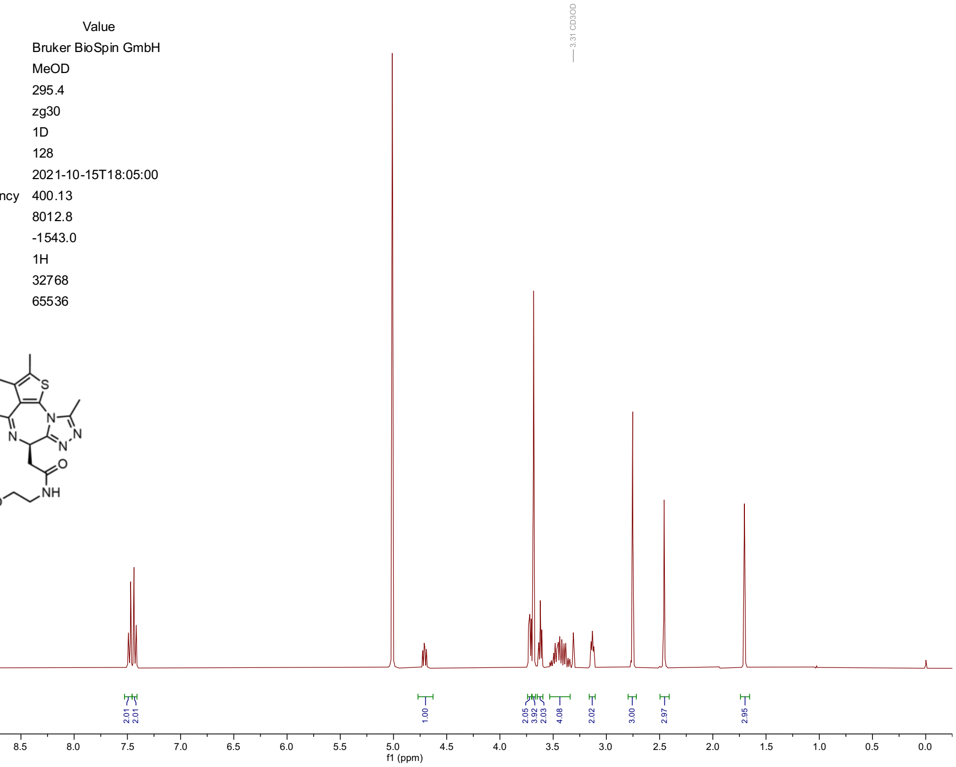


| Parameter | Value |
|---|---|
| Origin | Bruker BioSpin GmbH |
| Solvent | MeOD |
| Temperature | 296.2 |
| Pulse Sequence | zgpg30 |
| Experiment | 1D |
| Number of Scans | 16000 |
| Acquisition Date | 2021-10-16T10:05:00 |
| Spectrometer Frequency | 100.62 |
| Spectral Width | 24038.5 |
| Lowest Frequency | -1819.5 |
| Nucleus | 13C |
| Acquired Size | 32768 |
| Spectral Size | 65536 |

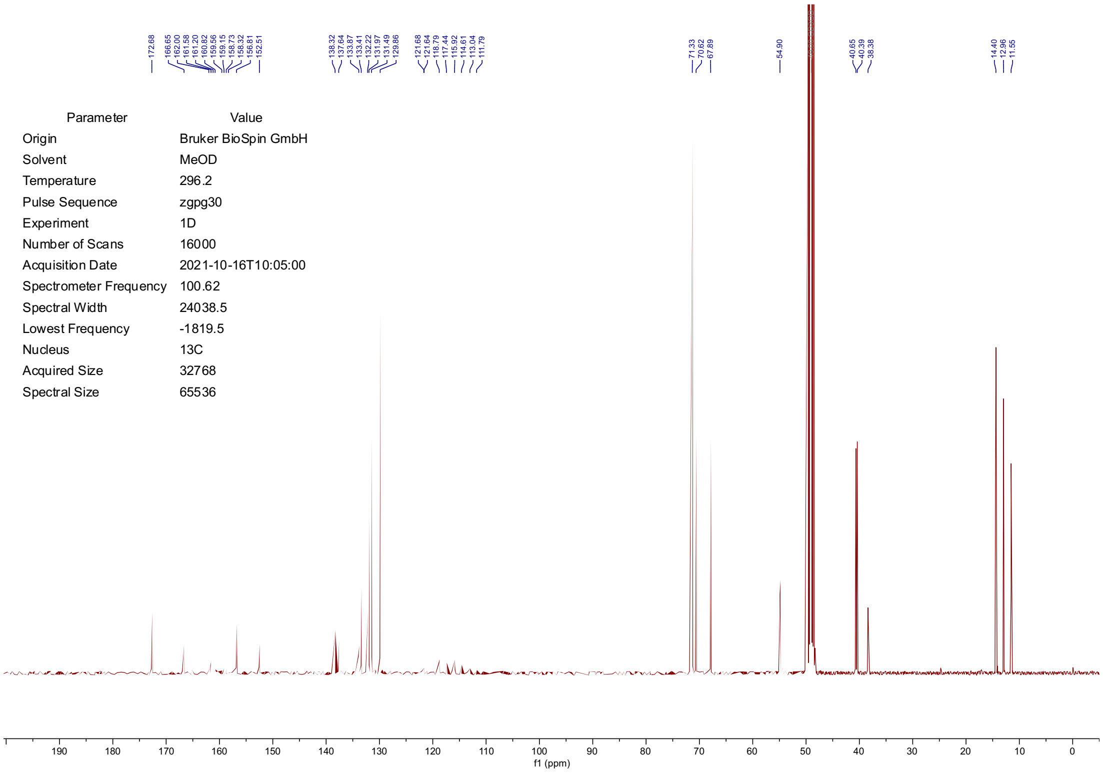

| Parameter | Value |
| --- | --- |
| Solvent | MeOD |
| Temperature | 298.0 |
| Pulse Sequence | zg30 |
| Experiment | 1D |
| Number of Scans | 48 |
| Acquisition Date | 2023-10-31T14:34:50 |
| Spectrometer Frequency | 400.13 |
| Spectral Width | 8012.8 |
| Lowest Frequency | -1543.3 |
| Nucleus | 1H |
| Acquired Size | 32768 |
| Spectral Size | 65536 |

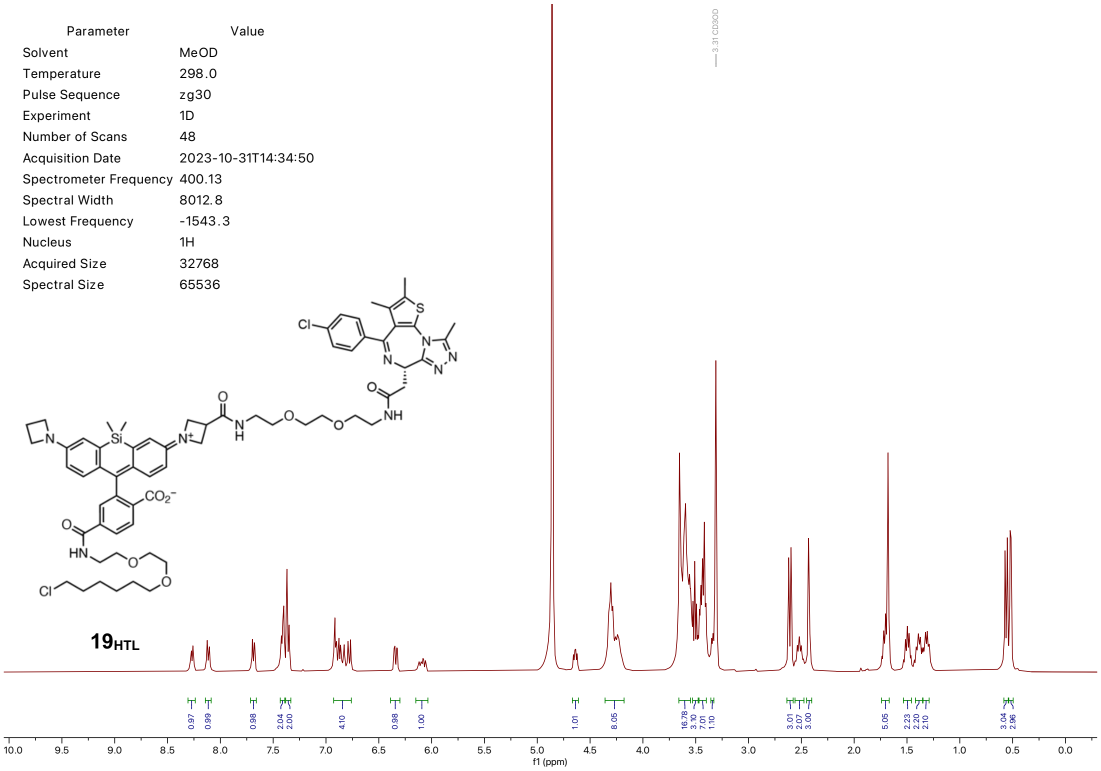


**19**$_{HTL}$

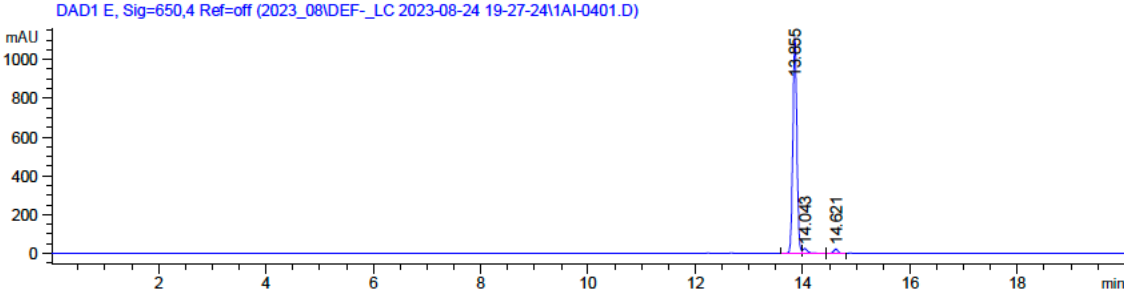


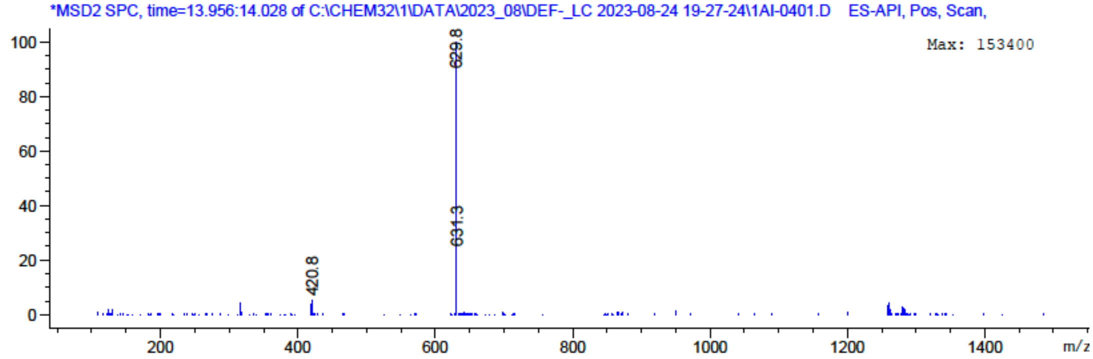

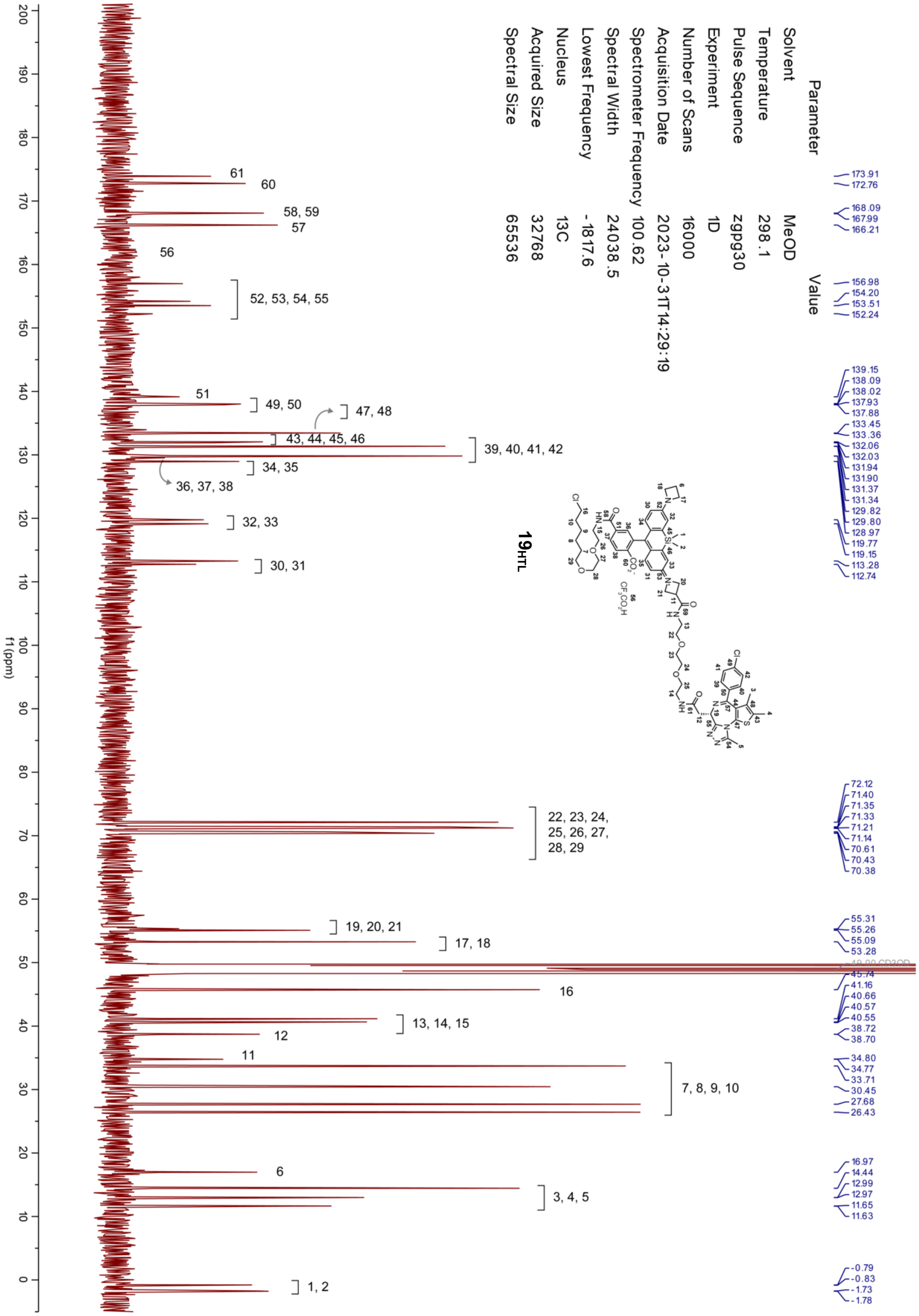

Parameter Value
Solvent MeOD
Temperature 298.1
Pulse Sequence zgpg30
Experiment 1D
Number of Scans 16000
Acquisition Date 2023-10-31T14:29:19
Spectrometer Frequency 100.62
Spectral Width 24038.5
Lowest Frequency -1817.6
Nucleus 13C
Acquired Size 32768
Spectral Size 65536
19HTL
173.91 172.76 168.09 167.99 166.21 156.98 154.20 153.51 152.24 139.15 138.09 138.02 137.93 137.88 133.45 133.36 132.06 132.03 131.94 131.90 131.37 131.34 129.82 129.80 128.97 119.77 119.15 113.28 112.74 72.12 71.40 71.35 71.33 71.21 71.14 70.61 70.43 70.38 55.31 55.26 55.09 53.28 45.74 41.16 40.66 40.57 40.55 38.72 38.70 34.80 34.77 33.71 30.45 27.68 26.43 16.97 14.44 12.99 12.97 11.65 11.63 -0.79 -0.83 -1.73 -1.78
61
60
58, 59
57
56
52, 53, 54, 55
51
49, 50
47, 48
43, 44, 45, 46
39, 40, 41, 42
34, 35
36, 37, 38
32, 33
30, 31
22, 23, 24, 25, 26, 27, 28, 29
19, 20, 21
17, 18
16
13, 14, 15
12
11
7, 8, 9, 10
6
3, 4, 5
1, 2
f1 (ppm)

| Parameter | Value |
| --- | --- |
| Origin | Bruker BioSpin GmbH |
| Solvent | MeOD |
| Temperature | 320.1 |
| Pulse Sequence | zg30 |
| Experiment | 1D |
| Number of Scans | 16 |
| Acquisition Date | 2023-06-23T22:13:36 |
| Spectrometer Frequency | 400.13 |
| Spectral Width | 8012.8 |
| Lowest Frequency | -1542.9 |
| Nucleus | 1H |
| Acquired Size | 32768 |
| Spectral Size | 65536 |

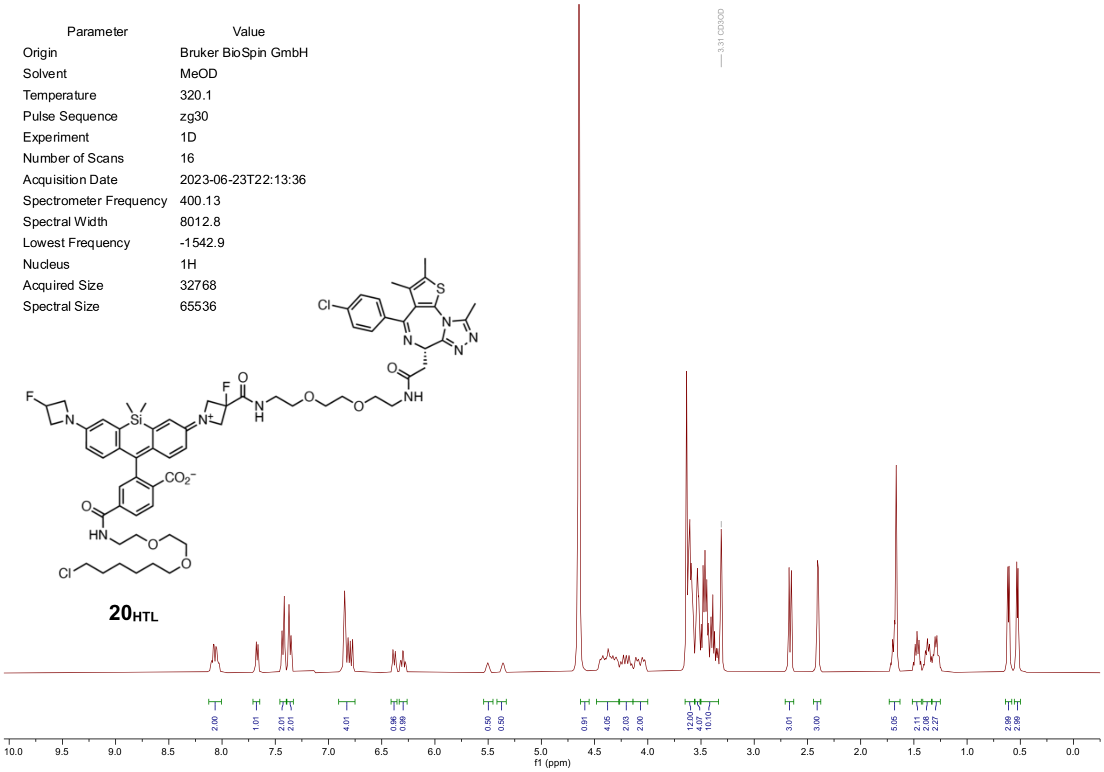


**20**$_{\mathbf{HTL}}$

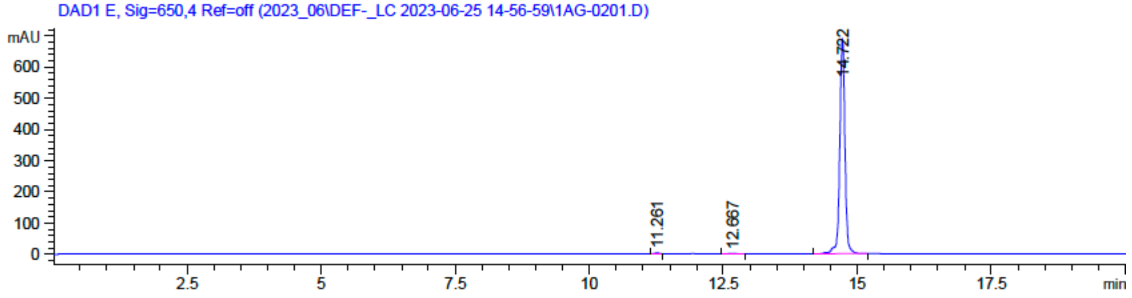


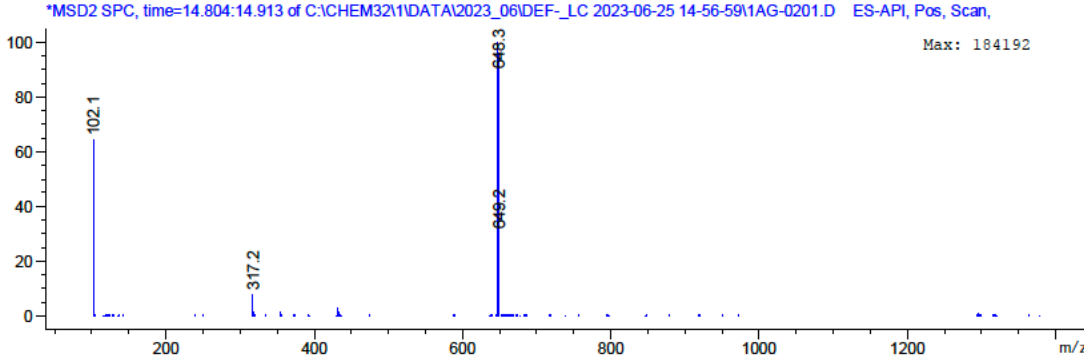

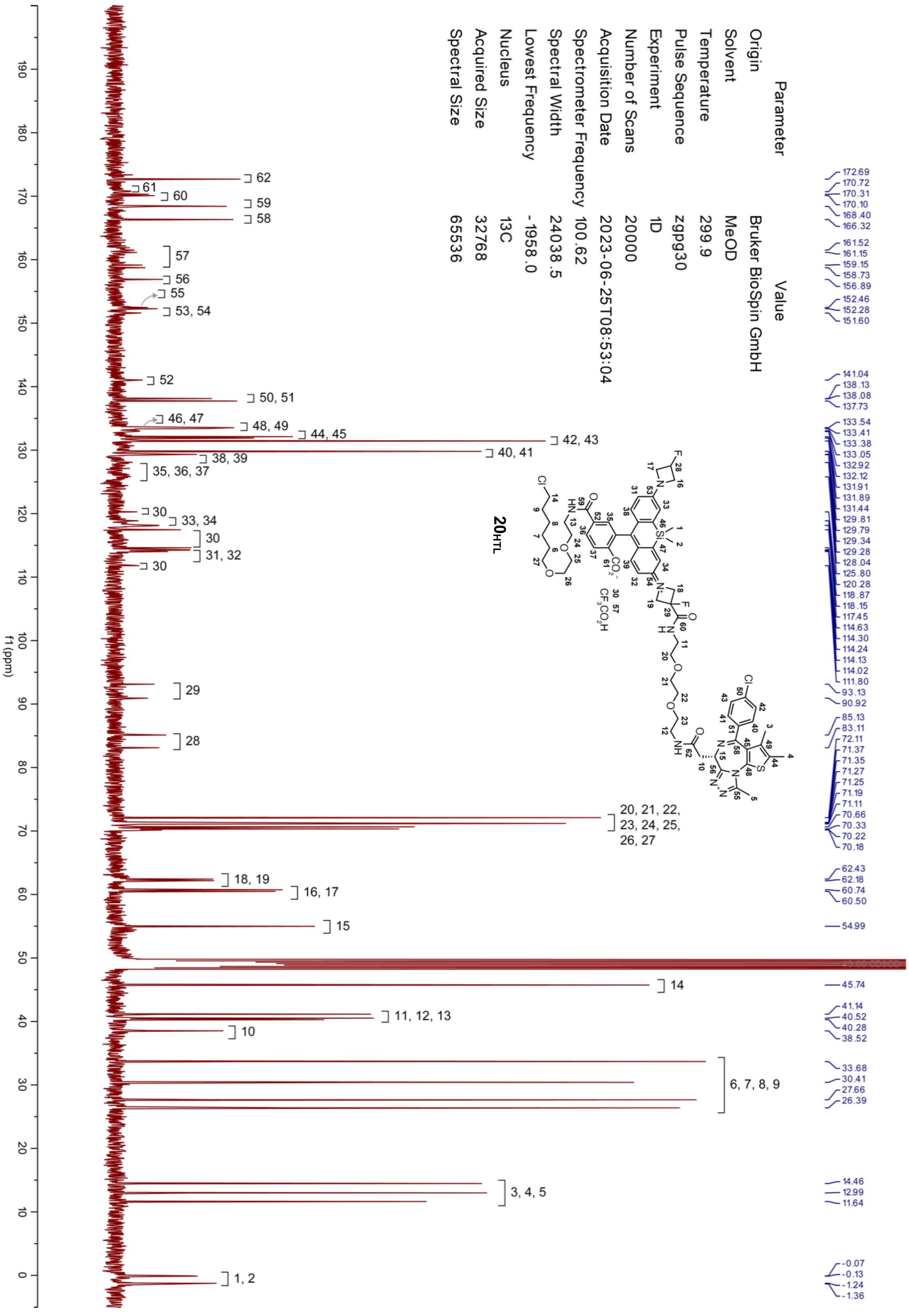
Parameter Value
Origin Bruker BioSpin GmbH
Solvent MeOD
Temperature 299.9
Pulse Sequence zgpg30
Experiment 1D
Number of Scans 20000
Acquisition Date 2023-06-25T08:53:04
Spectrometer Frequency 100.62
Spectral Width 24038.5
Lowest Frequency -1958.0
Nucleus 13C
Acquired Size 32768
Spectral Size 65536
20HTL
f1 (ppm)
172.69
170.72
170.31
170.10
168.40
166.32
161.52
161.15
159.15
158.73
156.89
152.46
152.28
151.60
141.04
138.13
138.08
137.73
133.54
133.41
133.38
133.05
132.92
132.12
131.91
131.89
131.44
129.81
129.79
129.34
129.28
128.04
125.80
120.28
118.87
118.15
117.45
114.63
114.30
114.24
114.13
114.02
111.80
93.13
90.92
85.13
83.11
72.11
71.37
71.35
71.27
71.25
71.19
71.11
70.66
70.33
70.22
70.18
62.43
62.18
60.74
60.50
54.99
45.74
41.14
40.52
40.28
38.52
33.68
30.41
27.66
26.39
14.46
12.99
11.64
-0.07
-0.13
-1.24
-1.36

| Parameter | Value |
|---|---|
| Origin | Bruker BioSpin GmbH |
| Solvent | MeOD |
| Temperature | 295.5 |
| Pulse Sequence | zg30 |
| Experiment | 1D |
| Number of Scans | 128 |
| Acquisition Date | 2021-10-19T10:23:00 |
| Spectrometer Frequency | 400.13 |
| Spectral Width | 8012.8 |
| Lowest Frequency | -1543.1 |
| Nucleus | 1H |
| Acquired Size | 32768 |
| Spectral Size | 65536 |

**$21_{HTL}$**

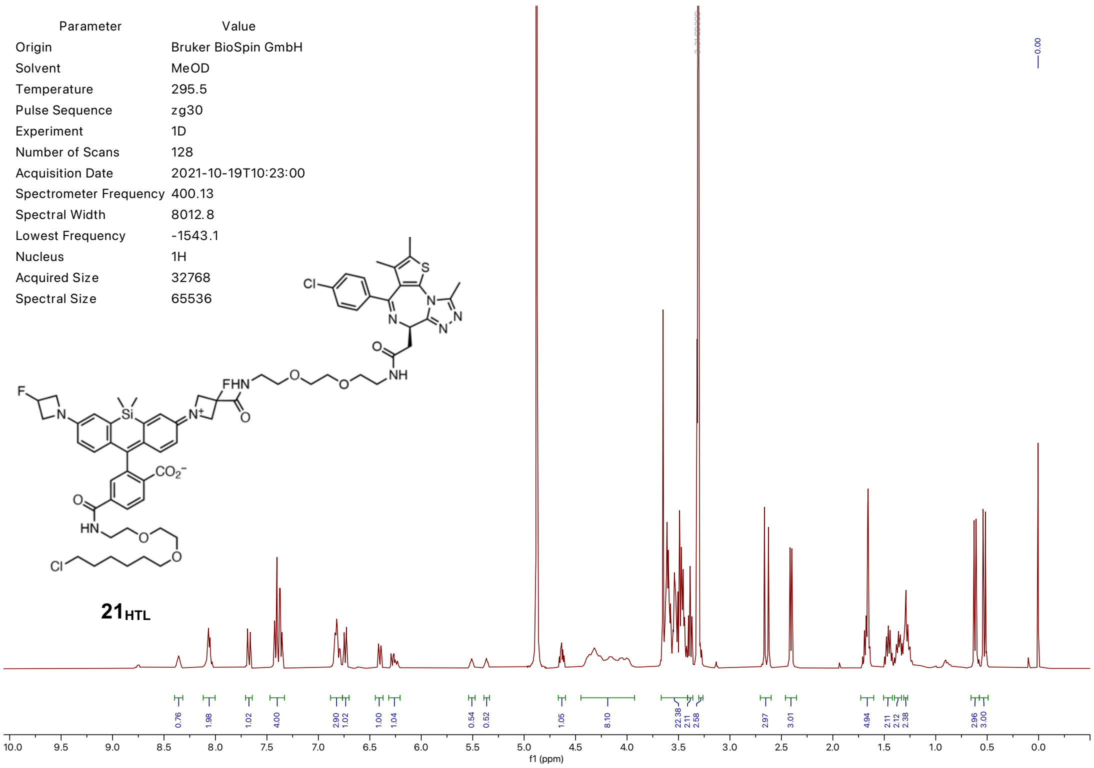


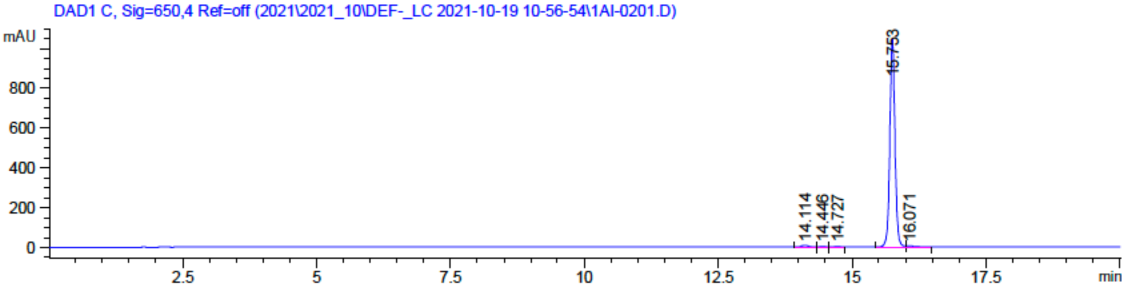


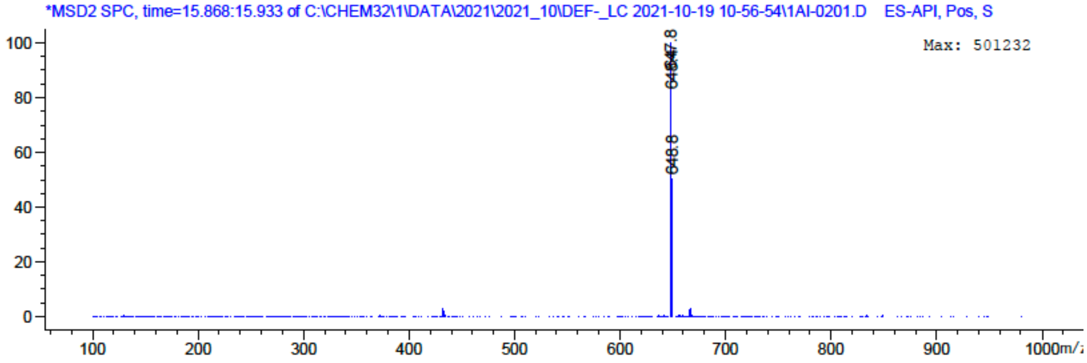
